# Nemo2.4: fast and accurate quantitative genetics forward-time simulations

**DOI:** 10.64898/2026.07.02.736177

**Authors:** Champak Beeravolu Reddy, Jobran Chebib, Olivier Cotto, Max Schmid, Frédéric Guillaume

## Abstract

We present Nemo 2.4, an advanced forward-time individual-based simulation framework designed to model the complex eco-evolutionary dynamics and genetic basis of quantitative traits. This tool addresses current challenges in evolutionary quantitative genetics by providing unprecedented flexibility and computational efficiency. Nemo 2.4’s modular architecture allows researchers to design custom life cycles by combining specialized Life Cycle Event (LCE) modules, from reproduction and dispersal to selection, crossing, and phenotype expression. The software supports diverse population models, including both Wright-Fisher (WF) and non-WF dynamics, spatially explicit models, and varying demography. Nemo 2.4 handles a wide range of genetic architectures, including both multi-allelic Quantitative Trait Loci (QTL) for general trait studies, and dense di-allelic Quantitative Trait Nucleotides (QTN) implemented with highly optimized bit-wise data structures. Crucially, it allows the simulation of QTNs on comprehensive genetic maps that incorporate other genetic elements, providing genomic-scale resolution. Key biological complexities are integrated natively: the model accommodates modular pleiotropy, dominance, and pairwise epistasis across multiple traits, facilitating the study of complex genotype-phenotype mappings. Furthermore, Nemo 2.4 models phenotypic plasticity through reaction norms and incorporates underlying liability thresholds, enabling the simulation of environmental influences on trait evolution with various forms of selection (e.g., Gaussian, linear, truncation). Due to its compiled design and memory-efficient data representations for large numbers of loci, Nemo provides a robust platform for running high-throughput simulations critical for testing theoretical predictions in polygenic adaptation and understanding evolutionary responses to changing environments.

## Introduction

We present here a new version of the individual-based, forward-in-time simulation software Nemo (Guillaume and Rougemont, 2006) with improved capacity to model the genetic basis and eco-evolutionary dynamics of quantitative traits. Nemo 2.4 meets the current advances and challenges in evolutionary quantitative genetics to investigate the eco-evolutionary dynamics of populations in changing environments.

Quantitative traits constitute the majority of traits studied by ecologists and evolutionary biologists, making it important to improve our capacity to model their genetic basis and better understand their eco-evolutionary dynamics. Examples of such traits cover most aspects of the morphology (*e.g.*, height, weight, colour), behaviour (*e.g.*, dispersal, cooperation) and life histories (*e.g.*, egg number, lifespan) of organisms. As such, quantitative traits play a fundamental role in how we conceptualize and investigate a species’ ability to adapt to its environment and evolve. They are similarly important for human health and agriculture, where diseases like Alzheimer, schizophrenia or diabetes in humans, or yield and growth traits in crops are known quantitative traits.

Understanding the genetic basis and evolution of quantitative traits in natural populations has long been a central goal of evolutionary biology. Quantitative genetics theory, building on the work of Fisher and Wright, provided a powerful analytical framework relating phenotypic evolution to the underlying genetic components under selection, drift, mutation, and gene flow (reviewed in Walsh and Lynch, 2018). However, these models mostly assume additive gene action, equilibrium conditions, and simple population structure, assumptions that enabled major theoretical advances, but which limit the range of questions that can be addressed analytically. In reality, the genetic architecture of complex traits involves non-additive effects: dominance, epistasis, and pleiotropy, which are widespread properties of quantitative trait loci (Mackay et al., 2009; Pickrell et al., 2016; Mackay and Anholt, 2024), and whose interactions shape trait evolvability in ways that additive models cannot fully capture (Hansen and Pelabon, 2021). Moreover, many pressing questions in evolutionary biology concern inherently non-equilibrium situations. Species responding to rapid environmental change, experiencing range shifts, or recovering from demographic bottlenecks are subject to complex feedbacks between demographic and evolutionary processes, called eco-evolutionary dynamics, where population growth, spatial structure, life history, and adaptive trait evolution are tightly coupled. In such settings, stochastic processes, particularly genetic drift and demographic stochasticity in small or fragmented populations, interact with selection and gene flow in ways that challenge analytical treatment. Forward-in-time, individual-based stochastic simulations address these limitations by explicitly tracking genotypes, phenotypes, and the demographic fates of populations through realistic life cycles, capturing the non-linear and stochastic dynamics that analytical models often simplify away. In practice, such simulations serve three broad purposes: testing and expanding theoretical predictions beyond the range of analytical models, generating realistic patterns of genetic variation as benchmarks for genomic inference methods, and building complex eco-evolutionary scenarios to evaluate hypotheses against empirical data.

Nemo was developed to meet these needs, proposing models that encompass the genomic to the geographic scales of genetic and environmental details. Nemo 2.4 accommodates models requiring different degrees of genetic realism, from few unlinked multi-allelic quantitative loci (QTL) to many di-allelic quantitative trait nucleotides (QTN) set on a genetic map, with flexible genotype-phenotype (G-P) maps incorporating dominance, epistasis, and pleiotropy. QTL-based traits efficiently model eco-evolutionary dynamics in large populations where fitness depends on the match between traits and local optima, as illustrated by studies of alpine species’ range dynamics under changing climatic conditions in existing populations (Cotto et al., 2017), the feedback of phenotypic plasticity on genetic divergence between populations (Schmid and Guillaume, 2017), or evolutionary rescue (Schmid et al., 2022). Models requiring finer genetic detail benefit from a large number of QTNs placed on chromosomal recombination maps, enabling ground-truth simulations where signals of selection can be dissected, for instance to assess how linkage disequilibrium among non-pleiotropic loci differently affects genetic correlations among traits relative to pleiotropic loci, and how genome-wide association analyses may not distinguish between the two (Chebib and Guillaume, 2021), or to test and develop novel statistical methods (Gilbert and Whitlock, 2015; Yeaman et al., 2018; Kemppainen and Guillaume, 2026).

The modelling of complex G-P maps requires a genetic-data representation that can accommodate non-additive allelic effects. The advent of genomics and genome-wide association studies (GWAS) called for simulators capable of bridging genomic-scale data representations with population-scale processes. Many widely used simulation platforms, such as sfs code (Hernandez, 2008), SLiM (Messer, 2013; Haller et al., 2026), and fwdpp (Thornton, 2014), implement approximations of the infinite site model where mutations appear at unique random positions on defined stretches of DNA sequences. In these implementations, only segregating mutations are tracked, making the approach memory-efficient and scalable to large genomes. We refer to this as a “sparse” genetic architecture because individuals carry information only at sites where mutations segregate. While efficient for population genetic modelling, the sparse approach complicates the modelling of quantitative trait architectures where allele identity must be tracked at many loci, whether polymorphic or not. This tracking is important when modelling non-additive allelic effects (dominance and epistasis) across large numbers of loci. Side-stepping the infinite site model, Nemo 2.4 implements a finite-site model where mutations occur recurrently at pre-defined loci with specified recombination rates, following early genetically-explicit implementations (e.g., simuPOP, Peng and Kimmel, 2005; Nemo, Guillaume and Rougemont, 2006; quantiNEMO, Neuenschwander et al., 2008). This “dense” representation explicitly stores allelic states at every locus in every individual, naturally supporting built-in models of locus-specific dominance, pairwise epistatic interactions among all loci, and variable pleiotropy where each locus can affect different subsets of traits. Both multi-allelic continuum-of-allele QTL and di-allelic QTN models are available, with a multivariate model of mutational co-variation enabling variation in the structure of the mutational and genetic variance-covariance matrices (Walsh and Lynch, 2018). Nemo 2.4 offers these as built-in generic models easily parameterized from few basic parameters, whereas the scripting of complex multivariate G-P maps can be cumbersome or not possible in sparse-architecture simulation platforms.

The quantitative trait model would be incomplete without accounting for environmental effects. Nemo 2.4 models phenotypic values that can be affected by both random environmental variation and deterministic environmental reaction norms. Phenotypic values can thus change randomly and plastically depending on the natal (developmental plasticity) or lifetime (labile plasticity) environments. In addition, in Nemo 2.4 the slope of a reaction norm can be tied to a quantitative trait and evolve, making plasticity an evolvable trait. These elements allow for in-depth modelling of quantitative traits with set heritability values, (mal)adaptive phenotypic plasticity, and genotype-environment interactions (GxE), as well as traits whose discrete phenotypes are determined by an underlying liability trait with a polygenic basis (Wright, 1934; Falconer, 1965). Liabilities and their evolving threshold traits effectively allow for the modelling of polyphenism and its evolution. Nemo 2.4 thus proposes the largest variety of quantitative trait architecture models available to date, and is an improvement over software derived from the original publication (Guillaume and Rougemont, 2006), including quantiNEMO (Neuenschwander et al., 2008, 2019) and Nemo-age (Cotto et al., 2020).

### Program architecture

The backbone of the program is the meta-population, a container of sub-populations (patches) containing individuals and their genetic elements. Nemo 2.4 then uniquely proposes a flexible and modular design organized around the *trait* and *life-cycle-event* (LCE) modules, that are optional and can be freely assembled. The *traits* encode different genotype-phenotype mappings, including quantitative trait loci, neutral loci, deleterious loci, and pairwise incompatibility loci. Each of them are encoded by different genetic elements that can be placed on a shared genetic map managing recombination among multiple chromosomes. LCEs are modifiers of the meta-population and encode all processes of a life cycle; mating and reproduction, dispersal, viability selection, age transition and mortality, density-dependent growth, and more (see the manual). The order of the LCEs is free to choose, with limited constraints, allowing for a wide variety of life-histories. In particular, Nemo’s life-cycle customization allows for both Wright-Fisher (WF) (Supplementary Example 1.1) and non-Wright-Fisher (non-WF) (Supplementary Example 1.2, 1.3) population models, spatially continuous populations with both gametic and zygotic dispersal, or alternative timing of key life-cycle events (*e.g.*, selection, mating, and migration). Additionally, Nemo features temporal LCE parameter values, which can then change during the course of a simulation. This ability is particularly useful to model shifting environmental conditions and to investigate evolutionary rescue (Schmid et al., 2022, Supplementary Example 1.2) or species range evolution (Gilbert et al., 2017; Cotto et al., 2017; Schmid et al., 2019, Supplementary Example 1.3).

A complete model requires to add the parameters specific to the selected *trait* and LCE modules, as well as those of the population and simulation components. The population component defines the number of patches (`patch_number`) and their carrying capacity (`patch_capacity`). The simulation component specifies the number of replicates (replicates) and the number of life cycle iterations (generations). Full examples, including parameter configurations, are provided in section *Examples* of the supplementary materials. More are available online.

#### A complete life cycle

The sequence of events in a life cycle is executed based on the user-defined order. Below is an example of a basic life cycle in a non-WF population, including reproduction, phenotypic expression, migration, and selection:

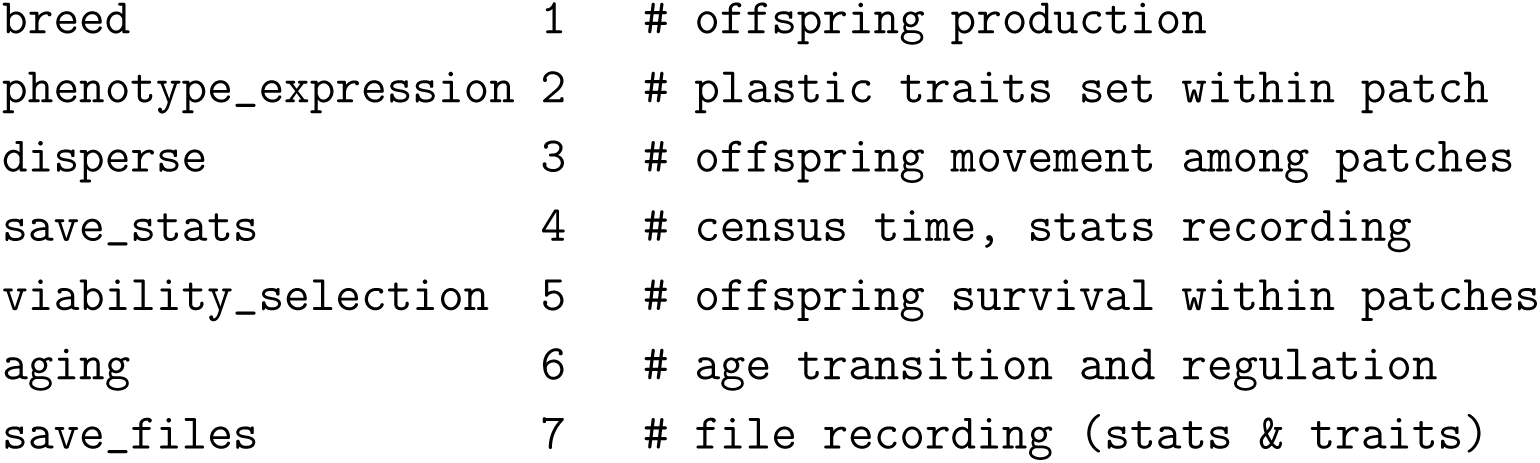

The timing of key events (e.g., phenotype expression, dispersal, and selection) can significantly impact model outcomes, especially when individual traits and fitness depend on within-patch environmental factors (e.g., trait optima or environmental cues). Adjusting the order of selection and dispersal allows for different scenarios: offspring can be selected in their natal patch (selection-dispersal) or their arrival patch (dispersal-selection). Similarly, phenotypic expression can be influenced by the environment in the natal patch (expression-dispersal) or in the arrival patch (dispersal-expression).

### Interface and I/O

The user interface is a simple text file containing parameters and values for chosen modules, which is passed as an argument to Nemo on the command line. To handle large parameter grid sets, we developed a mini language that allows encoding hundreds of simulations within single configuration files (see Supplementary Example 1.11). The resulting simulations can be distributed across CPUs, either locally or on HPC infrastructures. For job submission in large simulation campaigns, we developed nemosub, a helper program that uses the same configuration file as Nemo and automatically submits each simulation as an independent job allocation to computer clusters. In addition, Nemo natively supports running multiple replicates of the same simulation in parallel via its MPI layer (see Supplementary Examples, and online documentation).

In output, Nemo can write many component-specific files containing genotype and phenotype information in various formats (for quantitative traits: PLINK .ped file, SNP genotypes in 0/1/2 format), allelic values and frequencies, and summary statistics computed during a simulation run. Files can contain trait-specific information of sampled individuals or the whole meta-population. Nemo also exports and imports snapshots of the whole meta-population in binary format, which is useful as backups or to start a simulation from a burn-in (see Supplementary Examples 1.9-10).

### Implementation of the quantitative trait

The quantitative trait component in Nemo 2.4 implements two paradigms of the genetic architecture of quantitative traits: multi-allelic quantitative trait loci (QTL) and diallelic loci, which best represent quantitative trait nucleotides (QTN). The two models are toggled using a single parameter, `quanti_allele_model` with options continuous or diallelic (and `continuous_in_place, diallelic_asymmetrical` alternatives, see Supplementary Example 1.9, and Nemo’s manual).

#### QTL – the continuum-of-allele model

For studies focused on trait evolution and eco-evolutionary dynamics rather than detailed genomic data, Nemo efficiently models multi-allelic QTL using the continuum-of-allele model (Crow and Kimura, 1964). This model assumes that mutation can produce an infinite number of alleles at each locus. Allelic variation at a QTL is generated via a Gaussian mutational process, defined by its variance (effect size) and initial allelic values. The number of segregating alleles depends on the balance of evolutionary forces, with QTL becoming near-diallelic under strong selection or low mutation rates and highly multi-allelic under weak selection or high mutation rates (see e.g., Chebib and Guillaume, 2021). This implementation, grounded in uni- and multi-variate quantitative genetics theory, aligns with theoretical predictions for equilibrium variance (e.g., Lande, 1975; Turelli, 1984; Burger et al., 1989) and covariance (e.g., Lande, 1980).

We validated the simulation results obtained for continuous quantitative loci with theoretical expectations of the equilibrium additive genetic variance *V_a_* at mutation-selection-drift balance. The expectations are provided for two types of mutational regime, the Gaussian regime with “many mutations of small effect”, and the house-of-card (HoC) regime with “few mutations of large effect”. Figure 1 shows how well and how fast simulated populations reach the expected *V_a_* in the two mutational regimes. Almost all cases simulated reached their expectation within about 30,000 generations for a population of size *N* = 5000, with time to equilibrium increasing with the number of loci.

**Figure 1:**
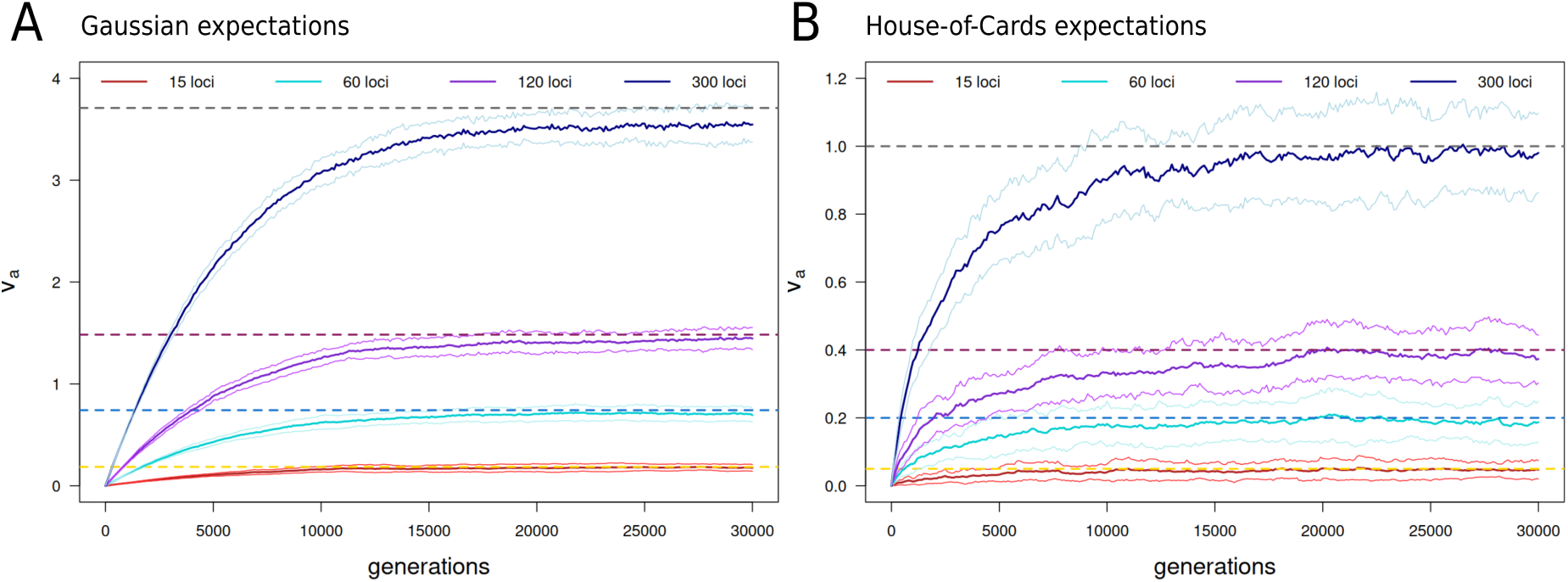
Model validation. – Dynamics of additive genetic variance (*V_a_*) at mutation-selection balance in a single population for one trait determined by a varying number of unlinked quantitative loci under the continuum-of-allele model. The dashed horizontal lines show the expected *V_a_* under the Gaussian (A) and House-of-Cards (B) approximation models (see Bürger, 2000; Walsh and Lynch, 2018). The solid lines are averages over 30 replicates. Smaller lines are one S.D. away from the average. **A**: Loci have mutation rate *µ* = 10*^−^*^3^ and mutational variance *α*^2^ = 10*^−^*^3^. The equilibrium variance is predicted as 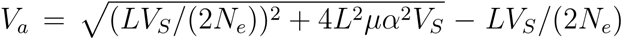 (Walsh and Lynch, 2018, p.1041). **B**: Loci have mutation rate *µ* = 10*^−^*^5^ and mutational variance *α*^2^ = 0.1. The equilibrium variance is predicted as 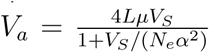 (Walsh and Lynch, 2018, p.1041). Population size is *N_e_* ≃ 5000 hermaphroditic individuals, simulated under the WF model for 30,000 generations. The initial population is monomorphic. Selection is stabilizing on trait optimum *θ* = 0 with strength *V_S_* = 100.

The following example shows the parametrization of four genetically independent traits with different mutational properties and number of non-pleiotropic loci (see also Supplementary Examples 1.1-6):

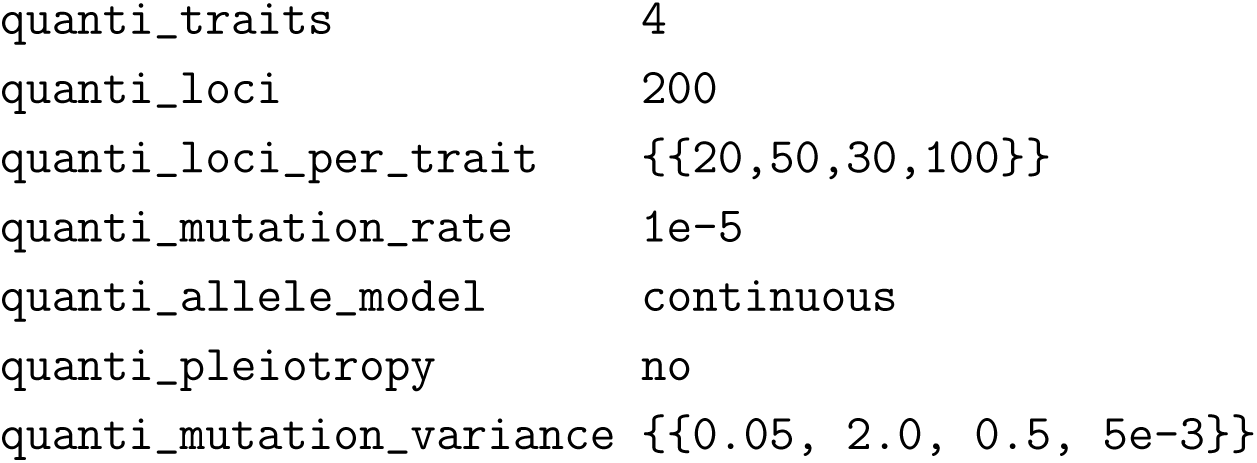

#### QTN – di-allelic model

Chromosome-scale dense genetic data can be more easily modelled with QTNs using the diallelic locus model. In this model, the allelic effects and map positions are specified for each locus. Mutations then land at pre-defined locations on the genome, with back mutations allowed (i.e., each mutation event flips between the two pre-defined allelic values). By default, the additive allelic effects *a_i_* are set symmetrically around zero as ±*a_i_* at each locus, but any two values can be passed. For instance, the `diallelic_asymmetrical` model encodes allele values {0*, b*}, with 0 as the default ancestral state, and *b* the derived state provided in input. The diallelic model allows for better monitoring of the effect of recurrent mutations at specific rather than random locations in the genome, easing the distinction between adaptive and non-adaptive genomic regions.

Practically, allelic values can be set randomly when starting a simulation by using a set of macros in the parameter file, as shown below for four traits each affected by 1000 loci (pleiotropy is off by default). Here, allelic effects are drawn from a Normal distribution with mean zero and standard deviation 0.1 (see also Supplementary Examples 1.7,9,10):

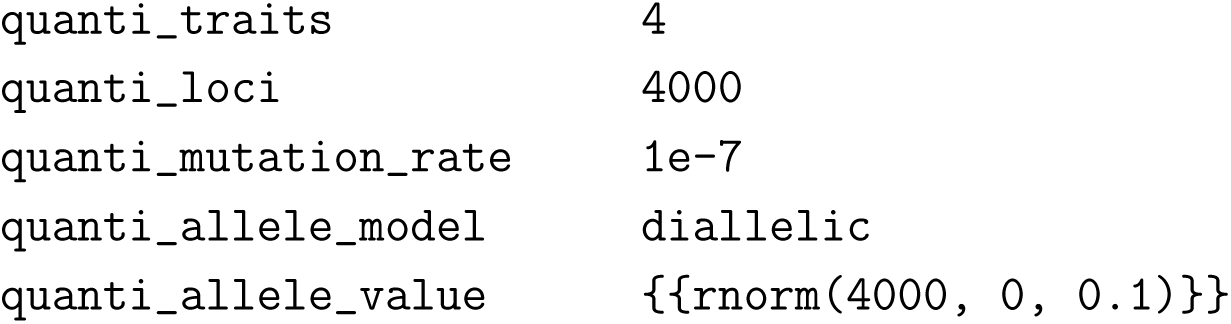

Since multiple equilibria exist for the genetic variance of di-allelic loci (see Bürger, 2000), we have not included simulation results to Figure 1. Figure S**??**, nevertheless, shows the dynamics and equilibrium value of *V_a_* for large numbers of di-allelic loci (10^4^ and 10^5^).

#### Multivariate phenotypes with pleiotropy

The number of phenotypes, or traits, encoded by the quantitative trait component is set with `quanti_traits`. Setting `quanti_traits` *>* 1 results in modelling genetically independent traits with distinct sets of loci affecting each trait unless a model of pleiotropy is chosen (parameter `quanti_pleiotropy`). Loci can either be fully pleiotropic (all loci affect all traits; option full to `quanti_pleiotropy`), or vary in which traits they affect (option variable), allowing for modular G-P maps. In that case, the pleiotropic degree of each locus is set with the pleiotropy matrix, which defines which locus affects which trait (see Figure 2). A pleiotropic locus carries one allelic value per trait (in haploid state) and these allelic values are affected by mutations with adjustable degree of correlation and variation. These characteristics are controlled by the mutational variance-covariance matrix—the M-matrix.

**Figure 2:**
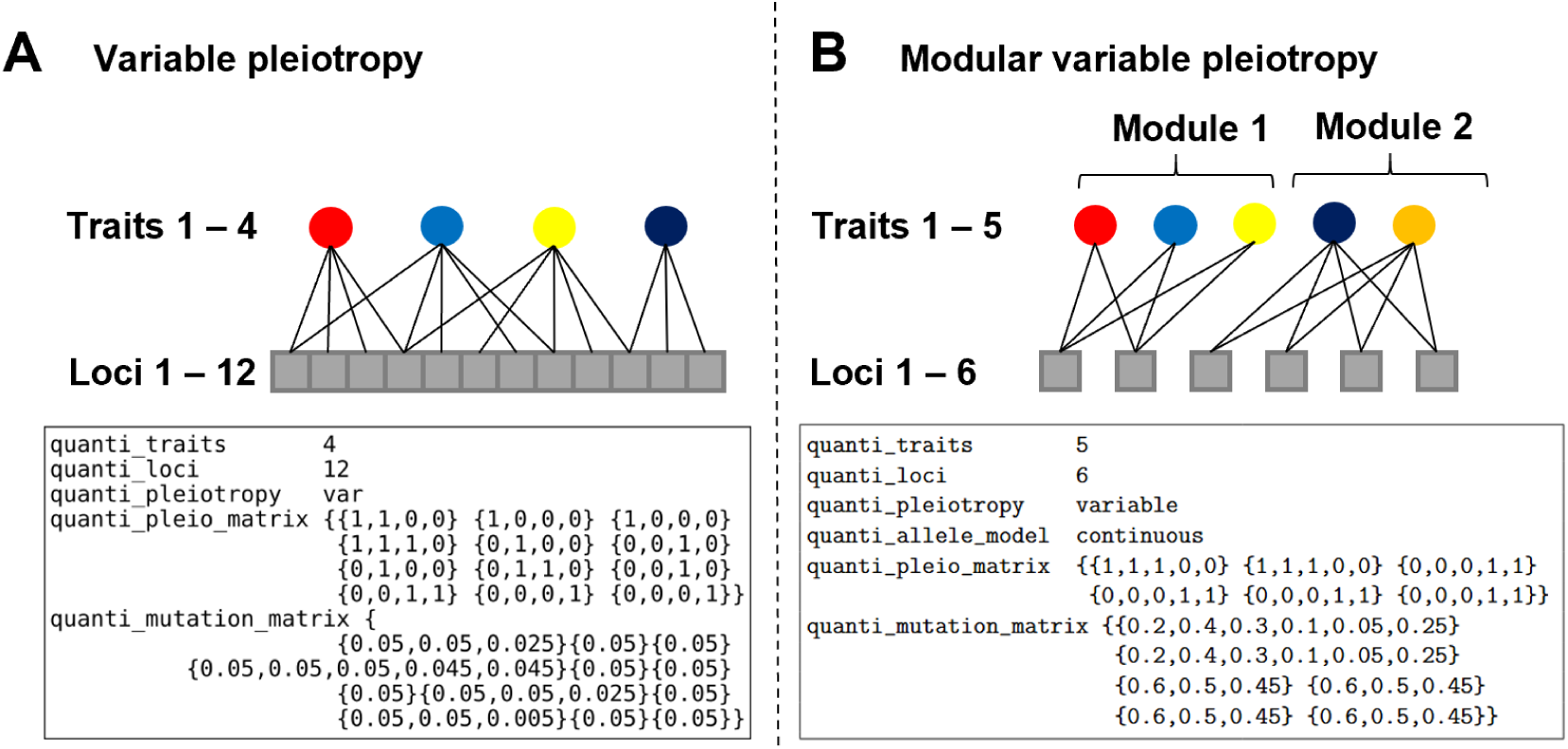
Two example parametrizations of genotype-phenotype maps with variable pleiotropy. **A**: Variable pleiotropy with 12 QTL (grey boxes) and 4 traits (colored circles). Loci either affect 1, 2 or 3 traits. The edges connecting loci to trait represent the pleiotropic connections of the genes set as 1 in the pleiotropy matrix (`quanti_pleio_matrix`), which has dimensionality loci×traits. The mutation properties at each locus are set with parameter `quanti_mutation_matrix`. That matrix holds the parameters of the locus-specific M-matrices with one M-matrix per row specified as {*var*_1_*, . . ., var_n_,* [*cov_i,j>i_, . . .*]}, with mutational variance for *n* traits and optional covariance for *n* ∗ (*n* − 1)*/*2 trait combinations. Loci affecting two traits thus receive mutational matrices with three parameters, two variances and one covariance. Non-pleiotropic loci only require the mutational variance. **B**: Modular variable pleiotropy with 6 QTL (grey boxes) and 5 traits (colored circles) split into two separate modules.

In the simplest case, with full pleiotropy, a single M-matrix is passed with per-trait mutational variance (diagonal) and pairwise covariance (off-diagonal). With variable pleiotropy, the M-matrix varies among loci when they vary in their pleiotropic degree (Figure 2). The mutational variance may differ among traits despite being affected by the same set of loci. In the following example, the four traits are affected by 200 fully pleiotropic loci with different variance and covariance, but same pairwise mutational correlation of 0.5 (set as covariance value, with 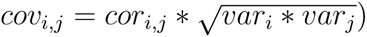:

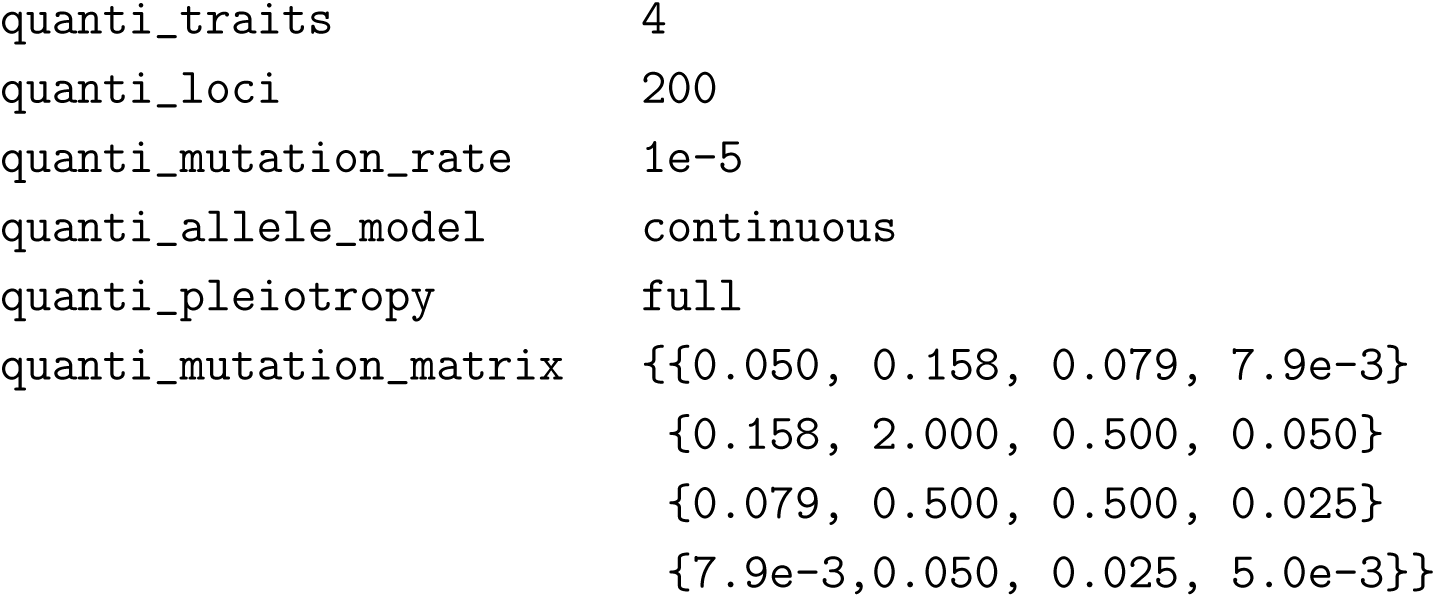

Nemo allows shortcuts to facilitate the parametrization of multi-dimensional G-P maps. For instance, in the above example, and since the correlation is the same for all traits, the M-matrix could be specified with two parameters:

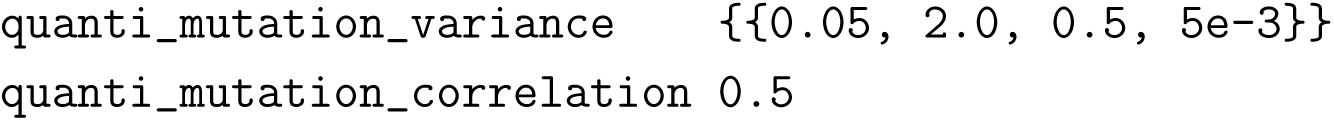

#### Non-additive effects – dominance and epistasis

In addition to the additive effects described above, the dominance and epistatic effects can be set explicitly with `quanti_dominance_effects` and `quanti_epistatic_effects`, respectively. For large numbers of loci, coefficients can be written to a separate file and linked to the parameters in the input file, as shown here:

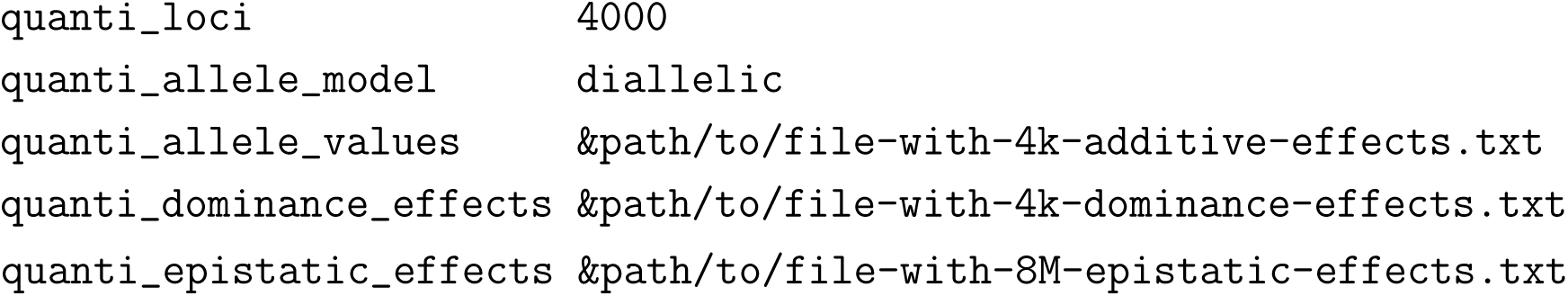

Dominance scales the genotypic value of a heterozygote locus *i* as *G_i_* = *a_i,_*_1_ + *a_i,_*_2_ + *k*|(*a_i,_*_2_ −*a_i,_*_1_)|, with *a_i,_*_1_ *< a_i,_*_2_, the two allele values at the locus. The dominance coefficient *k* is unbounded, and allows for under/over-dominance for |*k*| *>* 1, and additivity for*k* = 0. Epistasis modifies the genotypic value of a locus *i* to *G_i_* + Σ_*j*≠*i*_ *ɛ_ij_G_i_G_j_*, with *ɛ_ij_* the pairwise epistatic coefficient (Hansen and Wagner, 2001). Networks of interacting genes can be modelled as well, with parameter `quanti_epistatic_network`, thus enabling reduced number of interaction coefficients in input. Figure S1 shows how the epistatic model reproduces the theoretical expectation from Carter et al. (2005).

#### Recombination

Variation in recombination rates among loci and along chromosomes is ubiquitous in nature and a central feature of most modern forward-time IBS software. Nemo 2.4 implements a multi-trait recombination or genetic map on which all mappable genetic elements can be placed, including QTL and QTNs. Several options exist in Nemo to set the locus positions either randomly, regularly, or following existing empirical genetic maps. By default, no genetic map is built and loci are independent, with a recombination rate of 0.5. Alternatively, genetic maps can be composed of chromosomes of varying size where locus map positions are specified in centimorgan (cM), following the simple rule of 1cM = 1% recombination. The minimum recombination between two adjacent positions can be adjusted with the `quanti_genetic_map_resolution` parameter, as shown below:

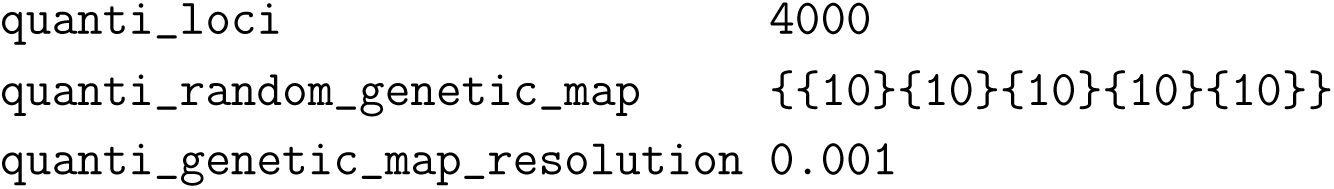

In that example, the 4000 quantitative loci are equally split among 5 chromosomes, or linkage groups, each 10cM long, set with `quanti_random_genetic_map`. Since the resolution is set to one-thousandth of a cM (with `quanti_genetic_map_resolution` 0.001), positions will be randomly drawn in interval [0, 10000) (instead of [0, 10)) and the re-combination rate between two adjacent positions will be 10*^−^*^5^ (i.e., base recombination rate for 1 cM distance: *r* = 10*^−^*^2^, × map resolution 10*^−^*^3^ gives *r* = 10*^−^*^5^). The locus map positions can be set specifically with `quanti_genetic_map`, for instance when based on an empirical genetic map. The positions are then usually saved in a separate file and chromosomes may differ in the number of loci they hold. Multiple traits, or genetic elements can be placed on the same genetic map with neutral, additive, or deleterious loci interleaved or even sharing the same map position (see Figure 3, and Supplementary Example 1.9, 1.11).

**Figure 3:**
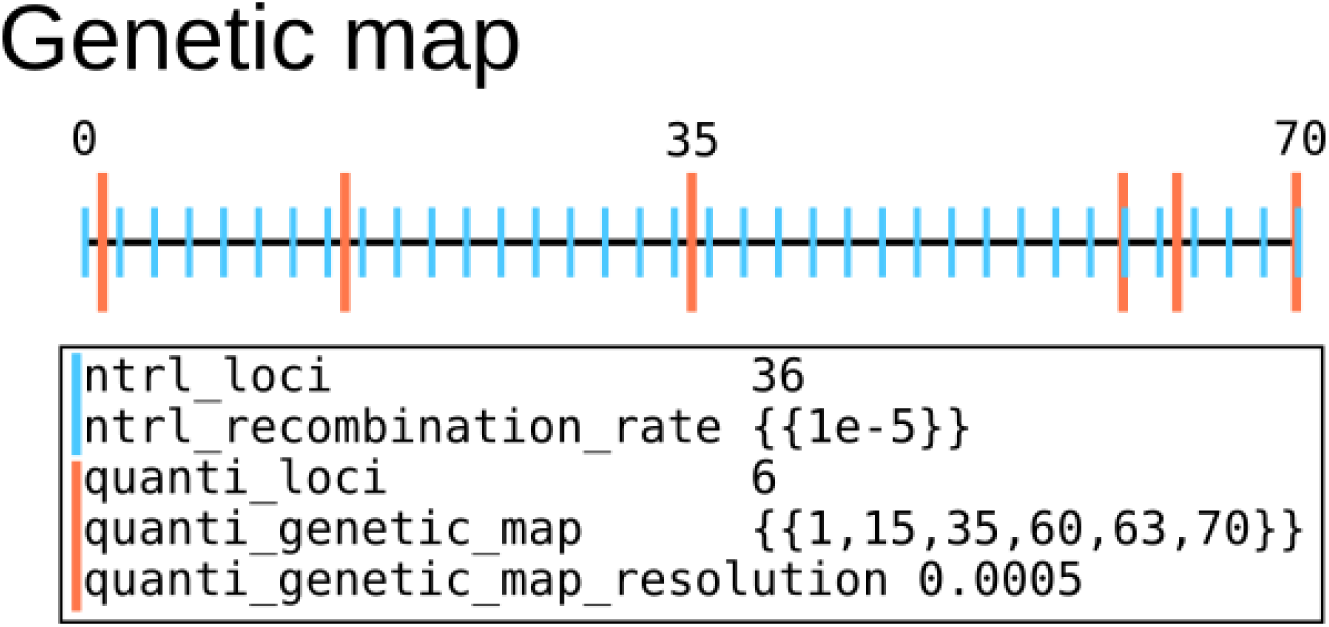
Example of a genetic map holding two types of loci: neutral (blue bars) and quantitative trait loci (red bars). The total map length is 35 × 10*^−^*^3^cM (remember that 1cM = 1% recombination). To place QTL loci between two neutral loci, the map resolution must be halved relative to the neutral trait to reach the minimum recombination rate of *r* = 5 × 10*^−^*^6^. This is achieved by setting `quanti_genetic_map_resolution` to 0.0005 cM. QTL at positions 60 and 70 are stacked at the same map positions as neutral SNPs making them physically linked with *r* = 0. Note that map positions are zero-based in Nemo.

#### Heritability

To simulate traits with heritability less than 1, environmental variance (*V_E_*) can be added to genotypic values when calculating phenotypes. This is achieved using the parameter `quanti_environmental_variance`, which specifies trait-specific *V_E_* values. Phenotypes are then computed by adding a normally distributed deviate with mean 0 and variance *V_E_* to the final genotypic value (including potential dominance and epistasis). The narrow-sense heritability is defined as *h*^2^ = *V_A_/V_P_* where *V_P_* = *V_A_*+*V_I_* +*V_E_* is the total phenotypic variance, *V_A_* is the additive genetic variance and *V_I_* is the interaction variance due to dominance and epistasis. During a simulation, the heritability *h*^2^ varies stochastically around its expected value. A specific *h*^2^ value can be reached by setting *V_E_* when *V_A_* is approximately known, such as after a burn-in simulation (note that if *V_I_* ≠ 0, then adding *V_E_* to the genotypic value changes broad-sense heritability, *H*^2^ = *V_G_/V_P_*)(see Supplementary Example 1.8).

Nemo 2.4 newly introduces a method to maintain constant *h*^2^ during simulations by adjusting *V_E_* as *V_A_* evolves. This approach works best for purely additive traits with precisely computed *V_A_*. For traits with non-additive effects (e.g., dominance or epistasis), broad-sense heritability (*H*^2^) can be set instead (Falconer and Mackay, 1996). Heritability parameters are controlled via the quanti modifier LCE (see Manual section 5.10). Maintaining constant heritability is useful when comparing crossing design scenarios or testing statistical methods with traits that have similar properties (e.g., Fraimout et al., 2024).

#### Phenotypic plasticity and liability

So far, we have considered a one-to-one mapping between traits and phenotypic values as configured with the quanti component. Nemo 2.4 also allows for more complex phenotypic models, either when a phenotype is encoded by multiple traits via a norm of reaction, or as a liability trait with discrete phenotypic values. In the norm-of-reaction (NoR) model, phenotypic plasticity is captured as a linear reaction norm: the phenotypic value *z* is set according to how the genotype *g*_0_ responds linearly to an environmental cue *e* as *z* = *g*_0_ + *e* · *g*_1_. The slope *g*_1_, representing the degree of phenotypic plasticity, can itself be linked to a quantitative trait and thus evolve in a simulation. Liability traits and phenotypic plasticity are governed by a dedicated LCE: phenotype expression, which sets the link to the quantitative trait(s) encoding the liabilities and the NoRs. Here is an example of two plastic phenotypes encoded by two NoRs affected by distinct environmental cues that vary between two patches. The first phenotype acts as the liability for quantitative trait 1 (see complete example with a moving trait optimum in Supplementary Example 1.6):

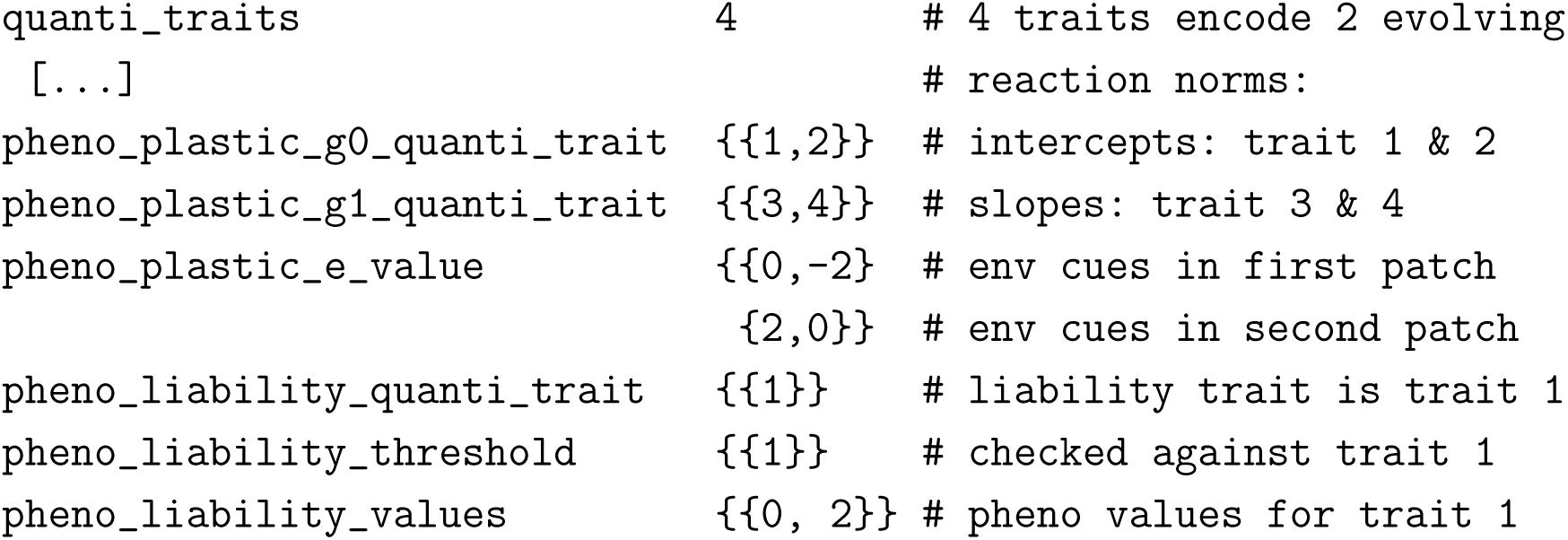

Because phenotypic plasticity can be linked to multiple quantitative traits, the NoR intercept and slope each can be controlled by distinct genetic loci (with varying levels of genetic linkage, epistasis and/or dominance among them), or by a set of pleiotropic QTLs controlling *g*_0_ and *g*_1_ (with varying levels of mutational correlation). The same applies to the liability traits.

#### Selection

Nemo 2.4 implements several selection functions acting on quantitative traits. The most commonly used is the multivariate Gaussian function, which allows for correlated and uncorrelated selection on multiple phenotypes. Quadratic, linear, and truncation selection can also be utilized for non-correlated selection. Gaussian and quadratic functions define an optimum trait value *θ* at which fitness is maximized. That optimum value is patch-specific and can change over time (see Supplementary Examples 1.2-6). These features enable modelling of spatially heterogeneous environments with shifting conditions. The trait optimum value can be a proxy for environmentally determined selection and provides a tool to model eco-evolutionary scenarios of adaptation to environmental variation (e.g., Cotto et al., 2017; Gilbert et al., 2017). The Gaussian model has two main parameters, the vector of trait optimum values *θ* and the selection variance-covariance matrix Ω describing the width and orientation of the fitness surface in phenotypic space around *θ* (see Manual, section 5.7.3).

#### Crossing design

Quantitative genetics seeks to dissect phenotypic variation into its genetic and environmental components to understand trait inheritance and predict their responses to selection. The methodology relies on controlled crosses in a common environment to produce individuals with known genetic relationships (Falconer and Mackay, 1996). Nemo 2.4 implements the cross LCE to model simple crossing designs with half-sib/full-sib crosses within patches (see Supplementary Example 1.7), as well as between-patch hybrids to estimate heterosis. A cross followed by dispersal and selection can simulate a reciprocal transplant to assess local adaptation. The cross LCE also accepts a pedigree file as input information, enhancing the toolkit available to quantitative geneticists to model trait inheritance on known pedigrees from monitored natural populations or to encode more complex breeding designs than implemented with the set of basic parameters in the cross LCE (see Supplementary Example 1.8). Simulating the inheritance of quantitative traits, and other genetic elements, on a known pedigree can help assess the accuracy of statistical methods (see e.g., Nietlisbach et al., 2017).

#### Performances

In figure 4, we show the CPU time and memory consumptions of simulations run for a set of populations sizes and number of QTL and QTN with varying recombination and mutation rates. The results show that computing time is dominated by population size and number of loci and, to a smaller extent, by recombination rate. Mutation rate only marginally affects computing time (Fig. 4a). In general, QTL simulations with a product of population size (*N*) and number of loci no larger than 10^6^ are executed in less than a minute for 1000 iterations (generations). For instance, the largest simulations in Figure 1 with *N* = 5000 and 300 QTL took about 15 minutes to complete, or around 30 seconds per 1000 generations. It is also clear from figure 4d that QTN simulations are faster when population size *N >* 10, with 4-6× and *>* 10× speedups for 10^3^ *< N* × *L <* 10 and *N* × *L* ≥ 10, respectively. QTNs are encoded on bit strings, with one di-allelic locus occupying a single bit in the bit sequence. This allows for large reductions of memory consumption (Fig. 4c) and execution time thanks to optimised bitwise operations. Such operations help when counting bits within bit strings and executing logical bit-wise instructions on large numbers of loci at once. The speedups can be seen in Figure 4b, particularly when QTN have equal allelic effects (marked as *equal* in Fig. 4b) relative to when all QTN have different effects, drawn from a distribution (marked as *distr* in Fig. 4b-d). Our benchmarking revealed that the reduction of execution time in the *equal* setting is marked starting at *L* ≥ 1000 with ∼ 2.7× speedup, reaching 12× for *L* = 10^4^ relative to the *distr* setting (*>* 100× relative to QTL, see table S1). In general, the QTN bitstring architecture is very parsimonious, allowing simulations to run under one minute per 1000 generations in all cases with equal allelic effects, and overpassing it for *N* × *L >* 10^7^ with unequal effects. CPU time consumption tends to converge for large *N* × *L* values irrespective of the population size, for both QTL and QTN-*distr* settings. Execution time strongly increases at high recombination rates, for instance when reaching *r* = 0.01 (see Fig. S3). Free recombination (*r* = 0.5) is punishing when simulating large numbers of QTN. A byte architecture fallback option exists for QTNs when modelling of freely recombining loci is necessary. In contrast, memory consumption is a near-linear function of *N* × *L*, independent of mutation rate and recombination rate. Overall, Figure 4c shows that a typical simulation of a chromosome segment bearing 10^5^ QTN would use less than one GB of RAM in populations of up to 10000 individuals, whether QTNs are polymorphic or not.

**Figure 4:**
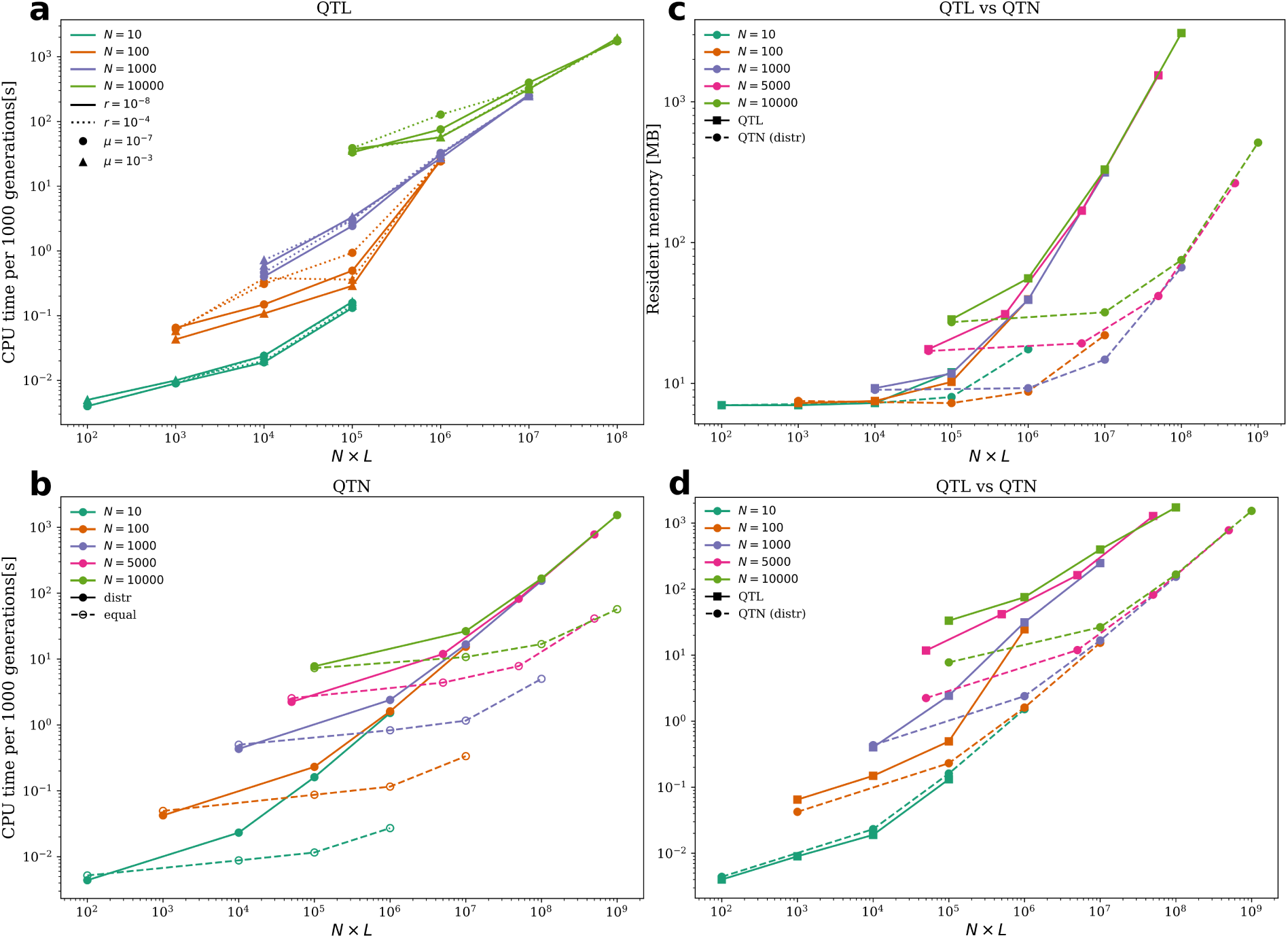
**Benchmarking** of simulations in Wright-Fisher populations of size between *N* = 10 to *N* = 10000 and for a number of loci *L* = {10, 100, 1000, 10000} for QTL, and *L* = {10, 1000, 10000, 1000000} for QTNs. Points represent averages of 10 replicates run for 1000 generations for QTL, and 5 replicates run for 5000 generations for QTN. Panels **(a,b,d)** show variation in computing time (CPU *user* ’s time) per 1000 generations in seconds (log-scale, y-axis) as function of *N* × *L* (log-scale, x-axis). Panel **(c)** shows the *peak resident memory* of the simulations in MB (mega bytes, log-scale, y-axis). Panel **(a)** shows execution time of the QTL simulations for two mutation rates (*µ* = {10*^−^*^7^, 10*^−^*^3^}) and two recombination rates (*r* = {10*^−^*^8^, 10*^−^*^4^}). Panel **(b)** shows CPU time of the QTN architecture (bitstring) for *µ* = 10*^−^*^7^ and *r* = 10*^−^*^8^, and for equal additive allelic effect across loci (*equal*, dashed lines) against the unequal effects drawn from a distribution (*distr*, solid lines). Panels **(c,d)** compare memory and CPU time consumption between QTL and QTN simulations for *µ* = 10*^−^*^7^ and *r* = 10*^−^*^8^, and same patch sizes. Only QTN simulations reached *N* × *L* = 10^9^. Results are presented for QTN with un-equal allele effects (dashed lines). Simulations were performed on a AMD Ryzen threadripper pro 3955wx processor and statistics were recorded with GNU time tool. The C++ code was compiled with GNU gcc (v11.4) with compilation flags: -O3 -march=native.

#### Comparison with other software

Nemo and quantiNemo are the two main IBM software proposing native implementations of quantitative traits. Yet, they differ markedly in their allelic models. Whilst Nemo proposes di-allelic and continuous models, quantiNemo proposes a discrete K-allele model where up to 256 allelic values per locus can be specified. It then approximates the continuum-of-allele model by discretizing normally distributed allelic values over the number of alleles chosen with a default span of ±20*α* (*α*^2^ = mutational variance). While this procedure doesn’t fully capture the tails of the normal distribution, it provides good approximation of the trait dynamics with 256 alleles (Fig. S6). A direct comparison is also possible with SLiM, one of the most widely used forward-time IBMs. Because quantitative traits are not built-in in SLiM, a fixed QTL or di-allelic QTN panel must be emulated in script, and as each locus is carried by individually tracked mutations, both memory use and runtime grow with the number of segregating loci. To quantify this, we benchmarked the three programs on an identical Gaussian stabilizing-selection model in one WF population. The three models provide identical equilibrium results for mean phenotype and additive genetic variance for both 100 unlinked QTL and 1000 di-allelic QTNs linked on a genetic map with five chromosomes (Fig. S6). They, however, perform very differently (Fig. 5), with Nemo being the fastest in all configurations and leanest in most cases. SLiM, being a “sparse” data simulator, is particularly less efficient at modelling fix sets of di-allelic QTNs, where its memory use and runtime are far beyond Nemo’s (Fig. 5B, S5). This is the very regime in which Nemo’s dense, bit-wise QTN model is most efficient. quantiNemo does worst memory-wise than the other software, especially for linked QTNs, for unclear reasons (with ∼4.5 GB RAM for *N* = 10000, *L* = 5000, vs. ∼51 MB for Nemo and ∼679 MB for SLiM, see Fig. S5, note that Nemo uses least memory even with byte-wise QTN data type). Since SLiM stores mutation effect in a single object per mutation, its memory consumption for QTL is less than Nemo’s, which stores floating-point allele values in each individual (Fig. 5A, S4).

**Figure 5:**
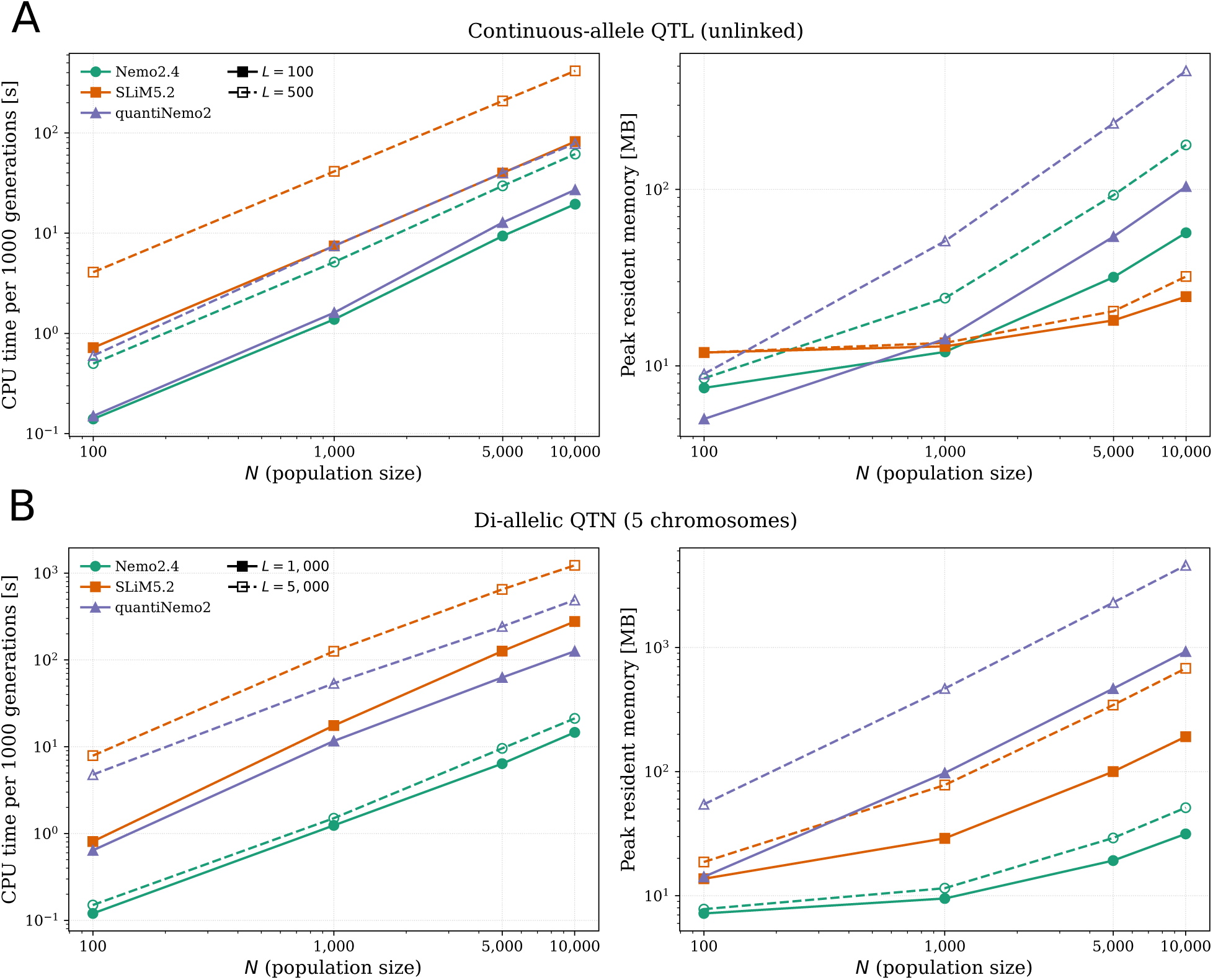
**Software performance comparison** between Nemo2.4.1, quantiNemo2.0.2, and Slim5.2, showing mean CPU time per 1000 generations (ticks) and mean resident memory (Max RSS) in MB of RAM for 5 replicates of a model with one WF population under Gaussian stabilising selection on one quantitative trait with *V*_S_ = 10. Population size varies as *N* = {100, 1000, 5000, 10000}. **A** – Trait encoded by *L* = {100, 500} unlinked (*r* = 0.5) QTL under the continuum-of-allele model, or an approximation of it. Loci had mutation rate *µ* = 1*e^−^*^4^ and mutation variance *α*^2^ = 0.05. **B** – Trait encoded by *L* = {1000, 5000} QTN linked on 5 chromosomes with *r* = 1*e^−^*^5^ on each, with *µ* = 1*e^−^*^5^ and additive effect ±0.01. Both models had environmental variance *V*_E_ = 1 added to the genotype. Results for a larger set of parameter values are in Supplementary Figures S4 and S5. Simulations were performed under same conditions as in Figure 4.

## Discussion

This new version of Nemo provides highly optimized, native implementations of complex genetic architectures and complex life cycles, thereby meeting the current and future needs of evolutionary quantitative genetics. The software’s rigorous validation against theoretical expectations, its memory-efficient bit-wise data structures, and its modular Life Cycle Event system make it a robust instrument for high-performance simulations of polygenic adaptation. Specifically, the native implementation of the continuum-of-alleles and di-allelic models, with matrices for variable pleiotropy, multi-trait dominance, and built-in Hansen–Wagner multilinear epistasis, makes this framework particularly powerful.

Nemo 2.4’s genetic data architecture permits two main domains of application, each with a distinct data structure. The QTL continuum-of-allele model is best suited when trait dynamics are more important than fine-grained genomic data generation. A typical model based on this architecture would implement 10 to 100 unlinked QTL with various degree of pleiotropy to model eco-evolutionary dynamics in complex demographic and spatial settings, for instance to understand species’ range shifts under climate change in multiple phenotypic dimensions (Cotto et al., 2017) or with the evolution of reaction norms (Schmid et al., 2019). The continuous model imposes a high memory cost for large numbers of QTL (*L >* 1000) and limits its usefulness in a genomic context. The second domain of application relies on the bit-wise genetic data architecture and applies best to the modelling of dense genetic maps with reduced recombination at the chromosome level. It allows for the simulation of very large numbers of di-allelic genetic markers (QTN) to generate genomic-scale data with empirically estimated recombination rate variation along chromosomes. The di-allelic model is thus ideal to investigate genomic signals of selection on polygenic traits when coupled with densely distributed neutral markers (Kemppainen and Guillaume, 2026). At the phenotypic level, QTL-based traits have no imposed limit to their possible values, since the continuum-of-allele model allows accruing allelic values over evolutionary time, unless specified otherwise (i.e., with allele model `continous_in_place`). In contrast, QTN-based traits have clearly defined phenotypic boundaries set by the sum of allele effects.

Nemo’s data architecture achieves high throughput for quantitative traits because it naturally accommodates the dense genetic architectures typical of polygenic adaptation, where thousands of alleles fluctuate at intermediate frequencies. As described above, the sparse approach used by other simulators tracks only segregating mutations, which is efficient when few variants segregate but slows down computation as their number increases (Matthey-Doret, 2021, see also SLiM benchmarks in Figures 5 and S5). The sparse approach additionally imposes a heavy computational load when calculating genotypic or phenotypic values across thousands of mutations, unless specific optimizations are implemented. This computational cost increases further when alleles at specific loci have dominance or epistatic effects, particularly when phenotype calculations rely on scripted rather than compiled code. The dense approach implemented in Nemo alleviates these hurdles. In particular, Nemo 2.4 introduces a significant performance optimisation by encoding allelic states in a bit-wise data structure, also applied to the other di-allelic locus types (i.e., neutral markers, deleterious mutations, and BDMIs). This optimisation drastically reduces the memory footprint of di-allelic loci. It also enables hardware-accelerated operations on large sets of loci, which drastically reduce computation time, especially for QTN models with equal allelic effects across many loci (Fig. 4b).

When choosing a simulation tool, performance, ease of use, and flexibility are usually considered together, and often trade-off with each other. While SLiM offers simple parametrisation of population genetics models based on the infinite-site model with high performance and high scripting flexibility, it is not well geared to model polygenic traits the way Nemo does, as exemplified by our benchmark. Similar endpoint results can be obtained but at a higher computational and scripting price for SLiM. Scripting-based simulation software may suffer from lower computational efficiency when large parts of the process are based on interpreted rather than pre-compiled code, for instance for phenotype calculations involving non-additive interactions across many loci. That said, the differences in performance between SLiM and Nemo that we have shown are more likely caused by the underlying difference in data structure between the “sparse” and “dense” genetic architectures than by the execution of interpreted code. Sparse models are generally better suited for population genetics than quantitative genetics models, where Nemo is a strong contender. For ease of use, Nemo facilitates model implementation through a set of standardized parameters, and pre-compiled code. This reduces the risk of coding error when scripting large and complex models and ensures the consistency of model outcomes necessary for standardization and comparison across studies, since each scripted implementation may differ in subtle, sometimes undetected ways. The flexibility in Nemo comes from its modular design and an extended set of features: the metapopulation backbone, the LCEs, and the *traits* (i.e., the different genotype-phenotype-fitness mappings).

Nemo is designed around the metapopulation container and the modular Life Cycle Events. Older simulators often enforced a hard-coded succession of events (i.e., reproduction-selection-migration, as in quantiNEMO; Neuenschwander et al., 2019), a choice mainly derived from the Wright-Fisher model. In contrast, Nemo allows users to directly define the sequence of events, a flexibility present since Nemo 2.0 (Guillaume and Rougemont, 2006). This enables variation of population models with seamless transitions between soft selection (relative fitness determines reproductive output) and hard selection (absolute fitness determines patch size), or between Wright-Fisher (soft) and non-Wright-Fisher (hard) models. Nemo 2.4 internally capitalizes on optimisations for the Wright-Fisher population model, activated by a single parameter switch. Non-Wright-Fisher models, on the other hand allow for a wider variety of population dynamics essential to eco-evolutionary models with stochasticity. In Nemo, population regulation is explicit and parametrized as a ceiling model (carrying capacity caps patch size), a Beverton-Holt model with density-dependent regulation, or both (as in Nemo-Age; Cotto et al., 2020). This enables the simulation of extinction and recolonization dynamics, “source-sink” meta-population structures, and evolutionary rescue scenarios where population size is a dynamic variable dependent on absolute fitness.

Nemo 2.4 supports finer biological nuances with separate LCEs for gametic migration (`breed_disperse`, akin to pollen flow) and zygotic migration (`seed_disperse`). This distinction allows researchers to model complex life histories, such as those of plants where gene flow occurs at different stages with different kernels. In general, alternative timing of the LCEs broadens the range of models that can be implemented. Nemo-Age further adds the capacity to model age- and stage-structured populations with complex life cycles (Cotto et al., 2020), and incorporates the quantitative trait implementation presented here (Nemo-Age v0.33). Nemo and Nemo-Age will ultimately be merged in a single software.

A further key design strength of Nemo 2.4 is its modular *trait* system built around a common genetic map encompassing multiple chromosomes. This allows researchers to mix the genetic elements of the available traits–neutral, deleterious, incompatibility (BDMI), and quantitative trait loci–on a shared map and utilize empirically-determined recombination maps to generate realistic patterns of polymorphism at the genomic level. This design enables the study of linkage disequilibrium (LD) between different classes of genomic elements without the need for elaborated scripting. The shared genetic map ensures that recombination events affect all traits consistently. This integration is vital for simulating “hitchhiking” effects (e.g., Kemppainen and Guillaume, 2026), where neutral diversity is eroded by selection on linked functional loci, or “background selection,” where purifying selection on deleterious mutations reduces diversity at linked neutral sites (e.g., Grossen et al., 2020). By treating these elements as distinct modules that interact via the recombination map, Nemo 2.4 facilitates the generation of synthetic genomic datasets that mirror the complexity of empirical data used in GWAS (e.g., Chebib and Guillaume, 2021; Kemppainen and Guillaume, 2026). Moreover, several built-in file formats for saving and retrieving genetic data ensure interoperability with empirical genomics pipelines, in particular PLINK 1.9 format for GWA and other genomic analyses (Chang et al., 2015).

In conclusion, Nemo 2.4 is an ideal simulation platform to model eco-evolutionary dynamics based on quantitative traits evolution and answer many questions in evolutionary quantitative genetics. It offers an extensive implementation of quantitative traits capable of modelling complex genotype-phenotype maps with modular pleiotropy, dominance, and epistasis beyond what is available in other simulation tools. The integration of multivariate phenotypes with multivariate selection surfaces enables researchers to study the effects of correlational selection on patterns of genetic variation at the genomic level. Nemo’s high flexibility in freely assembling existing modules, LCEs and traits, its mini-language developed for parameter grid specification, and its higher across-the-board computational performances make it an ideal tool to model large populations under complex demo-genetic scenarios in large simulation campaigns. The modular architecture of Nemo’s C++ code base enables seamless addition of new components, LCEs and traits, aided by development guidelines available on the project’s repository. Moreover, because of its research-driven development, Nemo’s simulation results have been thoroughly tested and validated against theoretical expectations, some of which were presented in this paper. Additional validations include two-patch equilibrium phenotypic divergence with gene flow in one (Yeaman and Guillaume, 2009) or two correlated quantitative traits (Guillaume, 2011), correlated selection on varying modular genetic architectures (Chebib and Guillaume, 2017), as well as models including phenotypic plasticity (Schmid and Guillaume, 2017). Equilibrium genetic covariance between traits affected by pleiotropic loci also matches theoretical predictions (see Lande, 1980), although not when non-pleiotropic loci are fully linked, demonstrating that linkage alone does not replicate the effects of pleiotropy, as show by Chebib and Guillaume (2021). Future implementations will provide increased flexibility and improved performances. Current API development will enable embedding Nemo in python scripting environments and link it with existing and future data repositories for forward-time simulations, such as stdpopsim (Gower et al., 2025).

## Acknowledgments

We would like to thank all previous and current users of Nemo who helped us improve the software. Multiple colleagues participated in developing various aspects of the code. We thank Sam Yeaman, Kim Gilbert, and Mike Whitlock for their invaluable input during the development of the quantitative trait. Camilla Stefanini, Katalin Csilléry, Nadja Verspagen, and Petri Kemppainen helped push Nemo’s capacities into new territories.

## Data accessibility

Git repository on bitbucket: https://bitbucket.org/ecoevo/nemo-release/. Website: https://nemo2.sourceforge.io/.

## Supplementary Information

## Supplementary figures

**Supplementary Figure 1:**
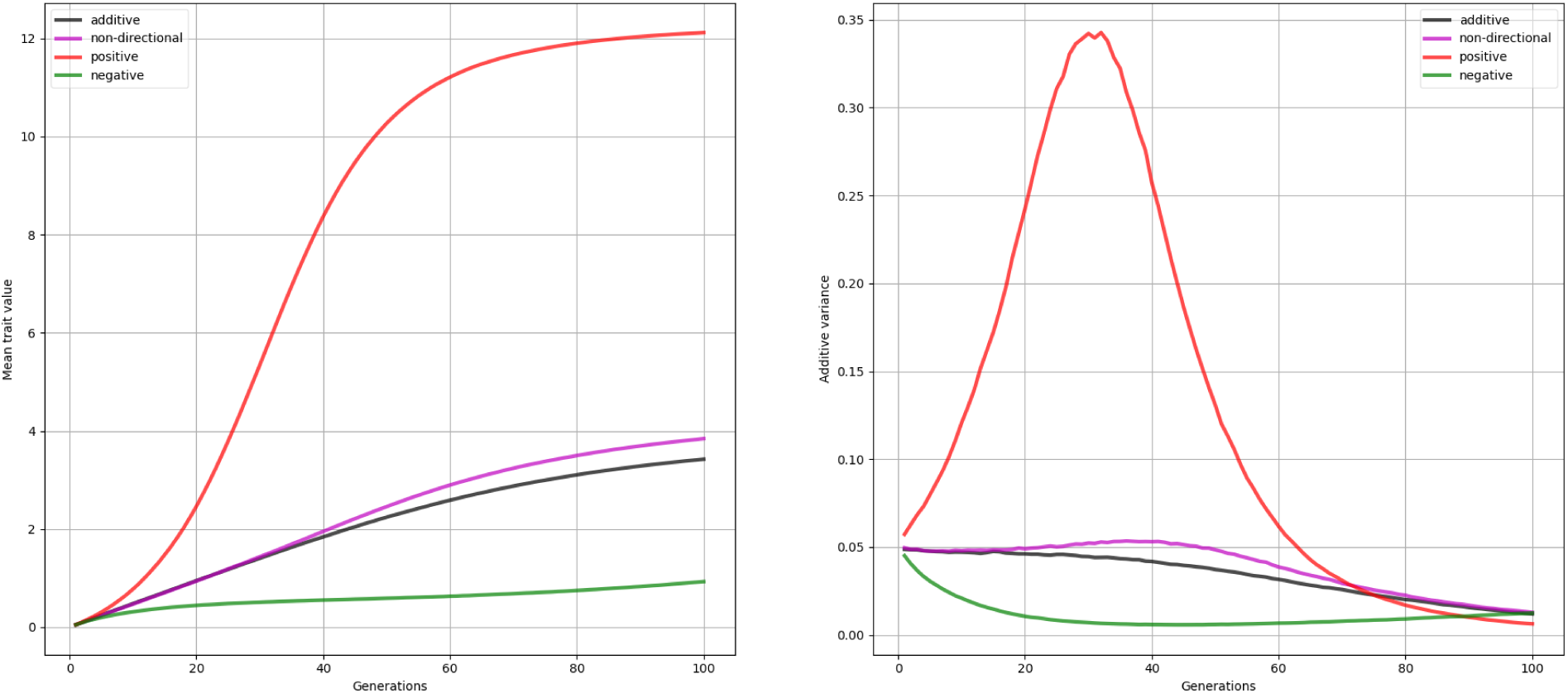
Expected evolution of trait mean and additive variance with and without epistasis. The figure reproduces the theoretical expectations of Carter et al. (2005) (see their Figure 1). The quantitative trait is controlled by 20 continuum-of-allele loci with additive effects (black), non-directional epistasis (purple), negative epistasis (green), and positive epistasis (red). The trait is under linear selection for 100 generations in a Wright-Fisher population with N=1000, and without mutation. The simulations shows the proportionality of *V_a_* between the two sets of loci, with *L* = 1000 on the left and *L* = 10000 on the right.

**Supplementary Figure 2:**
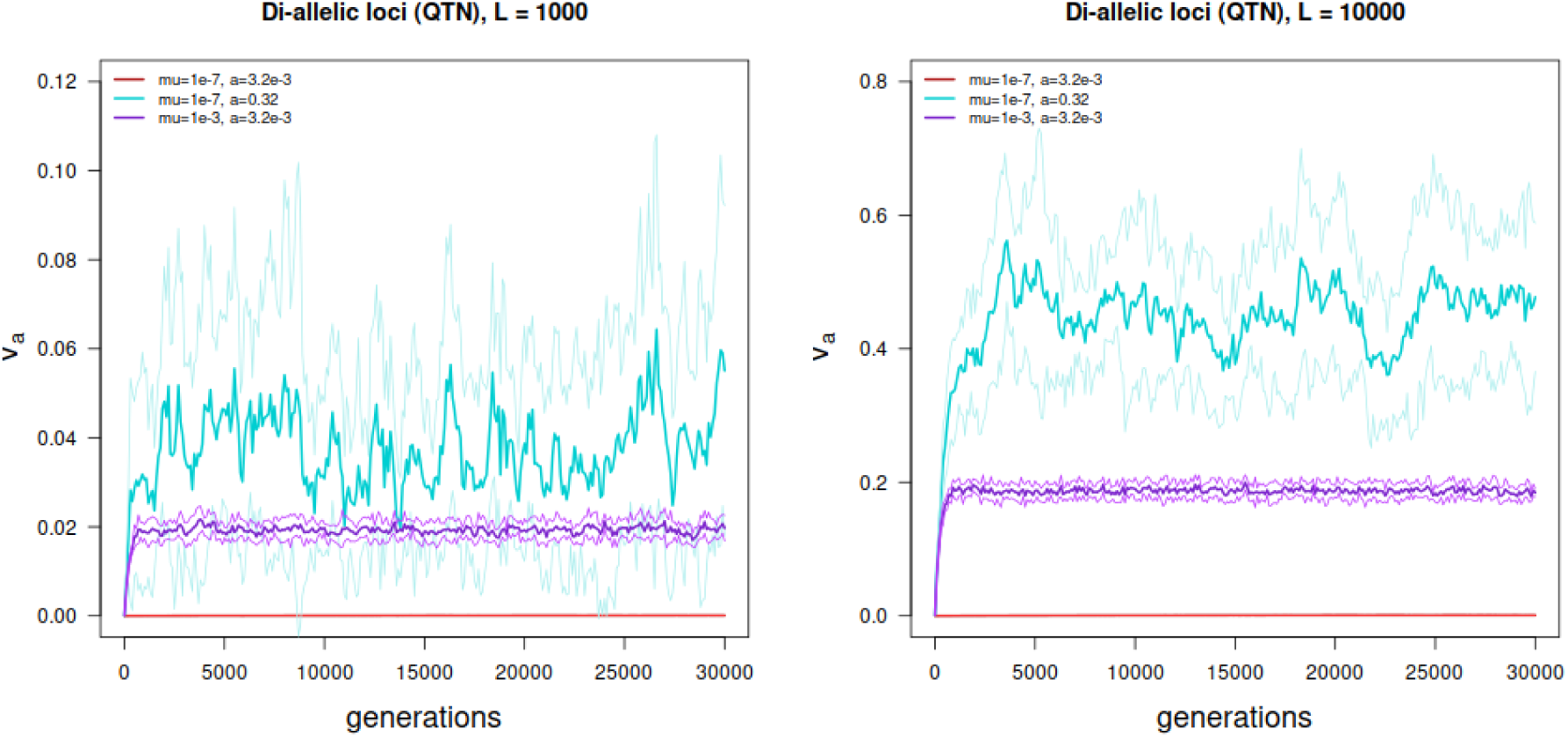
Equilibrium genetic variance *V*_A_ of a quantitative trait coded by *L* = 1000 (left) and *L* = 10000 (right) di-allelic loci, all with same allelic effect *a* = {0.003162, 0.3162}, and recombination rate *r* = 10*^−^*^5^. The mutation rate varied between *µ* = 10*^−^*^7^ (red and cyan) and *µ* = 10*^−^*^3^ (purple).

**Supplementary Figure 3:**
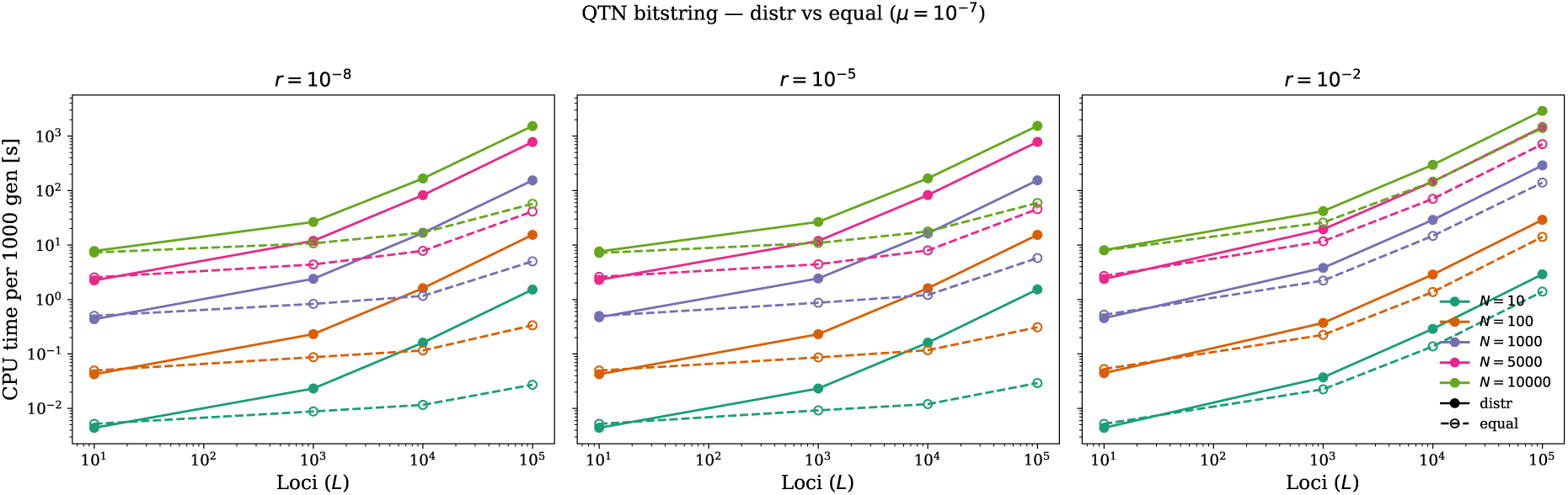
Execution time (CPU time) of the QTN bitstring architecture for three recombination rates and *µ* = 10*^−^*^7^. Details of the simulation settings and execution environment are in the main text.

**Supplementary Figure 4:**
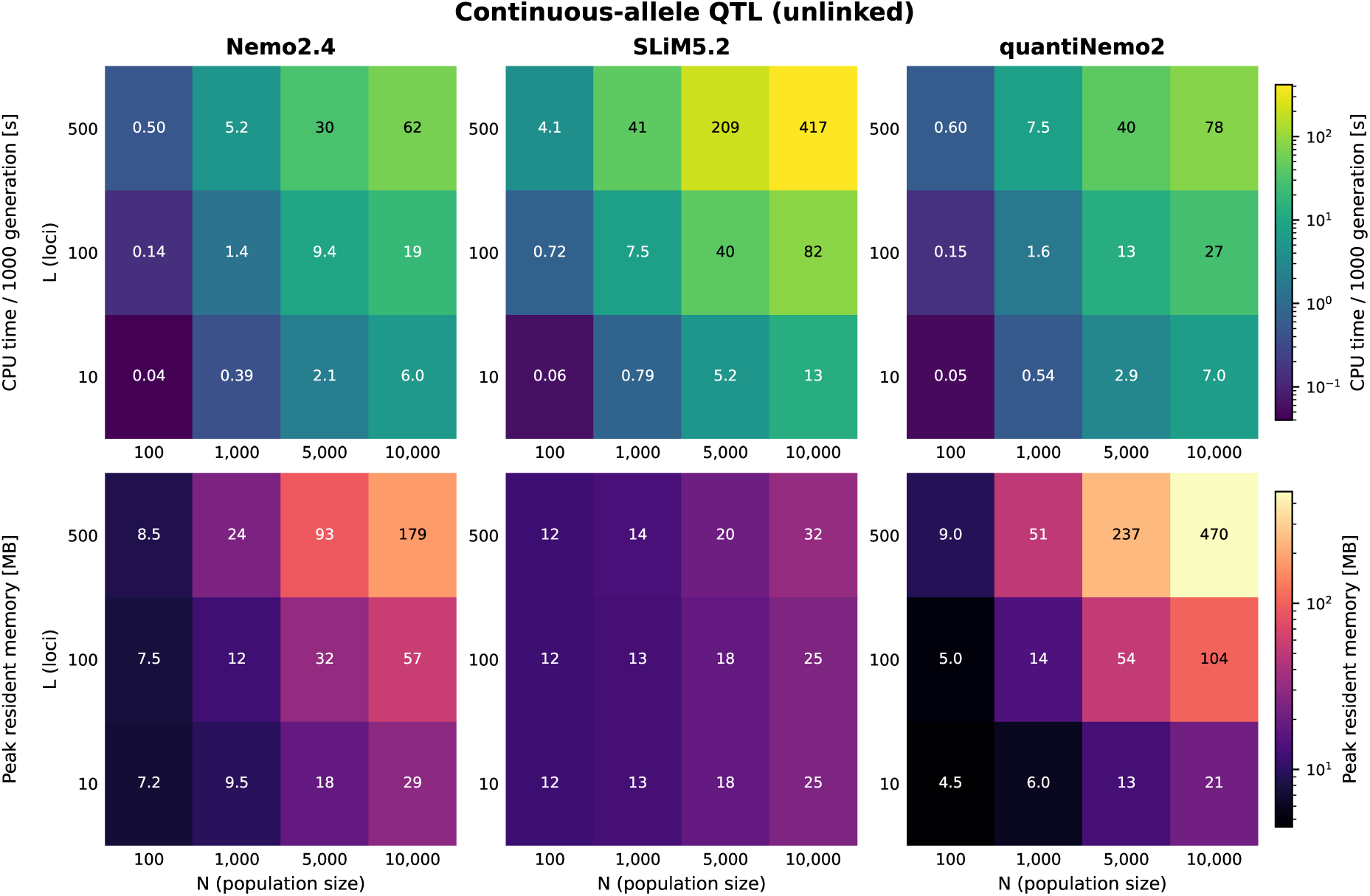
**Software comparison** benchmarking mean CPU time to 1000 generations (seconds) and mean peak resident memory (MB RAM) of nemo2.4, quantiNemo2, and SLiM5.2 over 5 runs. The model simulated one WF population under Gaussian stabilizing selection with *V*_S_ = 10 and *V*_E_ = 1. The quantitative trait under selection is encoded by *L* unlinked QTL under the continuum-of-allele model (`quanti_allele_model` continuous in Nemo) with *µ* = 1*e^−^*^4^ and mutational variance *α*^2^ = 0.05. The scripts are provided in appendix.

**Supplementary Figure 5:**
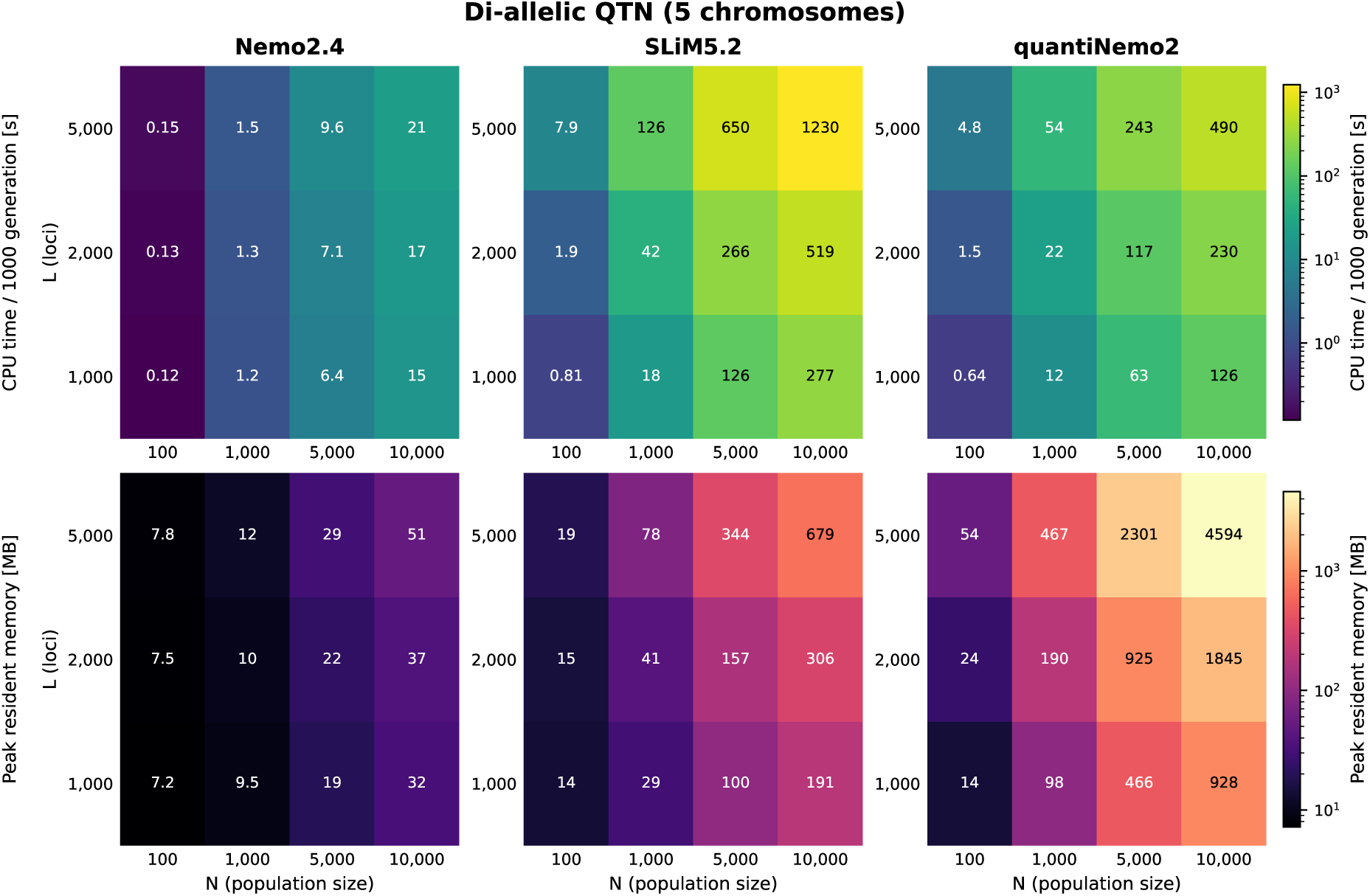
**Software comparison** benchmarking mean CPU time to 1000 generations (seconds) and mean peak resident memory (MB RAM) of nemo2.4, quantiNemo2, and SLiM5.2 over 5 runs. The model simulated one WF population under Gaussian stabilizing selection with *V*_S_ = 10 and *V*_E_ = 1. The quantitative trait under selection is encoded by *L* di-allelic QTNs linked on a genetic map with 5 chromosomes and *r* = 1*e^−^*^5^ on each of them. Alleles at each locus had effects ±0.01 with *µ* = 1*e^−^*^5^. The scripts are provided in appendix.

**Supplementary Figure 6:**
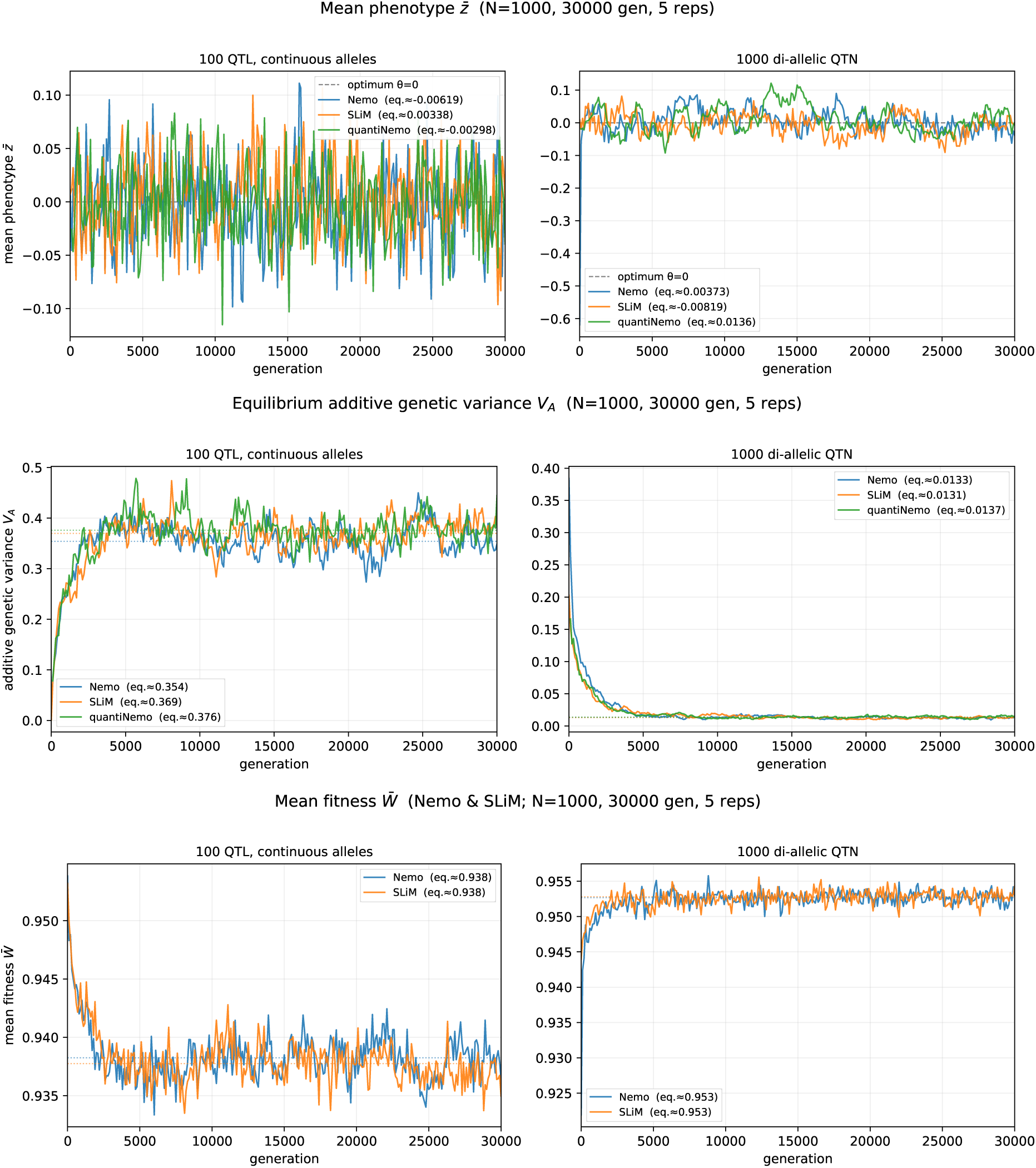
**Software comparison** showing the equilibrium values of the mean phenotype (top), the additive genetic variance *V*_A_ (middle), and mean population fitness (bottom) in one WF population with *N* = 1000, for a quantitative trait encoded by 100 unlinked QTL (left column) or 1000 linked di-allelic QTNs (right column). The simulation were run for 30,000 generations and 5 replicates with Nemo2.4.1, SLiM5.2, and quantiNemo2. Other parameters as in previous figures. The equilibrium values displayed on the graphs are averages computed over generations from 10,000 to 30,000.

## Supplementary tables

**Supplementary Table 1:** CPU time of the QTN model with *equal* allelic values at all loci in seconds per 1000 generations evaluated on 5 replicates. The reported time is *wall time* (user + system). The table shows the value of the population size (*N*), and number of loci (*L*) while keeping recombination rate at *r* = 10*^−^*^8^ and averaging over different mutation rates (see figure 4 in main text). The table also show the CPU time ratio relative to the *distr* QTN model and the multi-allelic QTL model.

| N | L | NL | CPU s/1000 gen | $\frac{\text{QTN-dist}}{\text{QTN-equal}}$ | $\frac{\text{QTL}}{\text{QTN-equal}}$ |
| --- | --- | --- | --- | --- | --- |
| 10 | 10 | 100 | 0.005 | 0.92 | 1.00 |
| 10 | 1000 | 10000 | 0.01 | 2.66 | 2.54 |
| 10 | 10000 | 100000 | 0.01 | 12.48 | 11.00 |
| 100 | 10 | 1000 | 0.05 | 0.92 | 1.05 |
| 100 | 1000 | 100000 | 0.09 | 2.69 | 7.91 |
| 100 | 10000 | 1000000 | 0.13 | 12.41 | 191.24 |
| 1000 | 10 | 10000 | 0.47 | 0.94 | 1.45 |
| 1000 | 1000 | 1000000 | 0.85 | 2.80 | 37.66 |
| 1000 | 10000 | 10000000 | 1.36 | 12.29 | 184.77 |
| 5000 | 10 | 50000 | 2.46 | 0.93 | 5.70 |
| 5000 | 1000 | 5000000 | 4.49 | 2.70 | 36.06 |
| 5000 | 10000 | 50000000 | 8.59 | 9.65 | 150.39 |
| 10000 | 10 | 100000 | 7.58 | 1.08 | 4.53 |
| 10000 | 1000 | 10000000 | 10.95 | 2.43 | 31.40 |
| 10000 | 10000 | 100000000 | 18.41 | 9.13 | 100.22 |

## 1 Examples

### 1.1 Gaussian stabilizing selection in a WF population

Simulation parameters for one of the validation simulations shown in Figure 1 in the main text. A single WF population is simulated with hermaphrodite individuals carrying 300 QTL under the contiuum-of-allele model. The quantitative trait is under weak stabilising selection for a phenotypic value of 0.

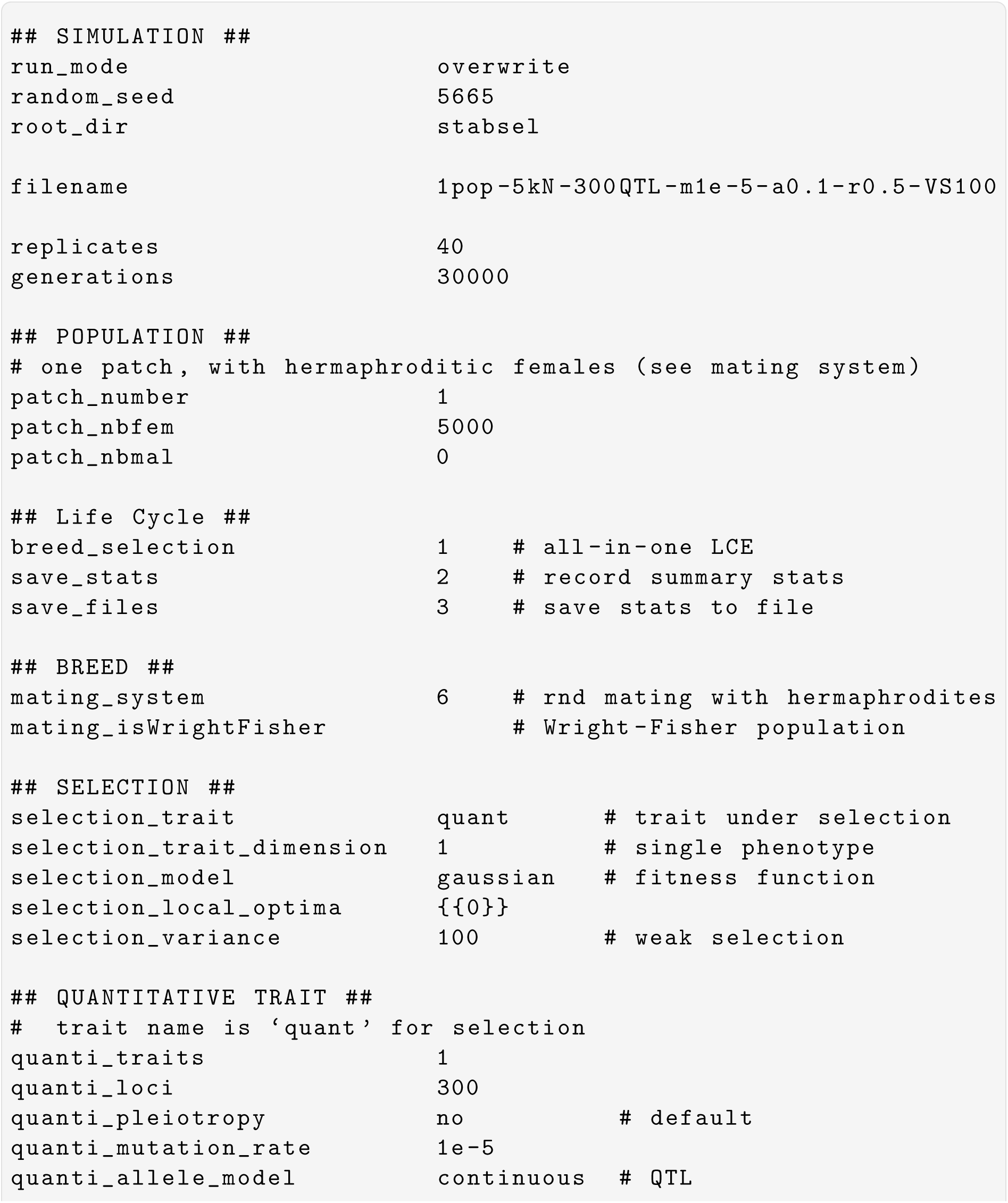
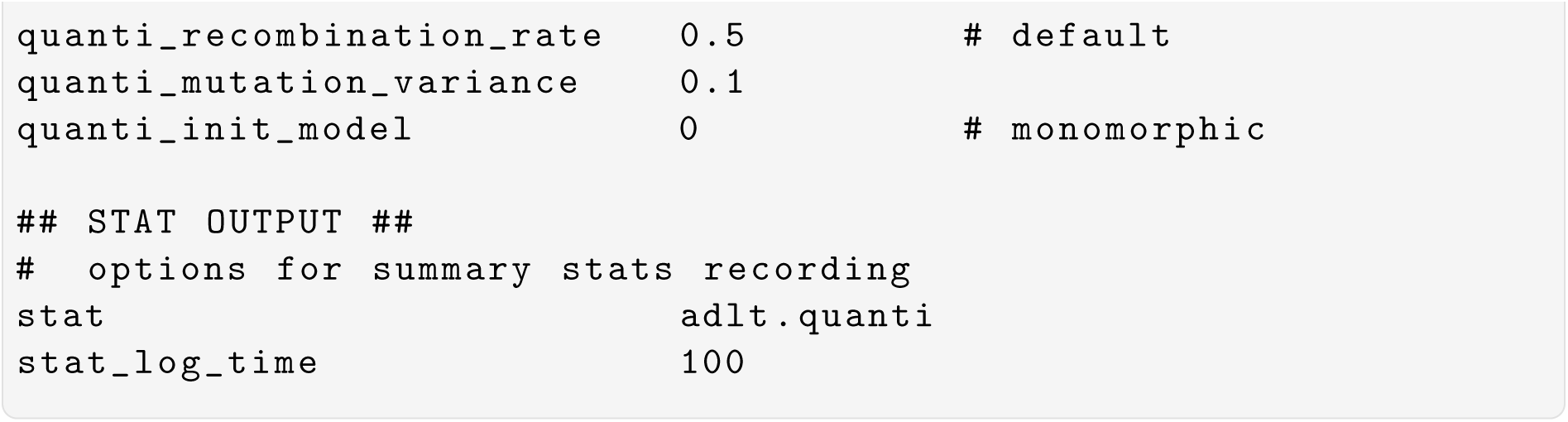

### 1.2 Evolutionary rescue after an environmental shift

This simulation is for a model of evolutionary rescue in a single population adapting to a local phenotypic optimum set at 1 originally, and moved to 3.2 after 5000 generations of stabilizing selection (burn-in phase). The simulation is run for 100 additional generations during which the population goes through a rapid decline in density, before being rescued by adaptive change of the trait, eventually reaching its new optimum value. The resulting demography is shown in Figure below.

**Supplementary Figure 7:**
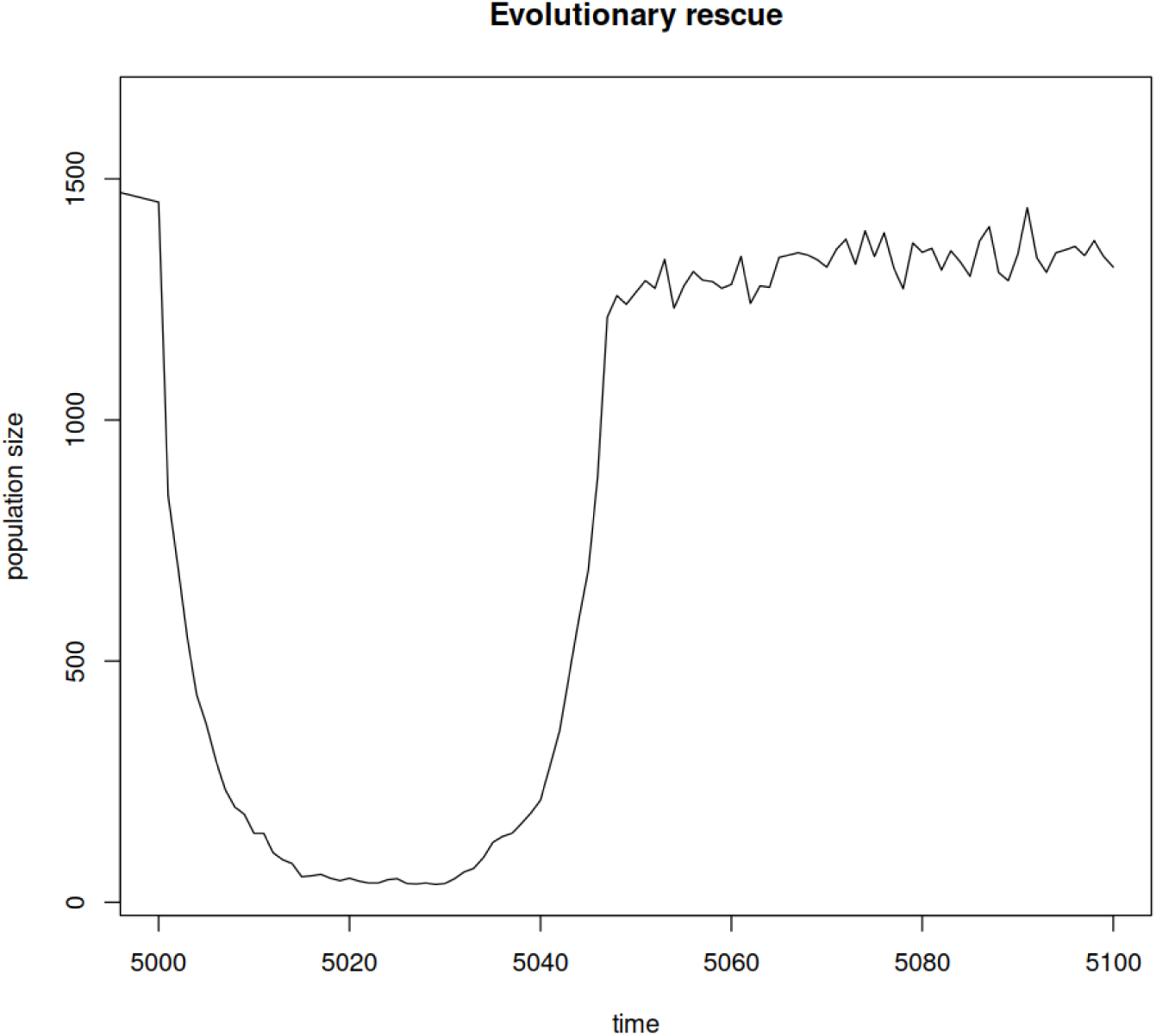
Example of an evolutionary rescue. The graph shows the change of population size over time after the local phenotypic optimum changes from 1 to 3.2 at generation 5001. The population is nearly extinct around generation 5020 but rebounds as the trait adapts to its new optimum. Result is shown for a single replicate.

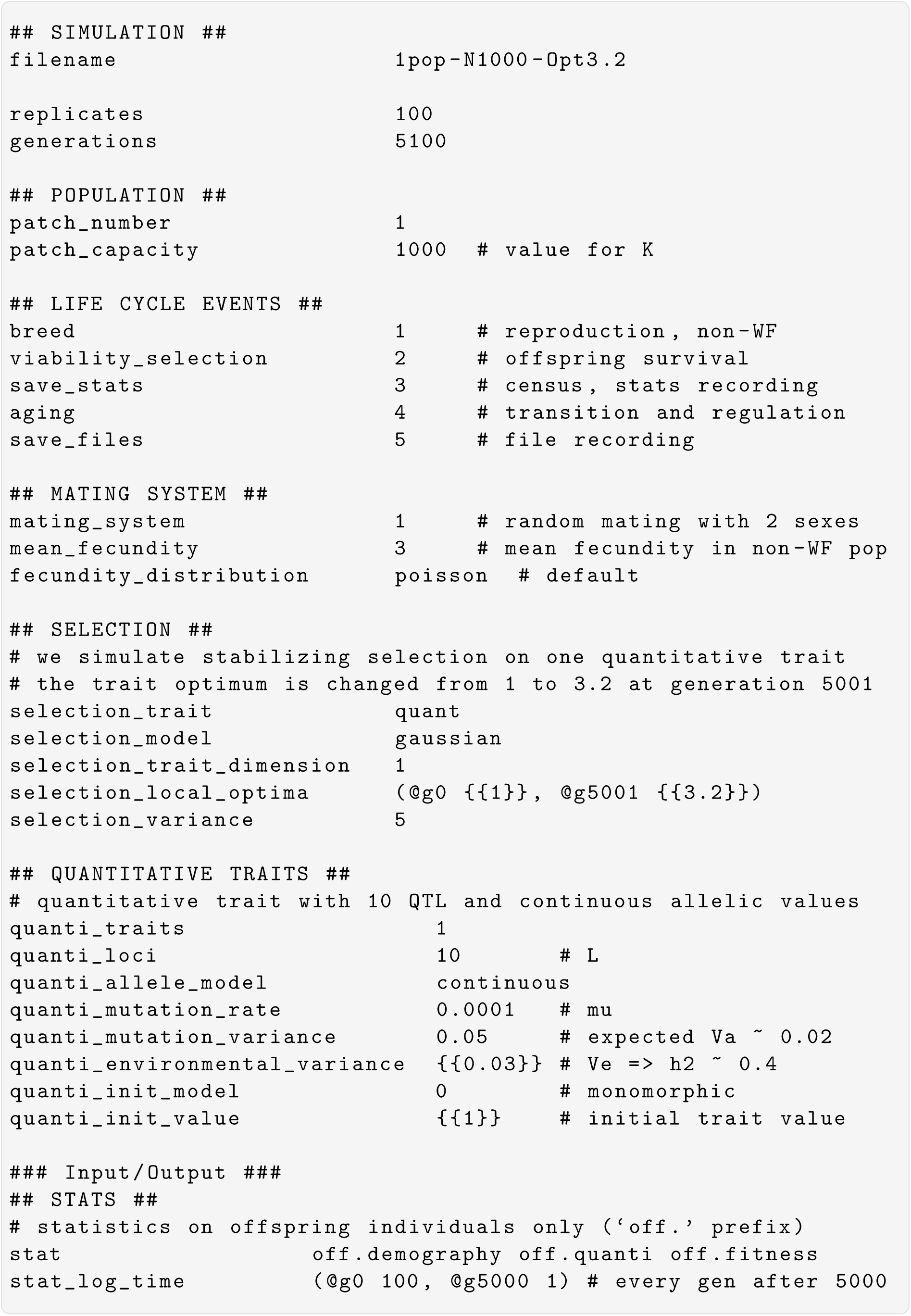

### 1.3 Species’ range expansion on a 2D lattice

A single population of 100 hermaphrodite females initially occupies the left column of a 10 × 50 lattice. Forward dispersal to the four nearest neighbours produces a colonisation wave across the grid. Local Gaussian selection acts on a single quantitative trait, with local optima forming a cline along the column axis (1, 2, …, 50 from left to right), so that establishment in newly reached patches requires adaptive change. The lattice is parametrised with the pre-set 2D lattice dispersal model (`dispersal_model` 4) and reflective boundaries. See Manual, section 1.4, for details on dispersal models.

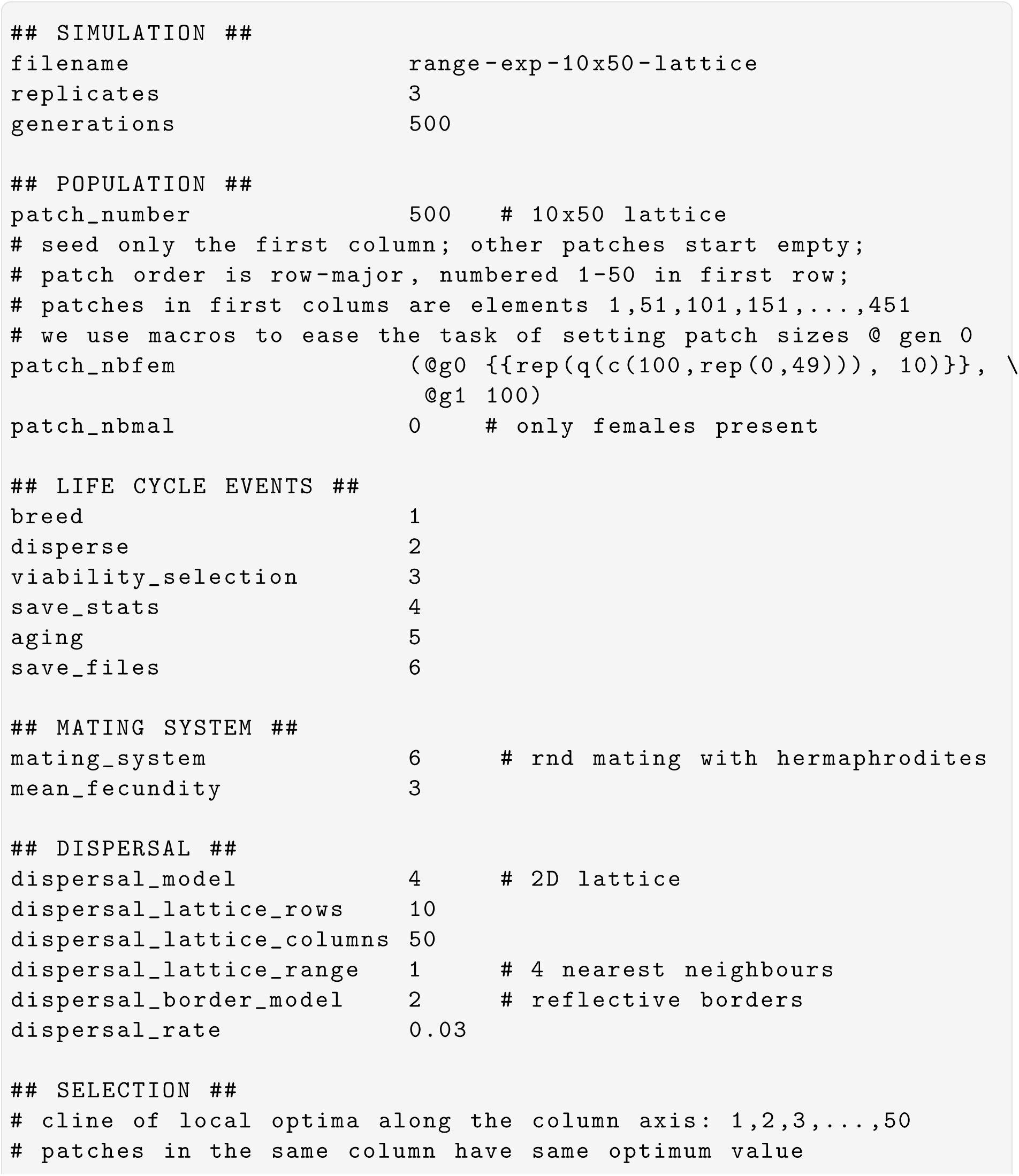
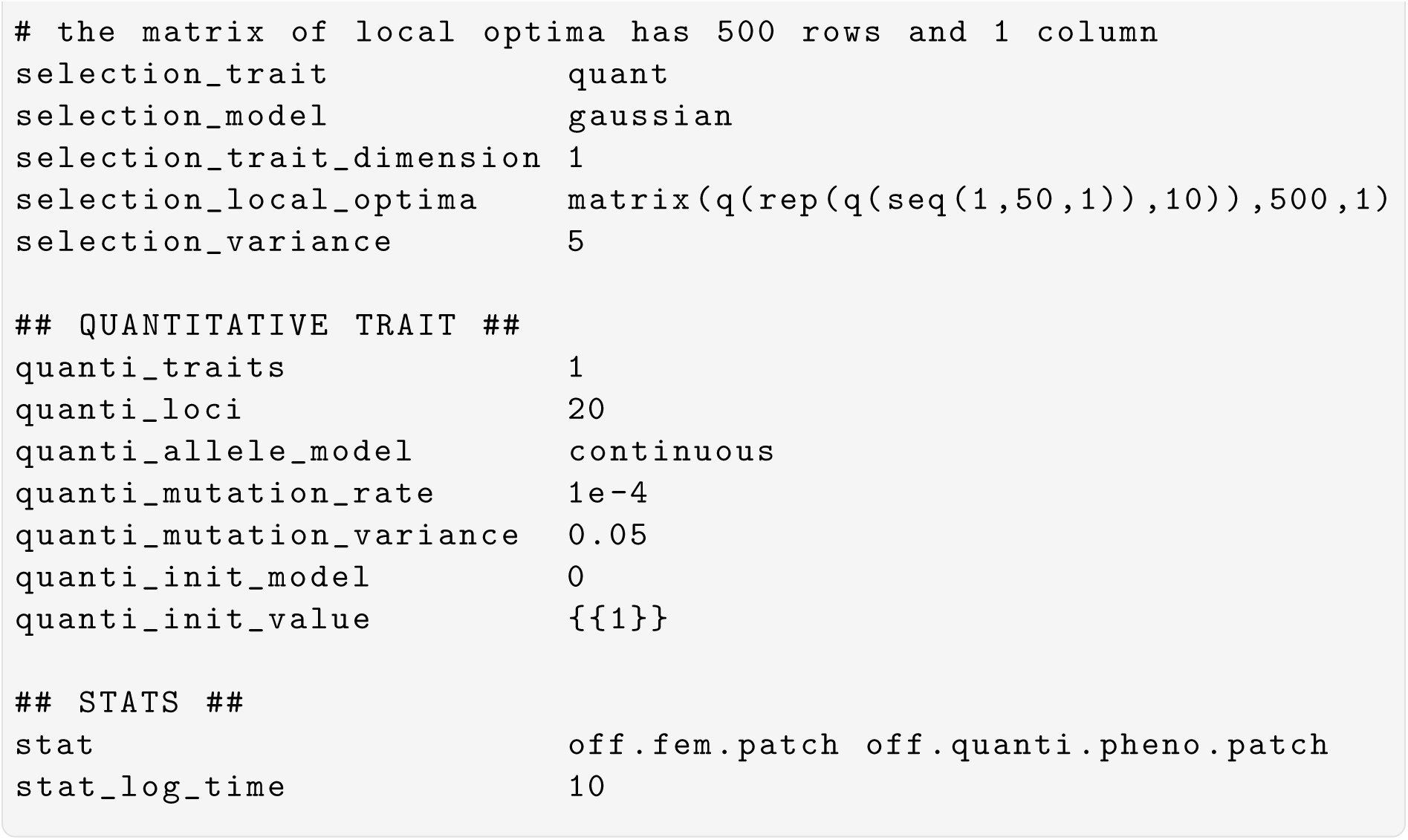

### 1.4 Linear selection on two epistatic networks

This example illustrates the network form of the epistasis parametrisation. Twenty QTL are organised in two independent modules of 10 loci (cliques), with no interaction across modules. Each module generates 10 × 9*/*2 = 45 pairwise epistatic coefficients, drawn here from a normal distribution with negative and positive mean (directional epistasis). The trait is under linear directional selection in a Wright-Fisher population (no mutation), as described in the Selection subsection of the main text.

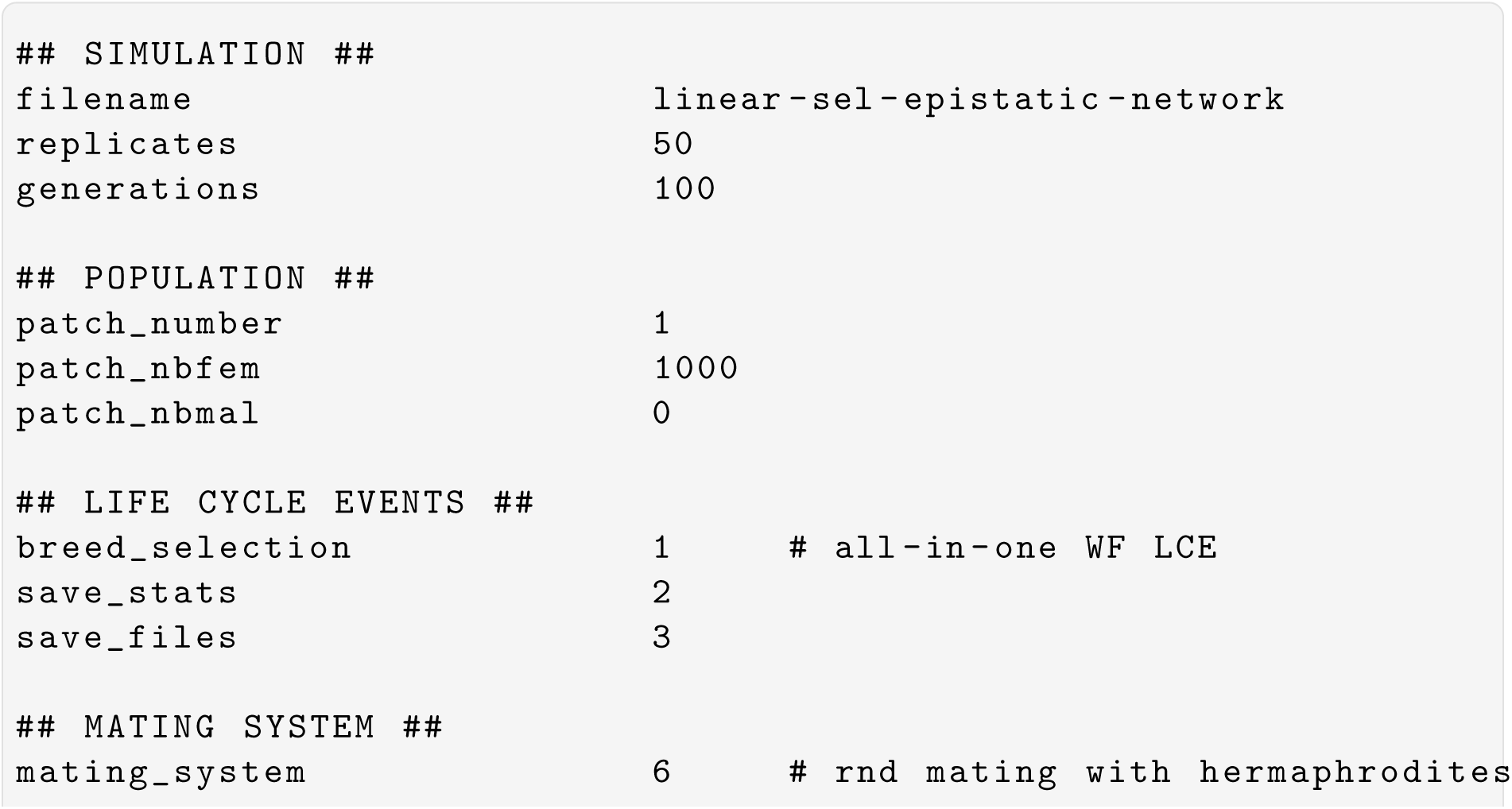
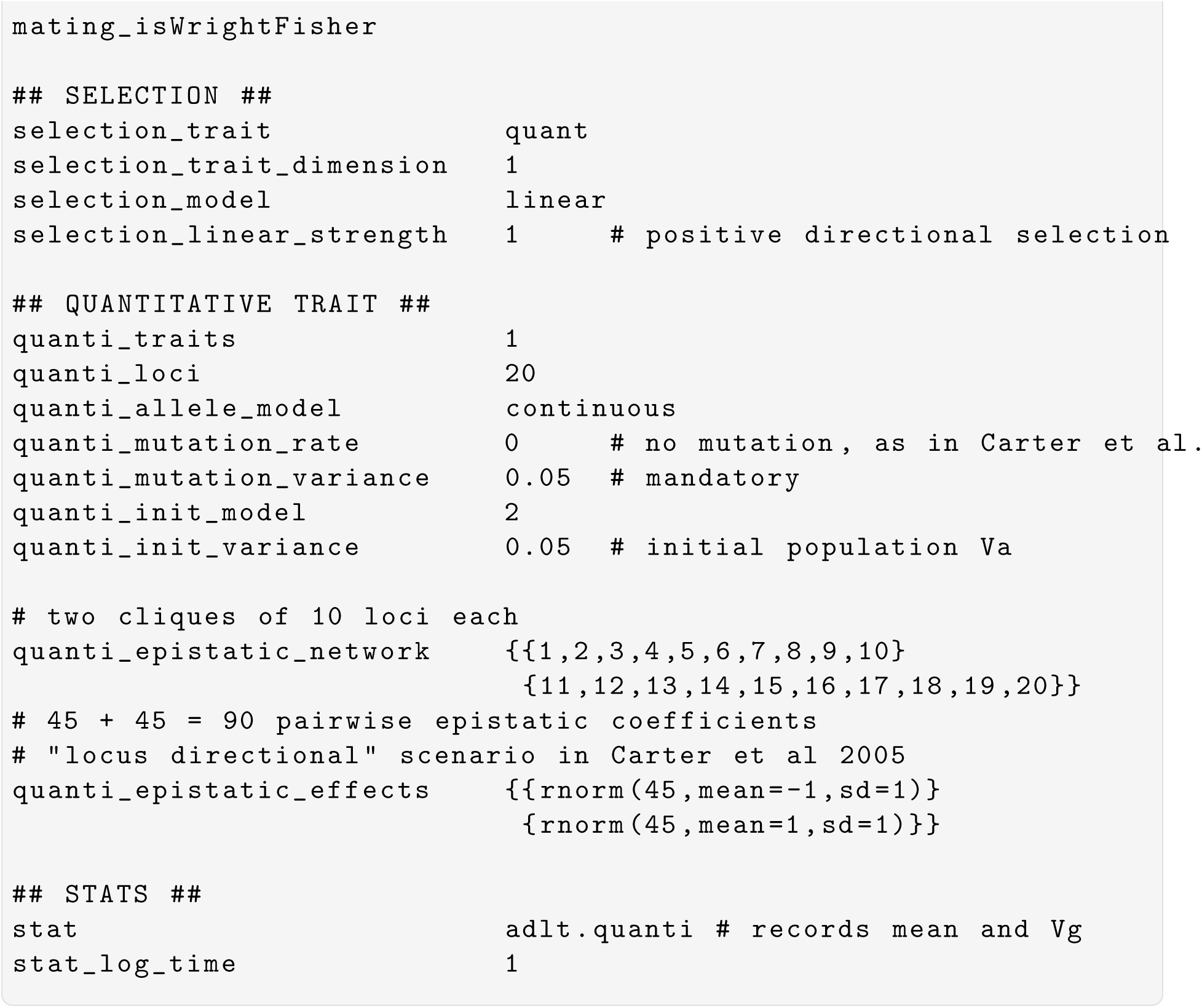

This example illustrates the network form of the epistasis parametrisation, inspired by the theoretical scenario of Carter et al. (2005) that is also reproduced in Supplementary Figure 1. Twenty QTL are organised in two independent modules of 10 loci (cliques), with no interaction across modules. Each module generates 10 × 9*/*2 = 45 pairwise epistatic coefficients, drawn here from a normal distribution with negative mean (negative epistasis). The trait is under linear directional selection in a Wright-Fisher population (no mutation), as described in the Selection subsection of the main text.

### 1.5 Plastic phenotype with a shifting environmental cue

This example decouples the environmental cue from the selective optimum, a biologically realistic scenario of climate change where the cue (e.g., photoperiod) remains stable across generations while the selective optimum (e.g., temperature) shifts directionally. The plastic phenotype, built from an evolving intercept *g*_0_ and a fixed slope *g*_1_ via the linear reaction norm *z* = *g*_0_ + *g*_1_(*e* − *g*_2_), cannot therefore track the shifting optimum through plasticity alone: adaptation must proceed through evolution of *g*_0_. The cue is held fixed at 0 while `selection_local_optima` ramps from 0 to 2 over 1000 generations via a per-generation rate of change. See the *Phenotypic plasticity and liability* subsection in the main text, and Manual section 2.5.

This example will generate three different simulations, one for each value of the parameter `pheno_plastic_g1_value`, which we have set to 0.0, 0.5, and 1.0. To know which output file corresponds to which simulation, and avoid overwriting results from previous simulations, we use the parameter value recall string ‘%1’ to insert the value into the file-name. The parameters receiving several values are called *sequential* parameters, and are referenced in alphabetic order in the recall argument. The example below will generate three filenames:

`plastic-slope0.0-shifting-optimum`,

`plastic-slope0.5-shifting-optimum`,

`plastic-slope1.0-shifting-optimum`.

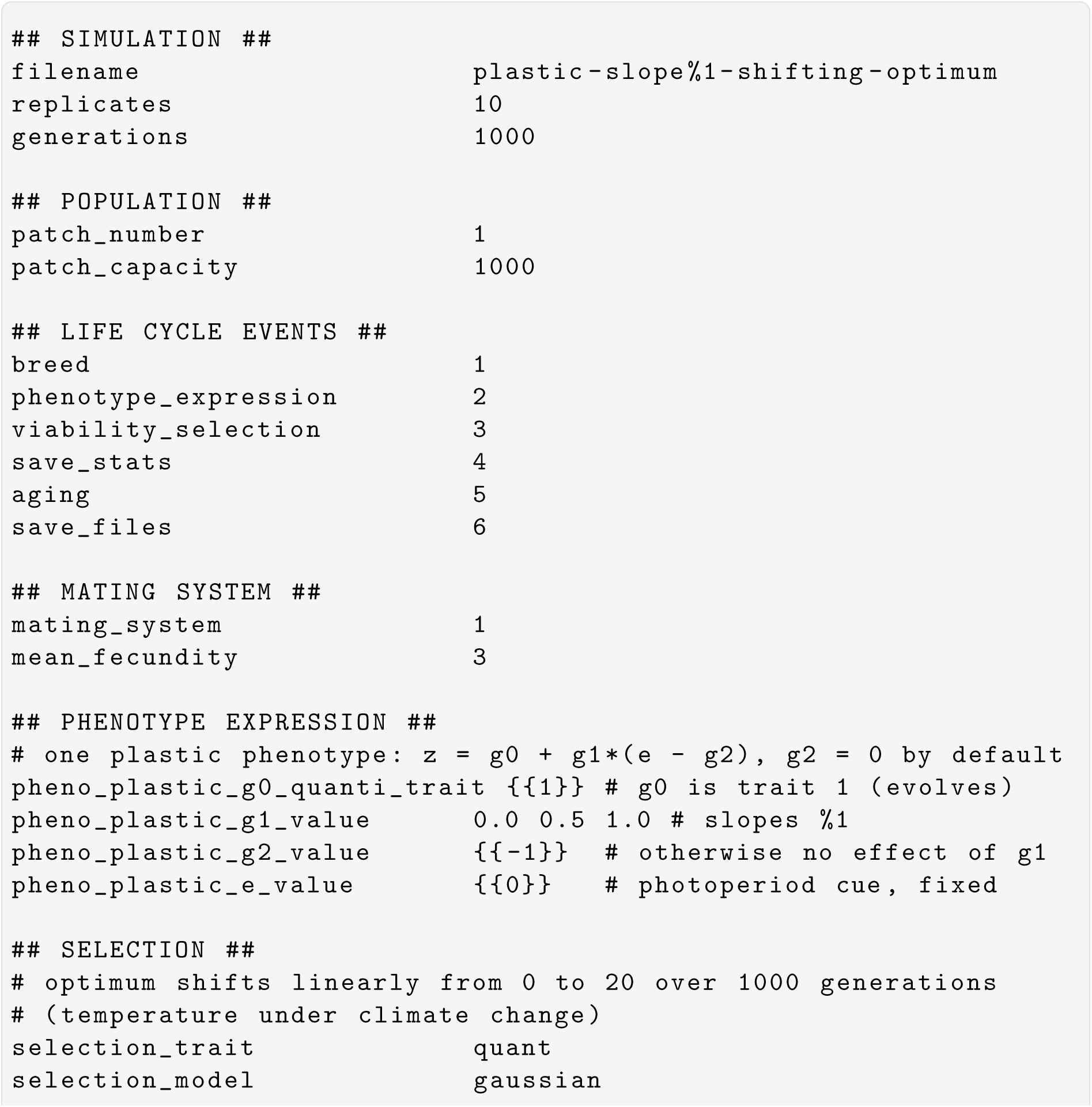
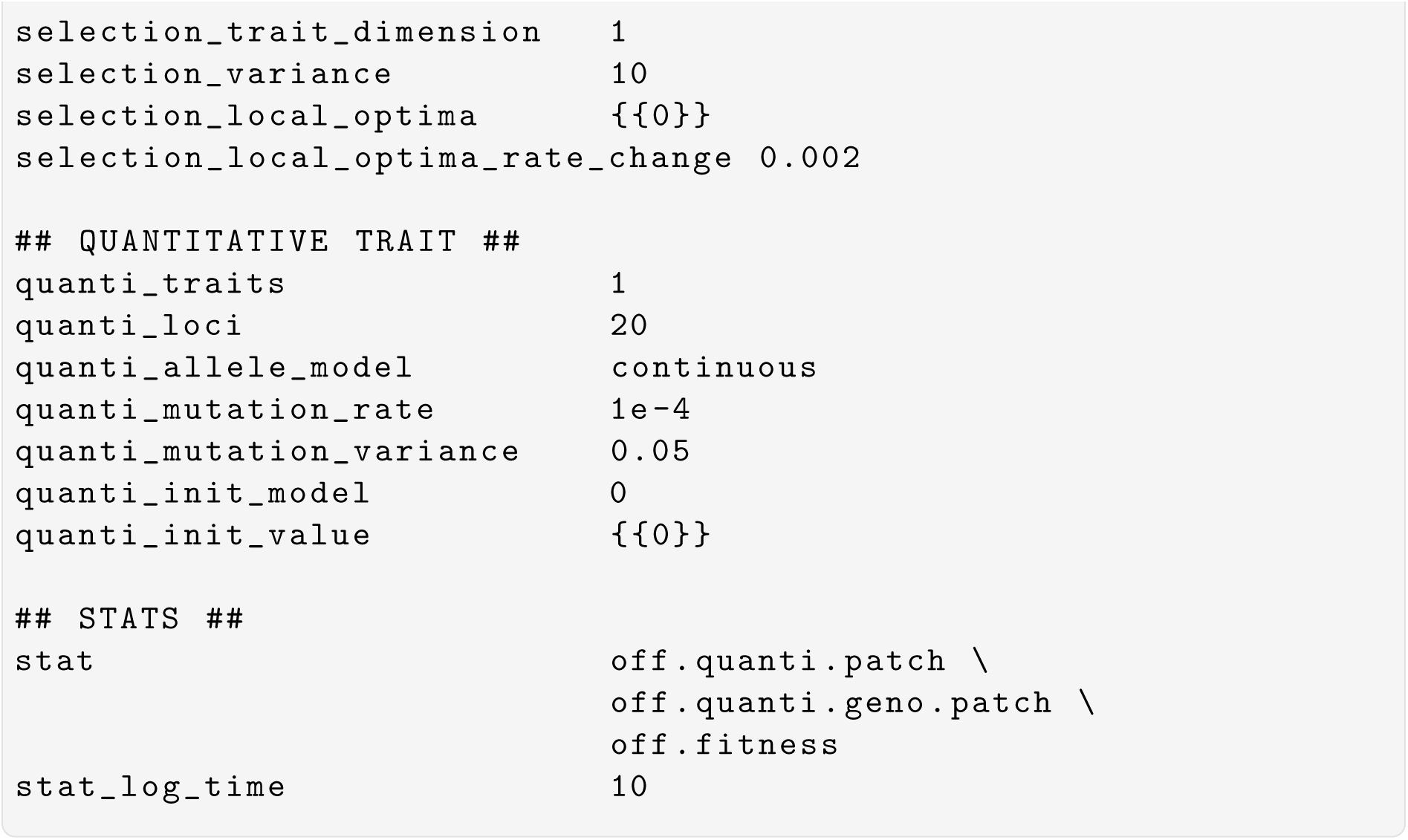

### 1.6 Plastic threshold trait with underlying liability

A dichotomous phenotype (e.g., presence/absence of a morphological feature) is expressed through a continuous polygenic liability with a plastic NoR. A single fixed threshold at 0 partitions individuals into two discrete phenotypic classes (0 and 2). This phenotype tracks a local optimum of {{0,2}} in patch 1 and 2, respectively. A second NoR trait tracks a second optimum with values {{-2,0}} in patch 1 and 2, respectively. Both NoR’s have an evolving slope encoded by two separate quantitative traits (3 and 4). Gaussian selection models two shifting trait optima with decreasing values during the first 2500 generations after which the shift is reversed to positive. See the Phenotypic plasticity and liability subsection in the main text, and Manual section 2.6.

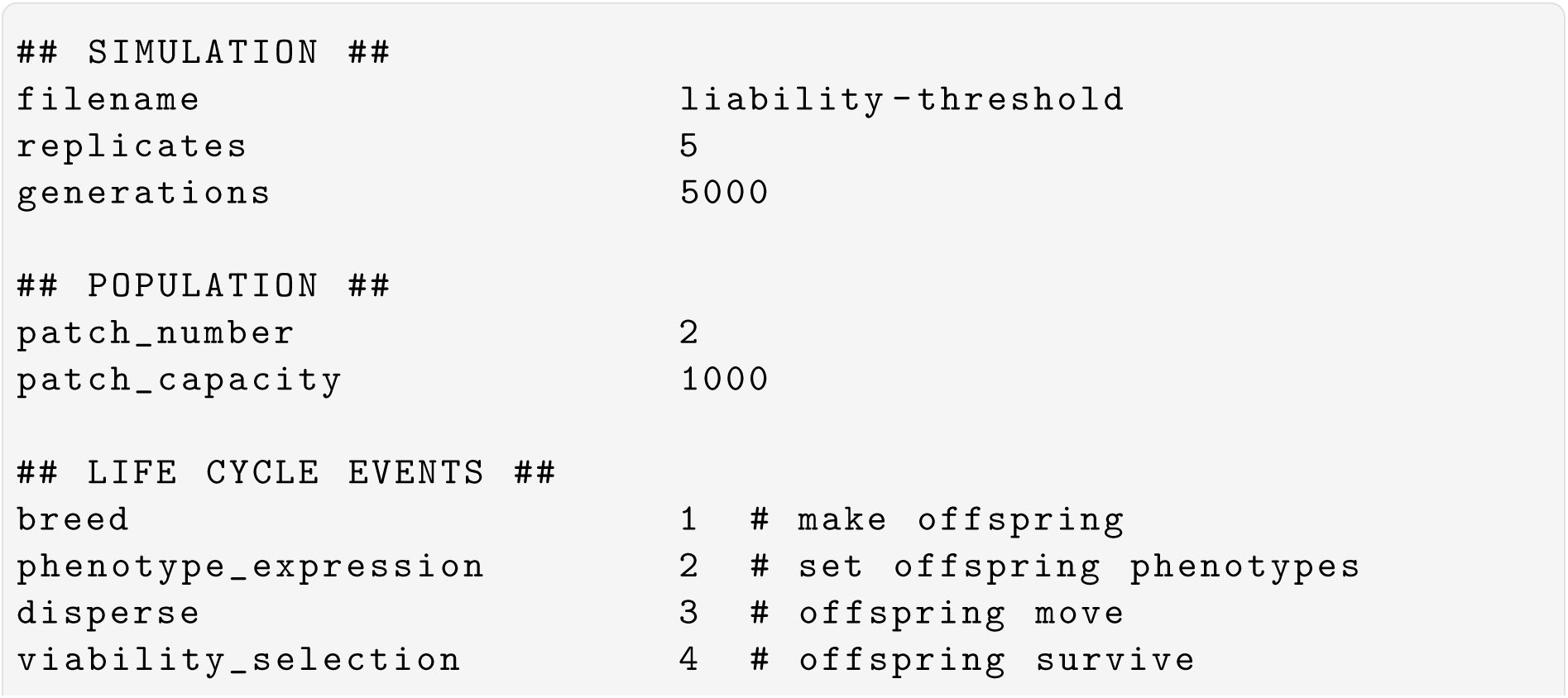
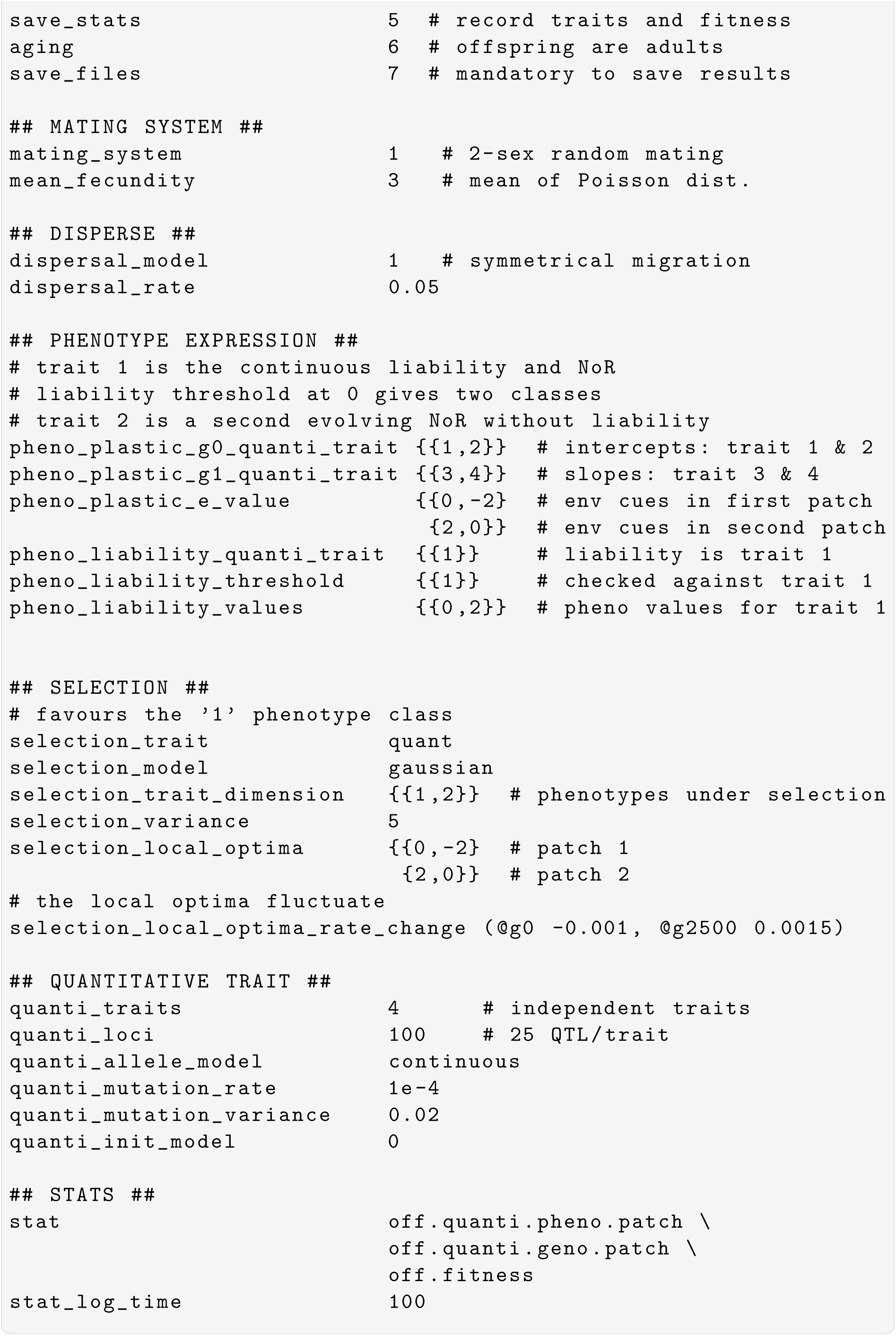

### 1.7 Crossing: half-sib/full-sib cross

A classic North Carolina I (half-sib/full-sib) crossing design is applied after a burn-in of stabilising selection on the quantitative trait. Ten sires are each mated with five dams, each producing ten offspring, enabling decomposition of phenotypic variance into sire and dam components, and thus estimates of *V_A_* and *V_D_*. To make the decomposition non-trivial, the quantitative trait is given a non-additive genetic architecture: 100 di-allelic loci with random per-locus additive effects drawn from an exponential distribution (rexp) and random dominance coefficients drawn from a normal distribution (rnorm) centred on zero. See the Crossing design subsection in the main text, and Manual section 2.10.

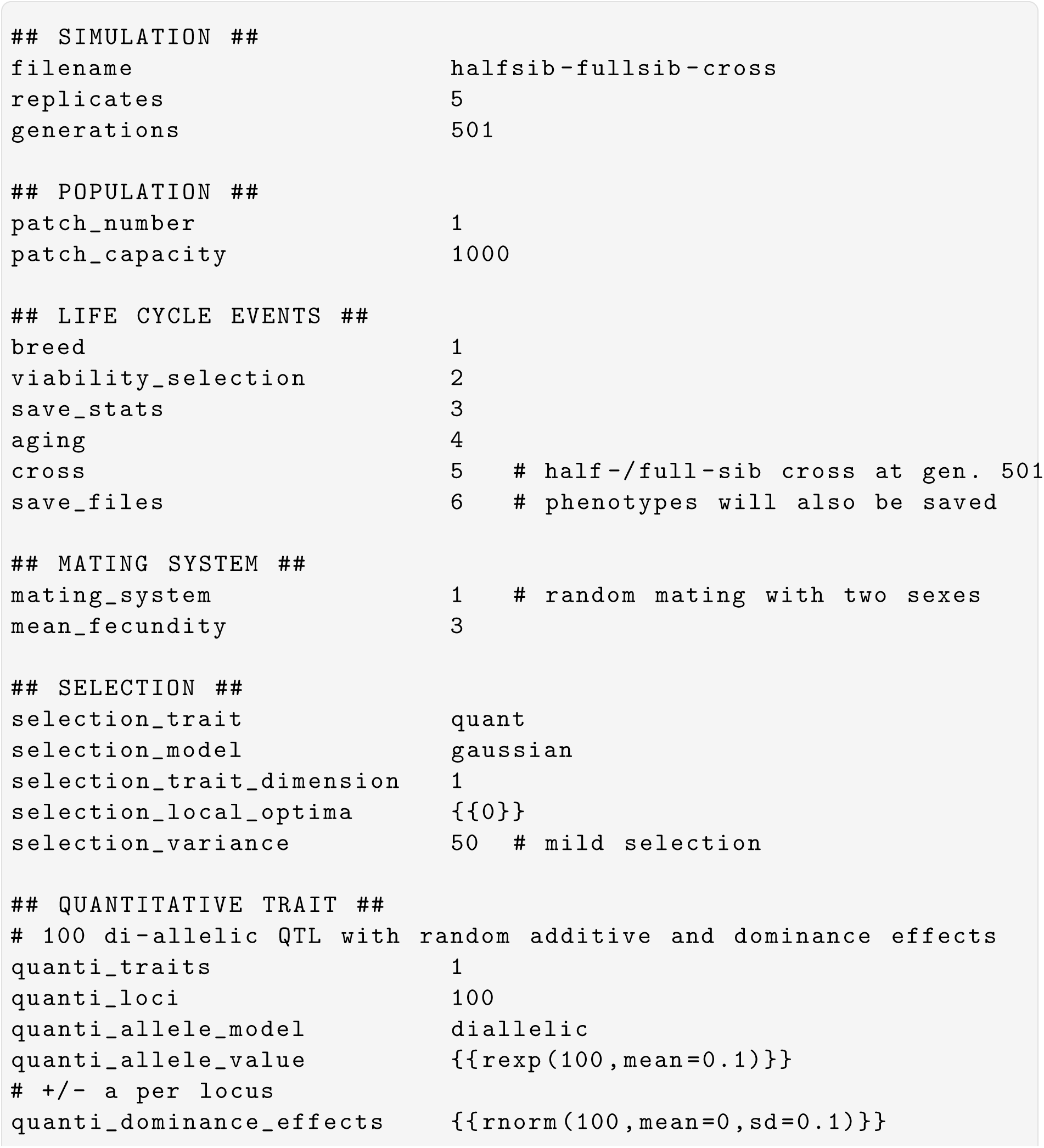
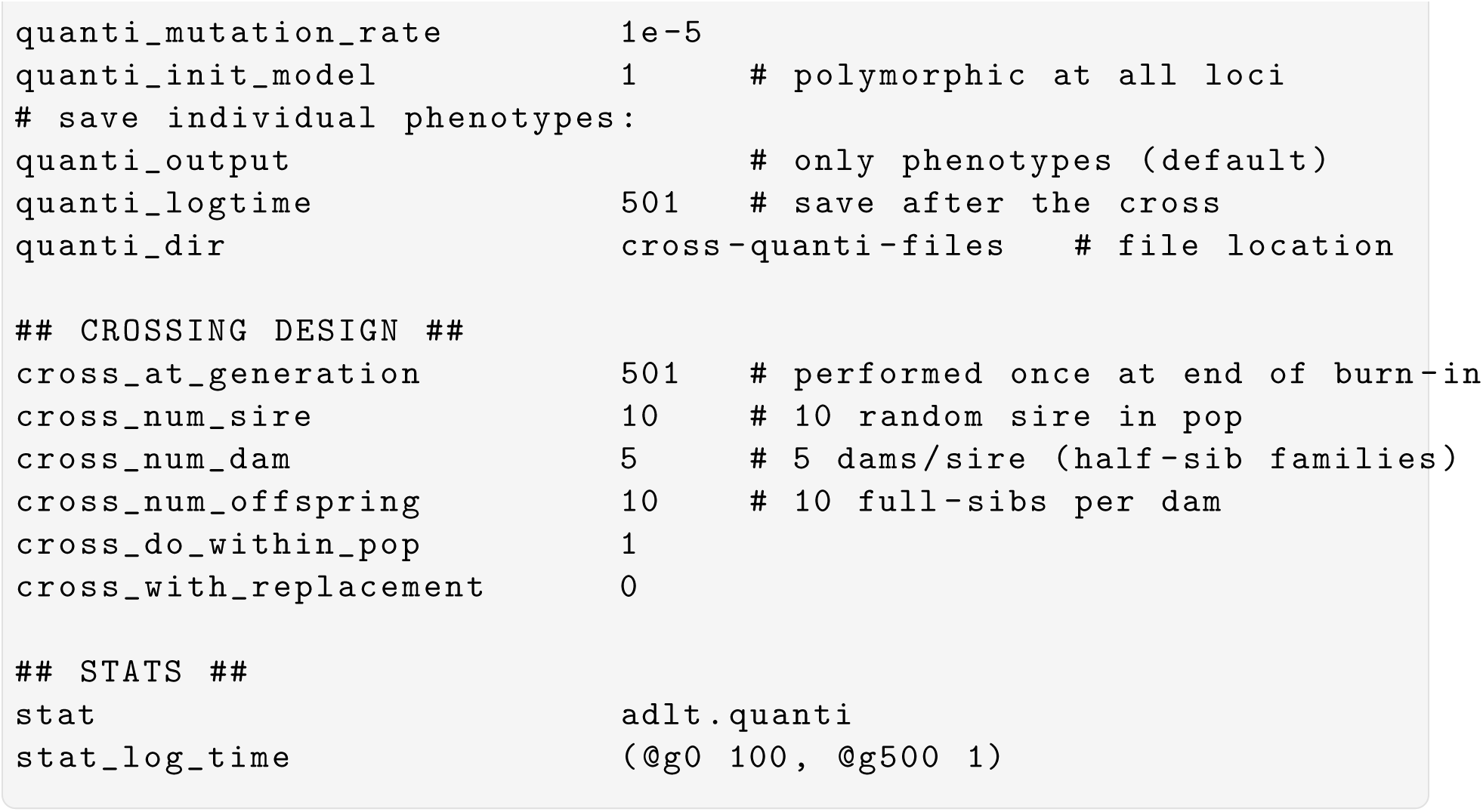

### 1.8 Crossing: full diallel

The cross LCE allows for far more than half-/full-sib matings thanks to the possibility to pass a pedigree file. The file can be used to design the exact kind of crosses one wants to implement and use it to create more than just an F1. In fact, Nemo 2.4 accepts pedigrees of any depth and size. For instance, the pedigree of a monitored population can be passed to model trait inheritance. This example is for a full diallel cross where each of 5 progenitors is crossed with every other progenitor, including itself. This simple example generates one F1 per mating.

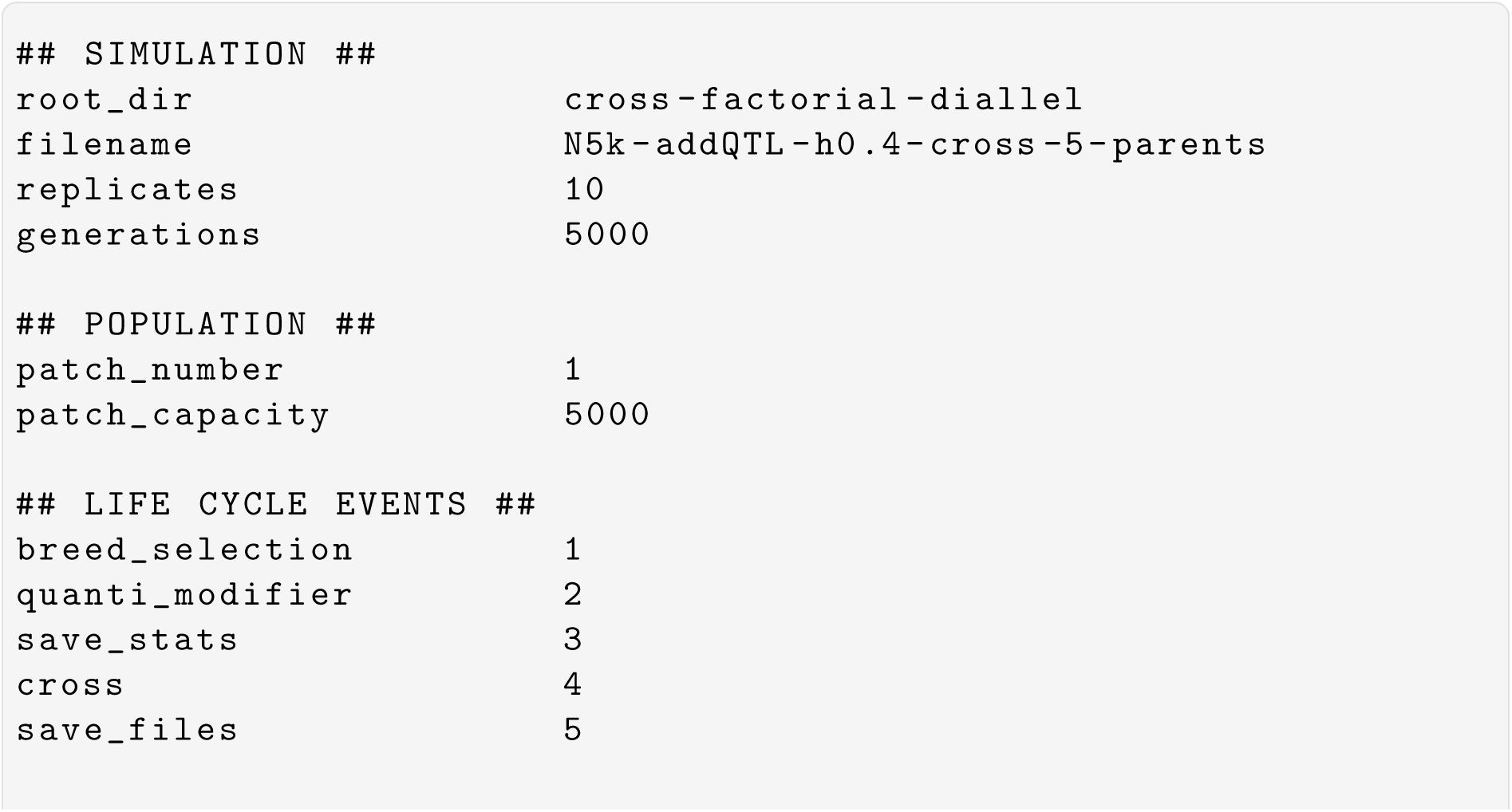
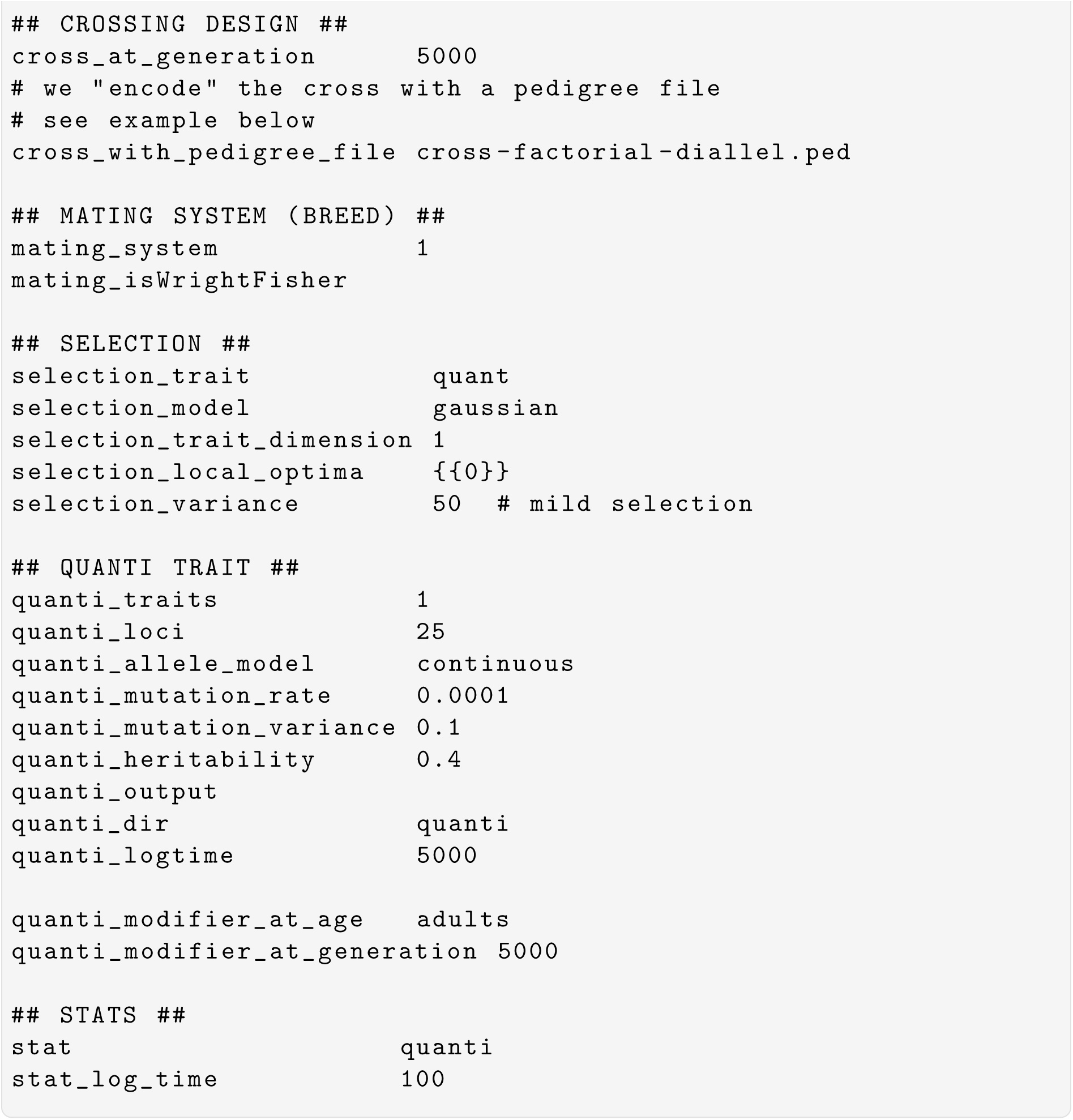

The pedigree file looks like shown below, where the first column is the individual ID, and the second and third are the IDs of the father and mother of that individual. Parents marked “NA” will be randomly drawn from the simulated population at generation 5000 to generate the five progenitors of the cross (individuals 1-5).

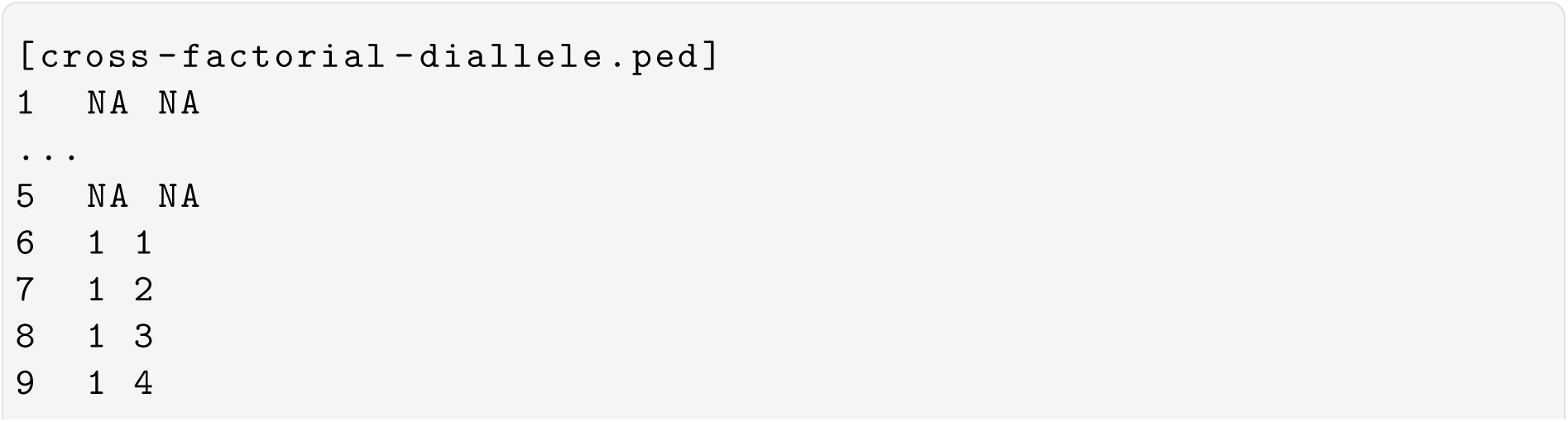
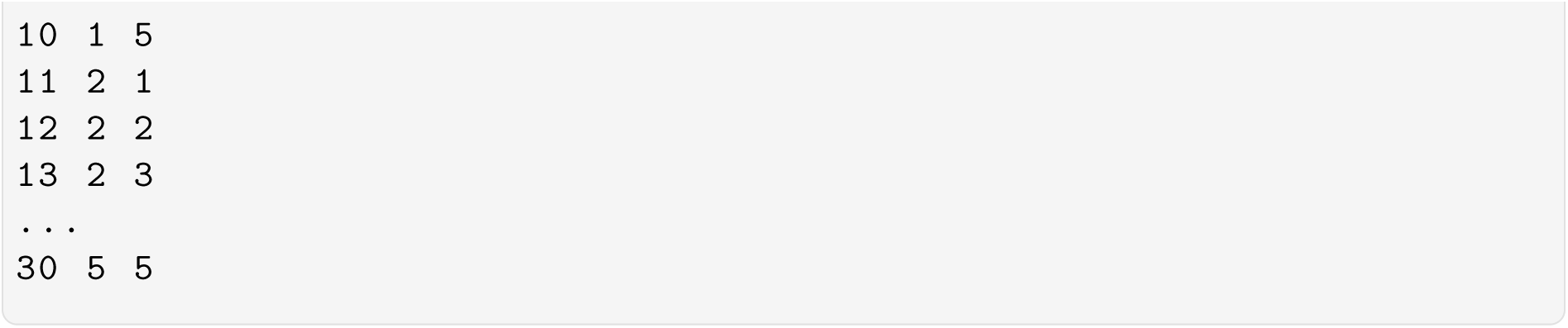

The .quanti output files will look something like below with the individual IDs as in the .ped file and the IDs of the parents of the 5 progenitors are those of the individuals from the simulated population. Note the difference between the phenotype (P1) and genotype (G1) values of the quantitative trait caused by heritability *h*^2^ = 0.4:

| pop | P1 | G1 | ... | father | mother | ID |
| --- | --- | --- | --- | --- | --- | --- |
| 1 | 1.386 | 0.5212 | ... | 2253 | 3862 | 1 |
| 1 | -0.1179 | -0.6365 | ... | 1470 | 4366 | 2 |
| 1 | -1.01 | -0.3717 | ... | 1452 | 3089 | 3 |
| 1 | 0.05319 | -0.1202 | ... | 1529 | 4125 | 4 |
| 1 | -1.485 | -1.299 | ... | 1281 | 3762 | 5 |
| 1 | 0.7711 | 0.6505 | ... | 1 | 1 | 6 |
| 1 | 0.02067 | -0.0999 | ... | 1 | 2 | 7 |
| 1 | 0.3113 | -0.4513 | ... | 1 | 3 | 8 |
| 1 | -0.0149 | 0.1225 | ... | 1 | 4 | 9 |
| 1 | -0.0065 | 0.01666 | ... | 1 | 5 | 10 |
| 1 | 0.6144 | 0.4789 | ... | 2 | 1 | 11 |
| 1 | -0.7534 | -0.6652 | ... | 2 | 2 | 12 |
| 1 | -1.661 | -0.3324 | ... | 2 | 3 | 13 |
| 1 | -0.3762 | -0.288 | ... | 2 | 4 | 14 |
| 1 | -2.716 | -1.526 | ... | 2 | 5 | 15 |
| ... |  |  |  |  |  |  |

### 1.9 Running and storing a burn-in simulation

The store LCE lets us save the whole population to a binary file which can later be reloaded at the start of a simulation and used to seed the first generation. The binary files are automatically compressed, with one file per replicate, saved at a given generation. All files are moved to a .tar archive at the end of the simulation. This default behaviour can be changed, for instance to prevent the default archiving of the files.

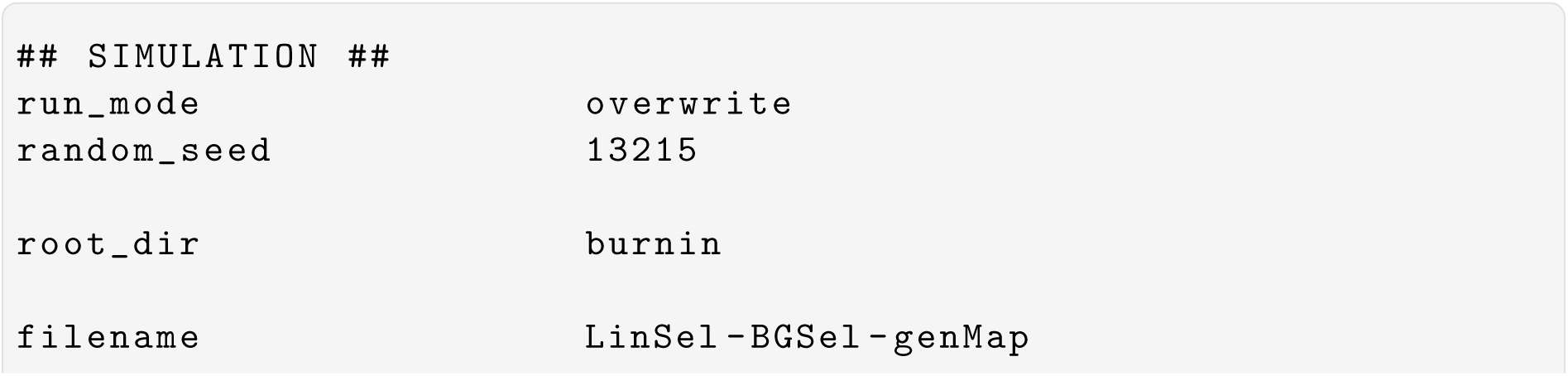
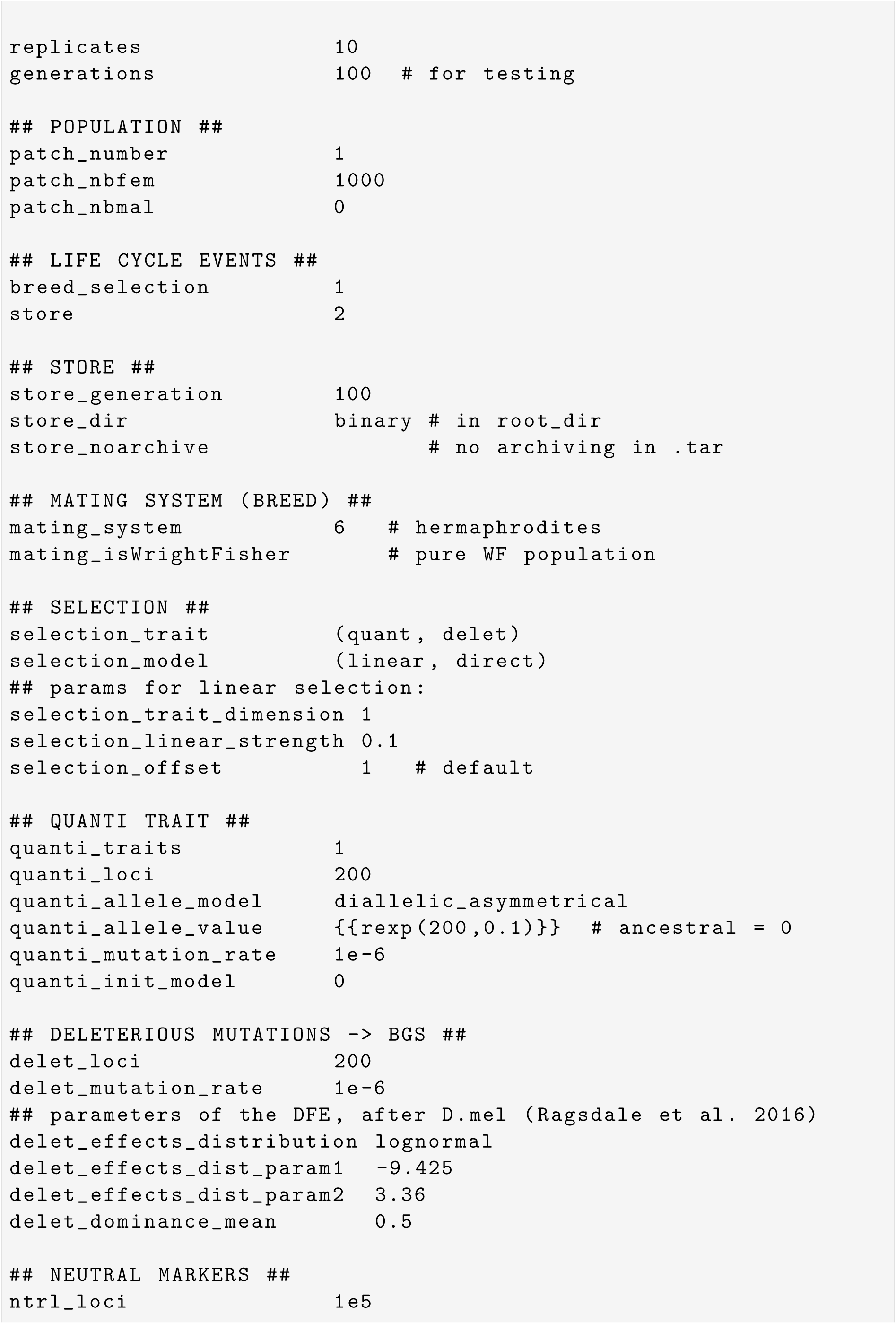
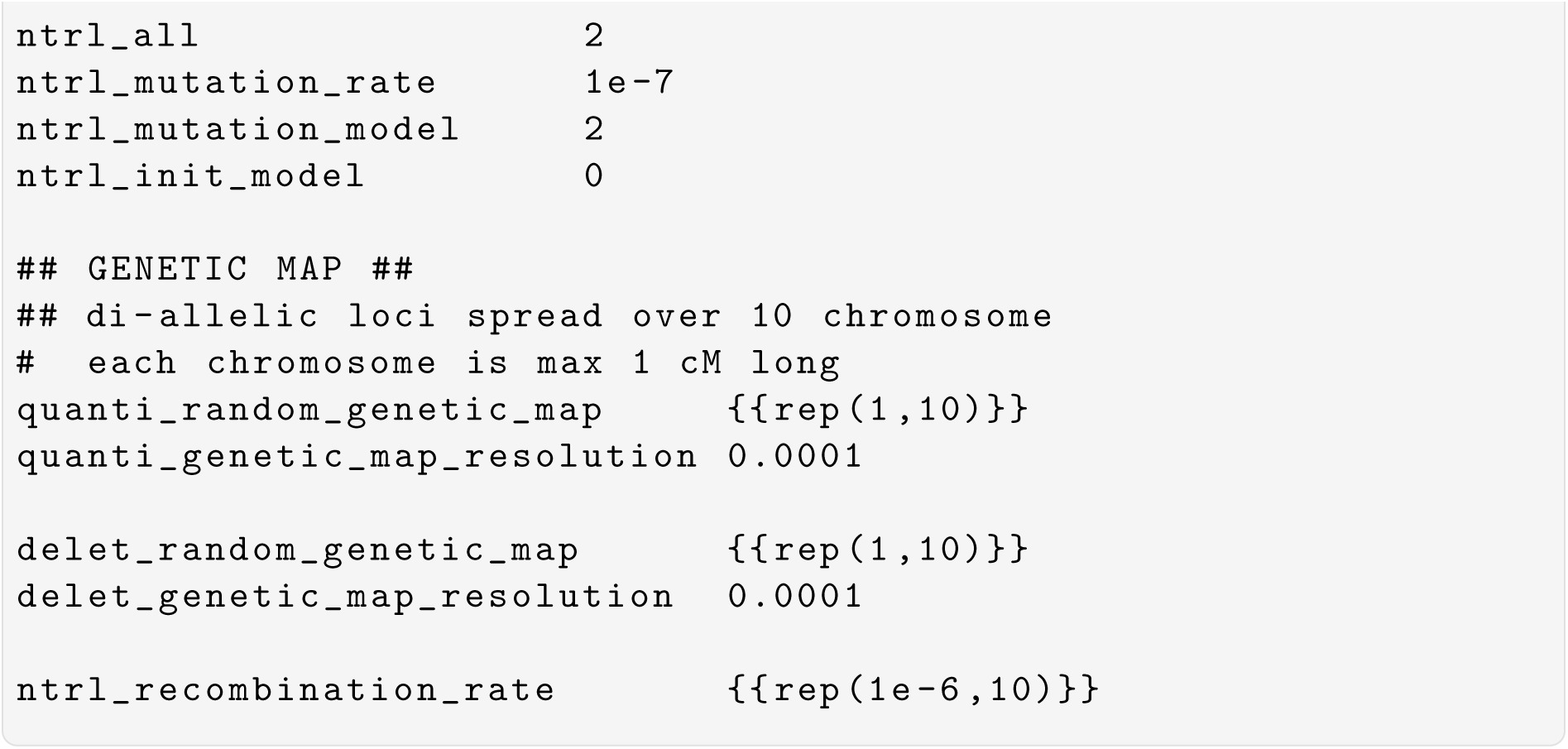

### 1.10 Load a stored population and save sampled data to geno-type files

In this example, we load the previously run burnin simulation to only compute statistics and sample individuals from the loaded population. The life cycle thus has only two LCEs, `save_stats` to compute the statistics and `save_files` to save the genetic information of a sample of 125 adults in the population. Since the two traits modeled, quant and delet are diallelelic traits, the SNP format can be used to save the data in SNP genotype format. We also trim the loci to minimum allele frequency (MAF) of 0.01 in order to include the locus genotype information to the output file. One file per replicate is then written to disk in the root directory specified in the configuration file shown below.

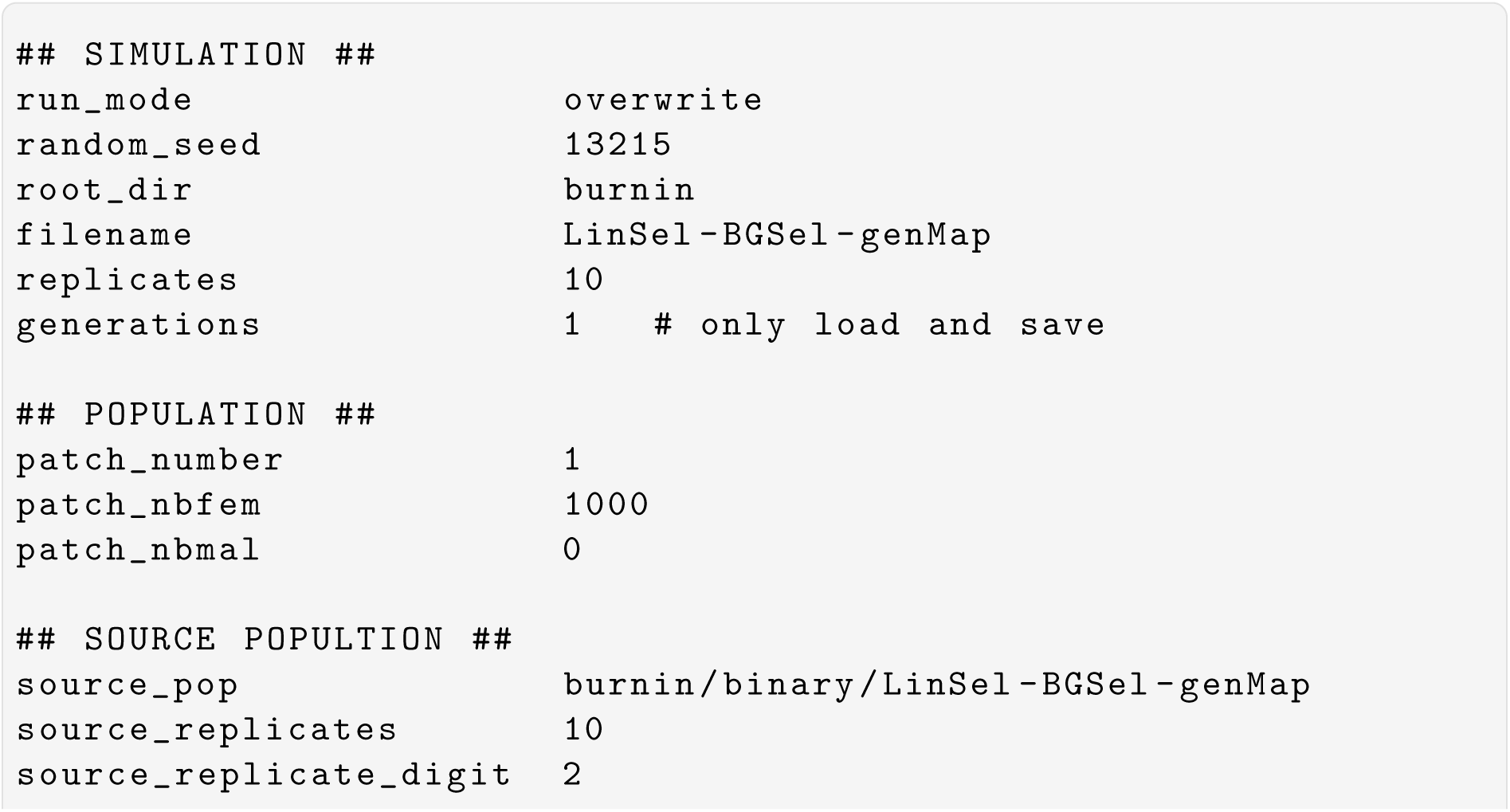
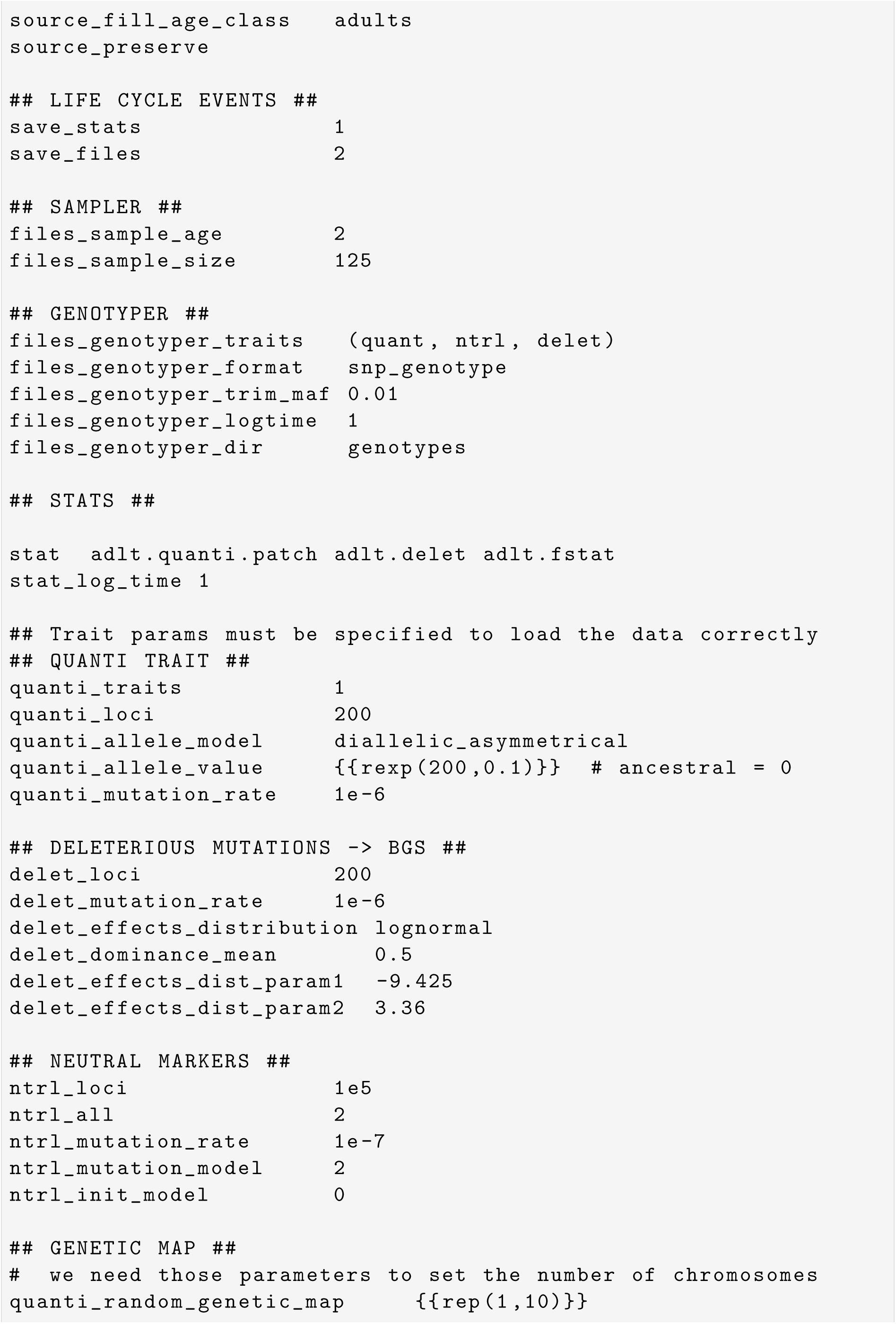
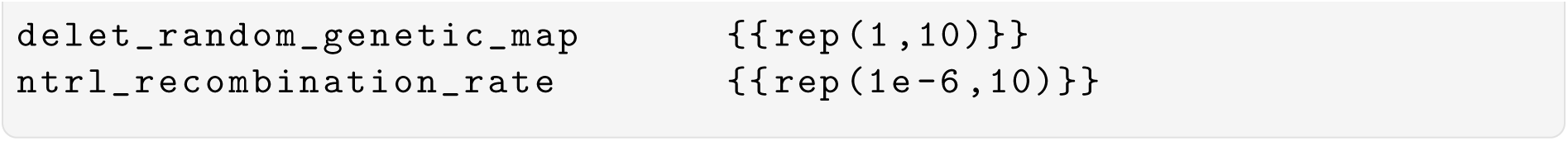

### 1.11 Encoding a large sequence of simulations in the configuration file with parameter substitution and recall

Many simulations can be encoded in a configuration file when parameters receive multiple values. Such parameters are called *sequential parameters* in Nemo’s jargon. The mini language we developed in Nemo allows us to then recall the parameter values using a substiution string with simple syntax. The substitution string starts with ‘%‘ symbol followed by the number of the sequential parameter: %4 refers to the fourth sequential parameter found in the file. The parameters are ordered in alphabetical order. To facilitate parameter recall and value assignment, it is possible to define dummy parameters, assign them multiple values, and recall those values for the relevant parameters in the file (see below). Furthermore, it is possible to link the values of two or more sequential parameters when they need to vary in sync, instead of generating the complete set of combinations of those values. Linked sequential parameter values are encoded in square brackets: param_1 [v1 v2 v3] and param_2 [a b c]. This setting will generate 3 simulations, while the same parametrization without the bracketing would generate 3 × 3 = 9 simulations.

In addition, it is possible to save parameter values to separate files and provide the path to the file as argument value when using the ‘&’ symbol. The example below uses this feature to separately store very large dispersal matrices, environmental maps (values of trait optima) and genetic maps with *>* 10^5^ values.

This example runs a model of local adaptation in a spatially varying environment on a 48 × 48 patch grid with gene flow and background selection. Individuals carry a single ∼10cM chromosome with 100 QTN, 200 deleterious sites, and ∼200,000 neutral SNPs. Three different map settings are modelled, encoded with the `1_map_res` and `2_map_len` dummy paramteres and their linked-sequetial values. The maps vary in size and number of SNPs, and QTN and deleterious mutation effects, which a redrawn for each new simulation. Three other parameters vary as well, in full combinatorial mode, generating 3 × 3 × 10 = 90 different combinations. The total number of simulations is then 3 × 90 = 270. Each simulation receives a different output filename of the form:

`bgs_chr1_V0.5_c1_env1`

`bgs_chr1_V0.5_c1_env2`

[…]

`bgs_chr3_V2_c2_env10`.

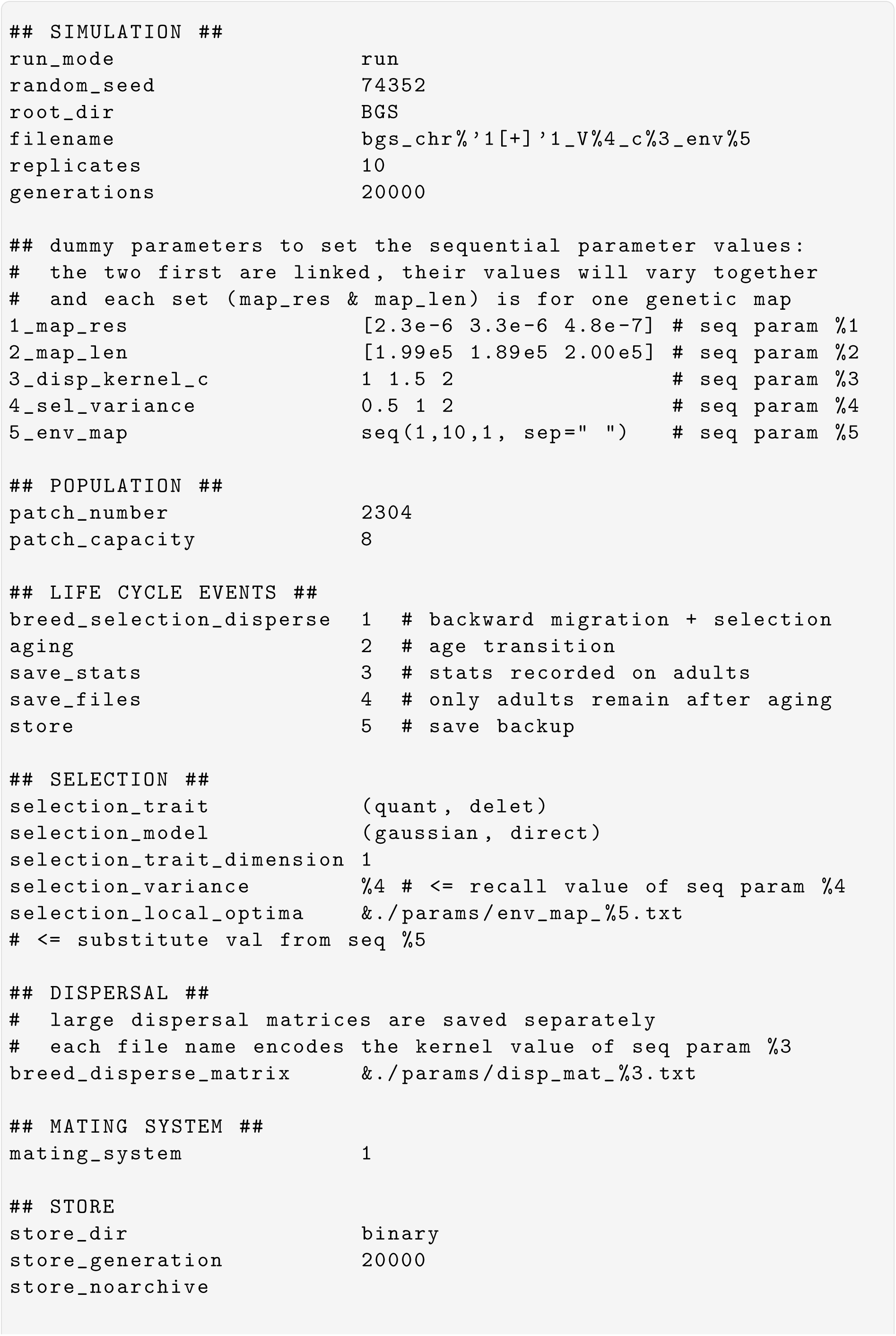
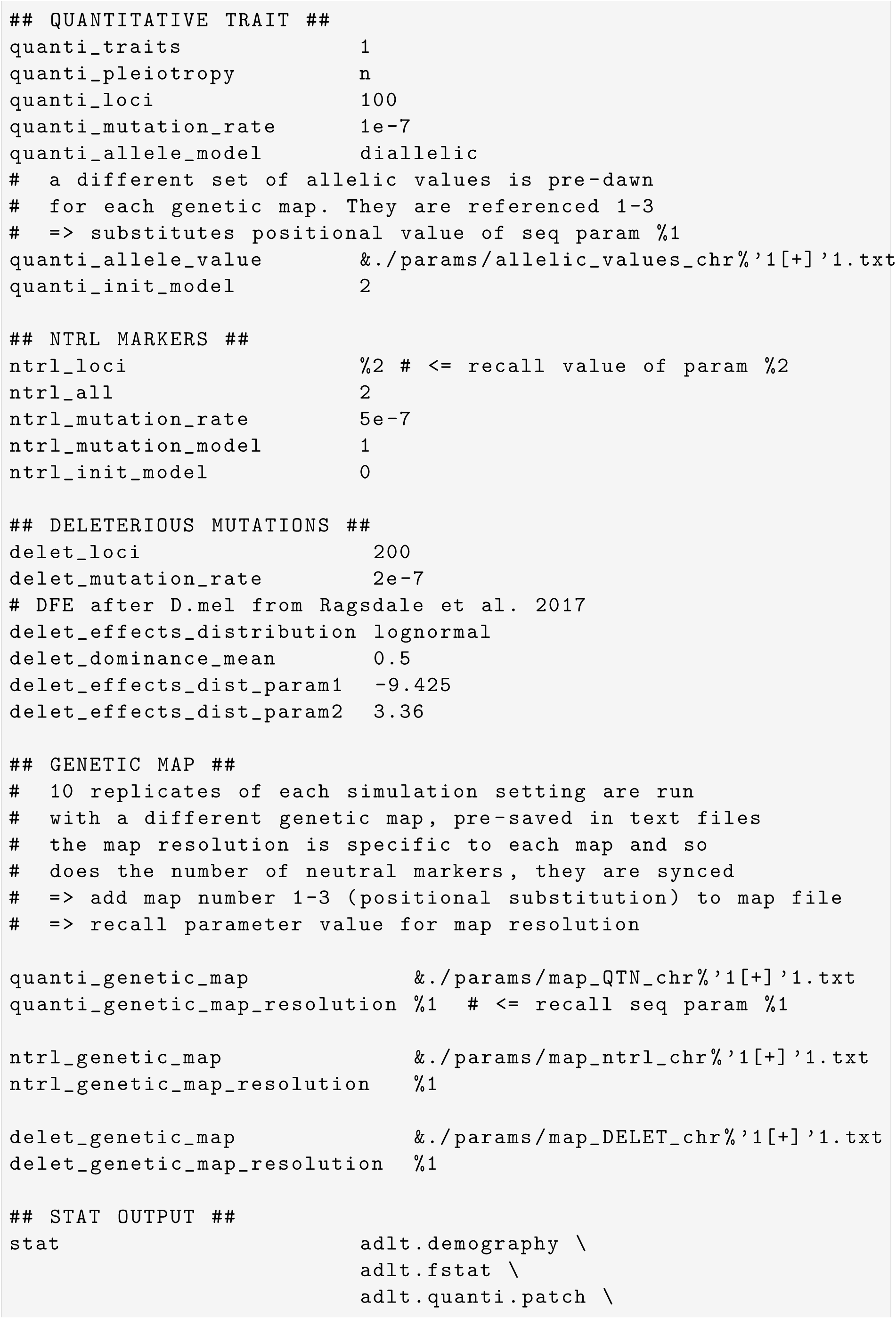
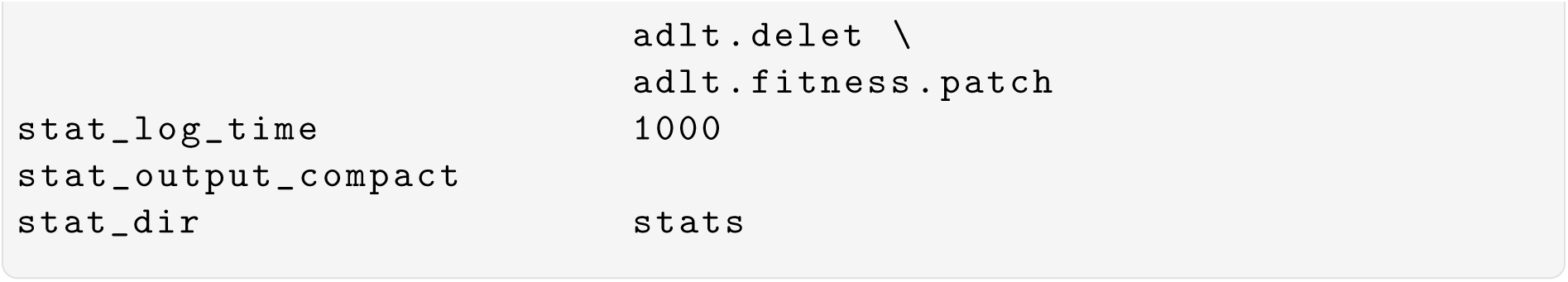

## 2 Appendix

This section collects the scripts used to run the software comparison benchmark. The parameter files for Nemo, quantiNemo, and SLiM scripts are pasted here. The parameters used in the equilibrium simulations are a subset of the simulations below, run for 30,000 generations/ticks instead of 1000.

### 2.1 Nemo 2.4

#### 2.1.1 QTL simulation

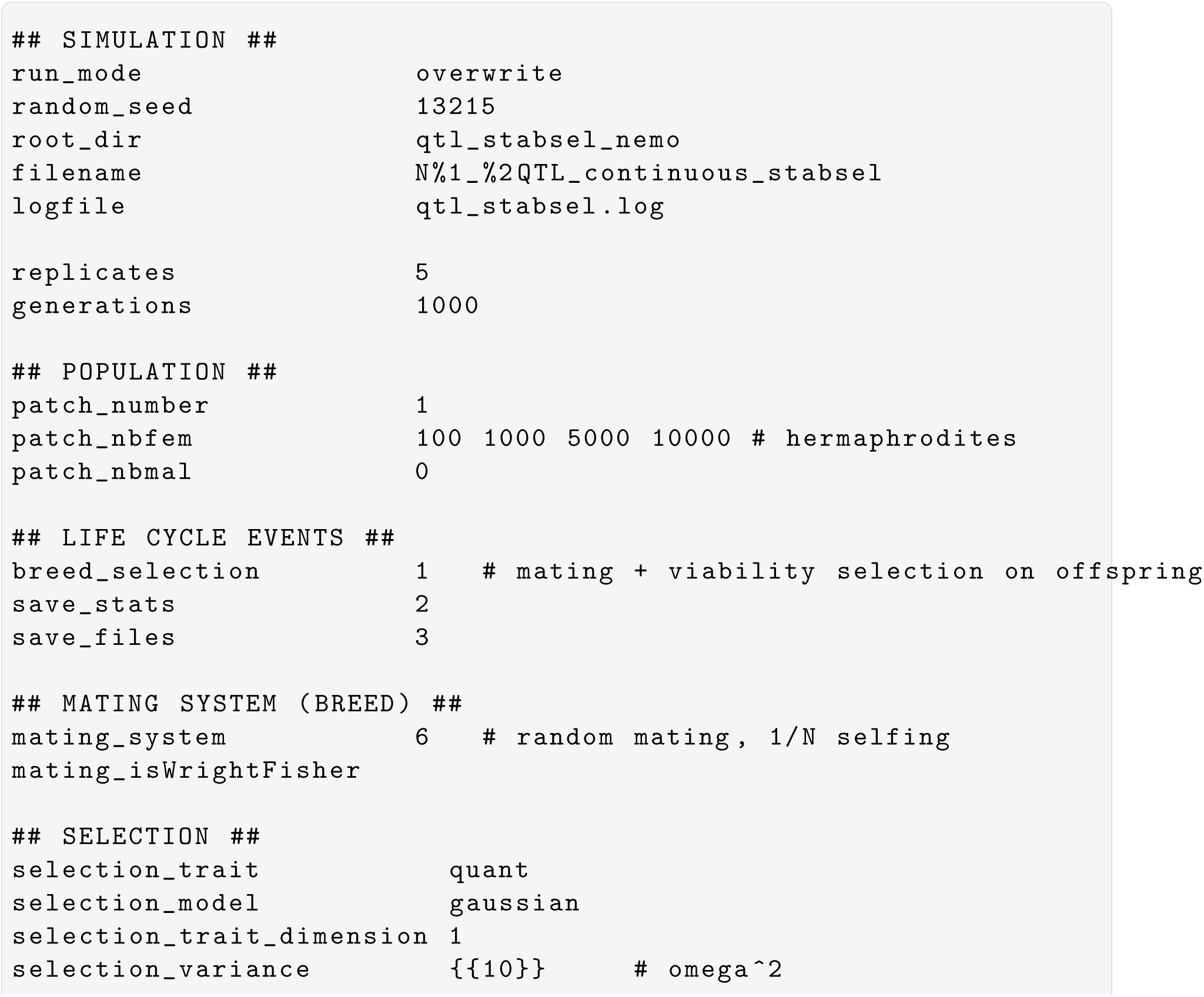
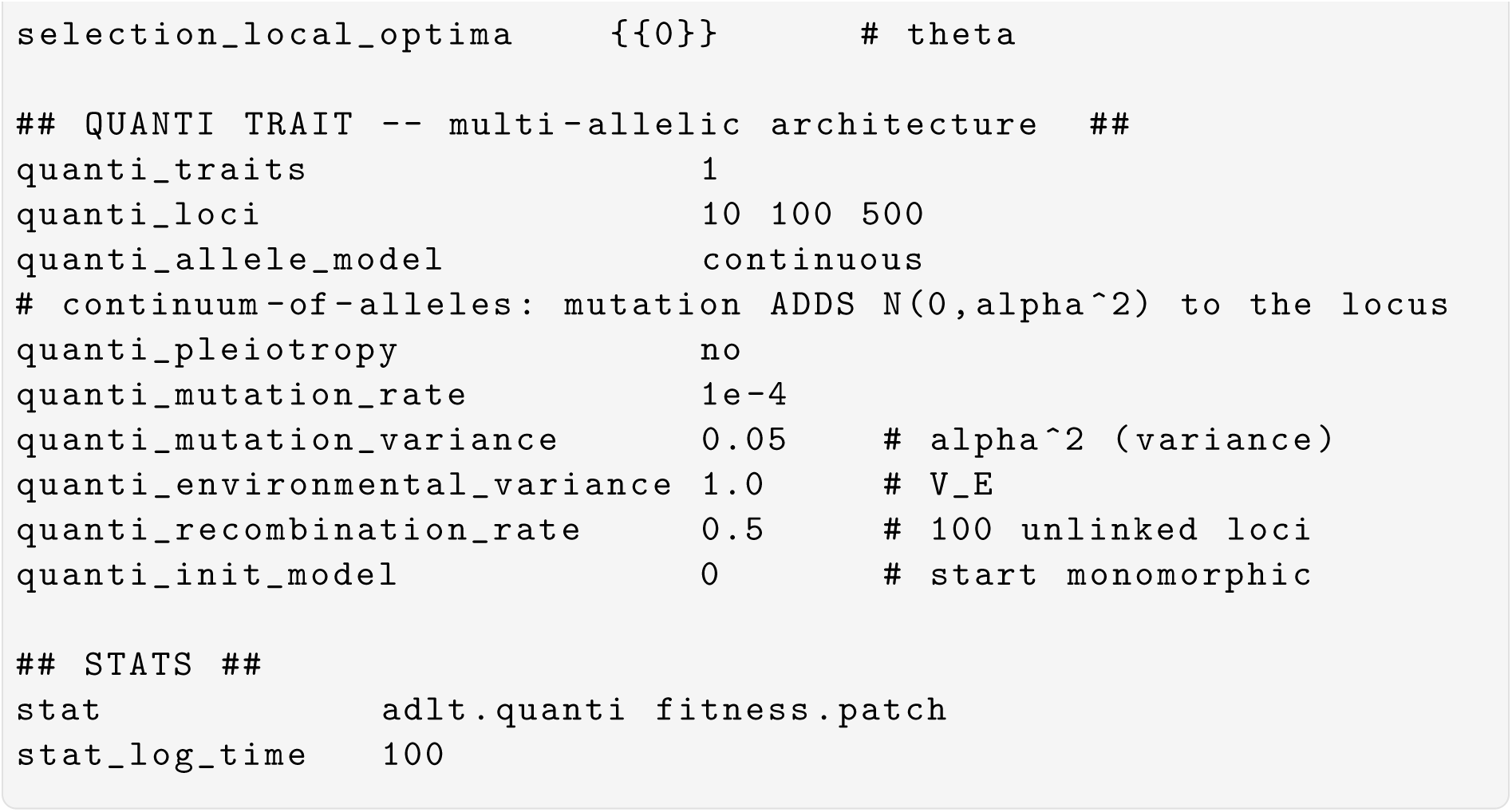

#### 2.2.2 QTN simulation

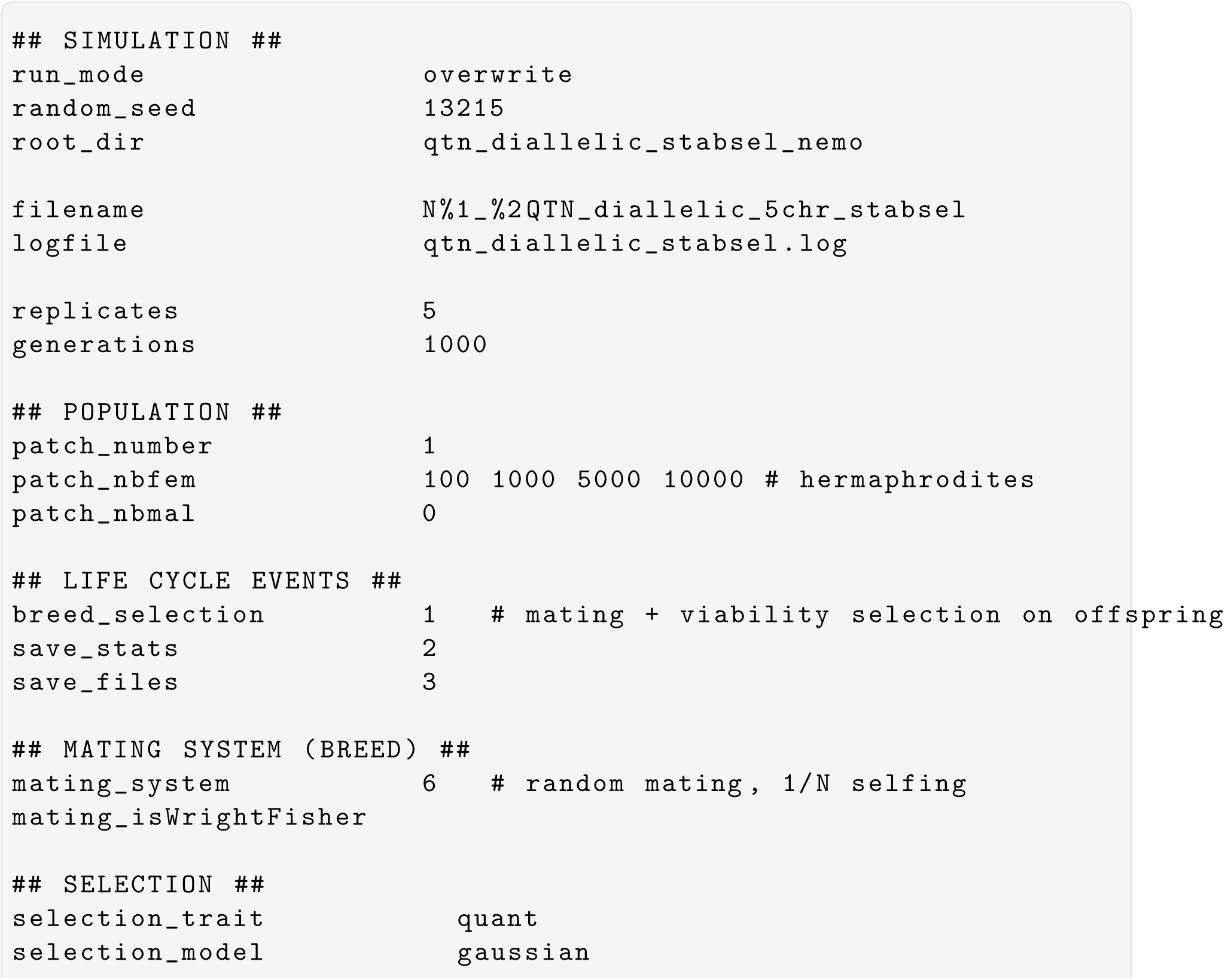
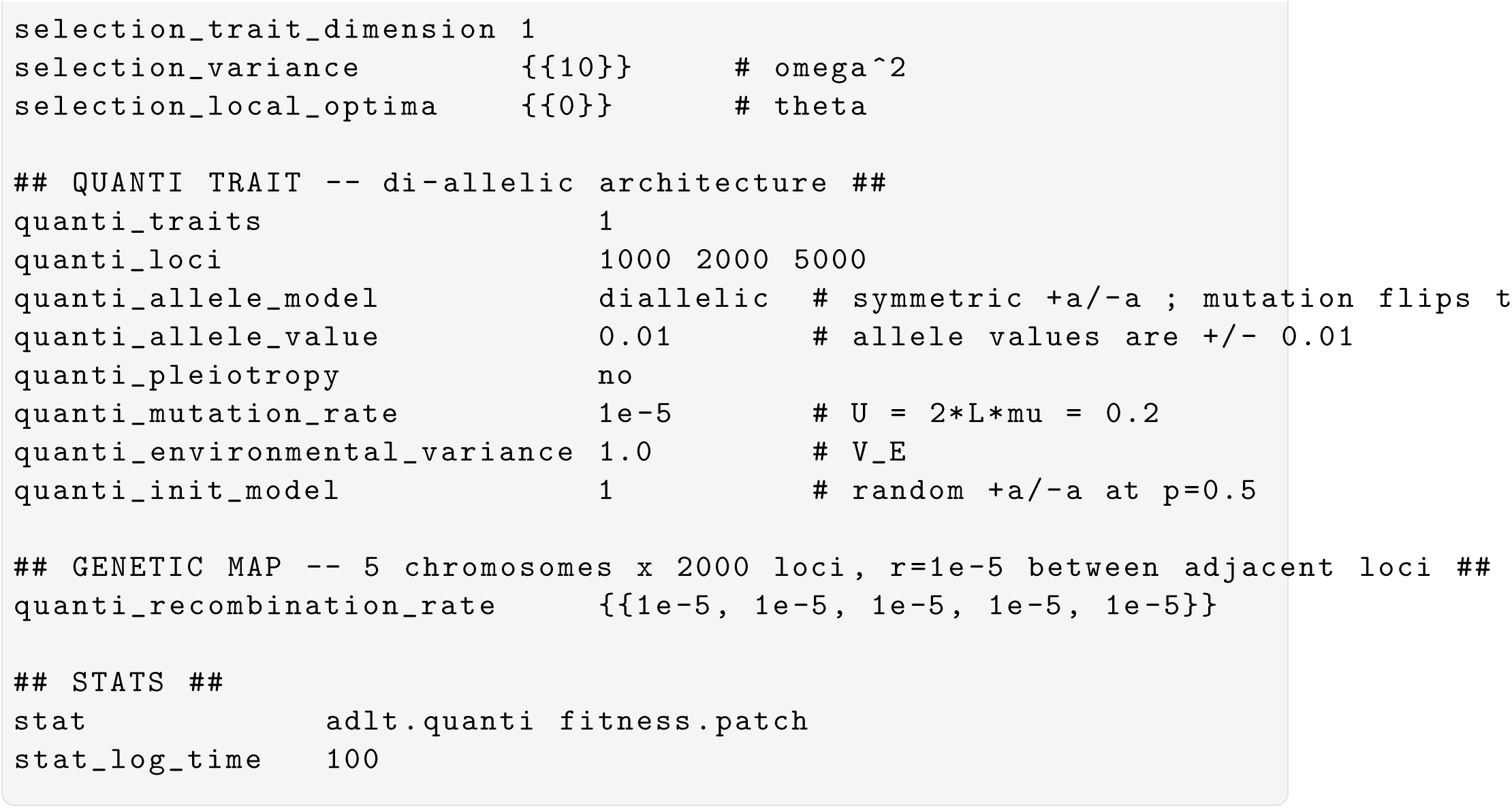

### 2.2 quantiNemo 2.0

Simulation parameters for quantiNemo2. The software does not have the sequential parameter machinery implemented in Nemo. Parameter values provided in the parameter file can be overridden on the command line, as for instance:

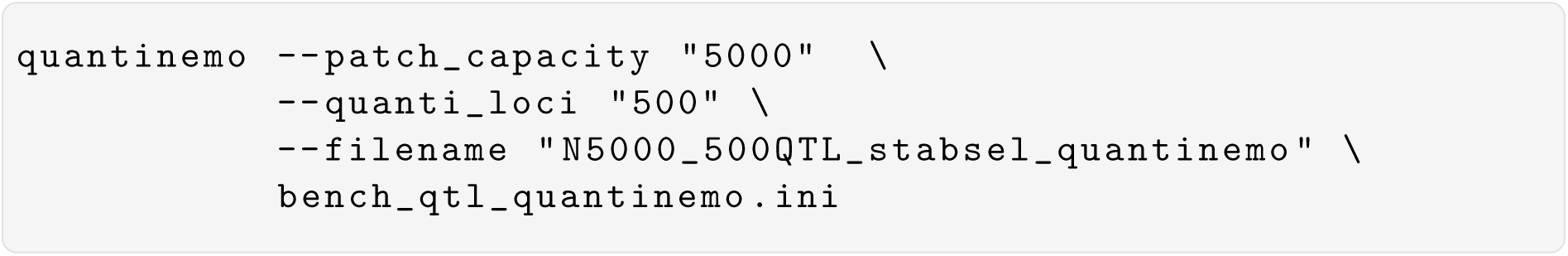

The file bench_qtl_quantinemo.ini contains the parameters.

#### 2.2.1 QTL simulation

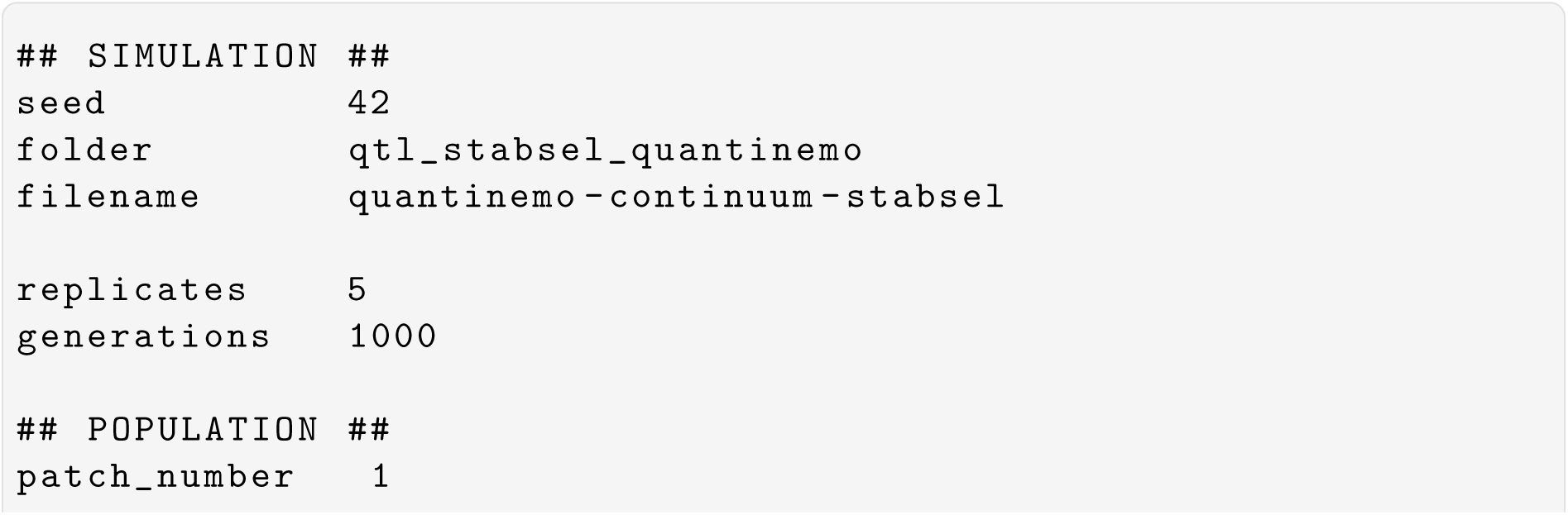
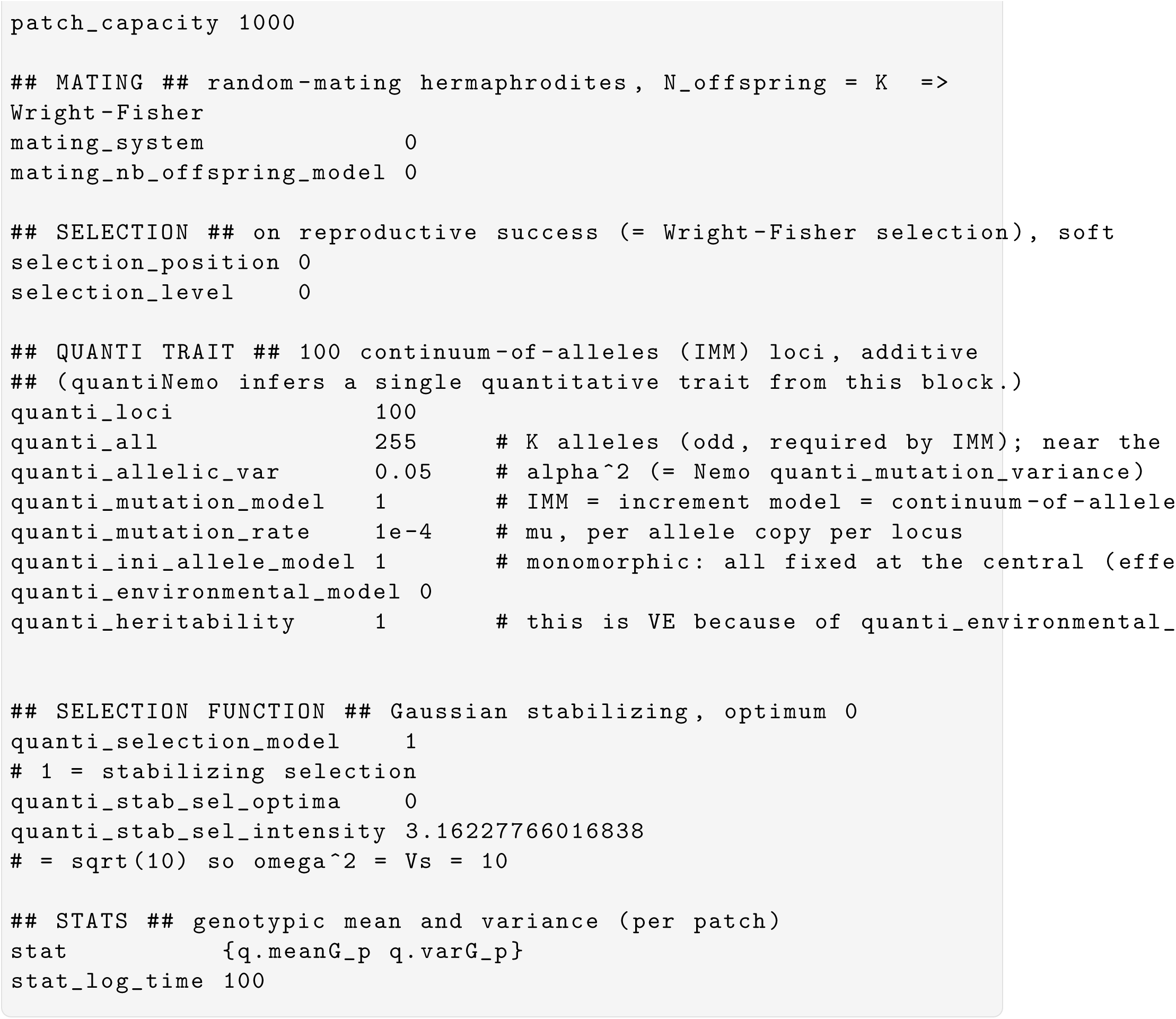

#### 2.2.2 QTN simulations

quantiNemo doesn’t allow setting a chromosome-wide recombination rate. Instead, the map position and index of each locus must be specified explicitly. For large number of loci, as here, the positions and indices are written to separate files and linked in the parameter file. These external files cannot be specified on the command line and must be set in the parameter file. There would thus be three different .ini file, one for each number of QTN.

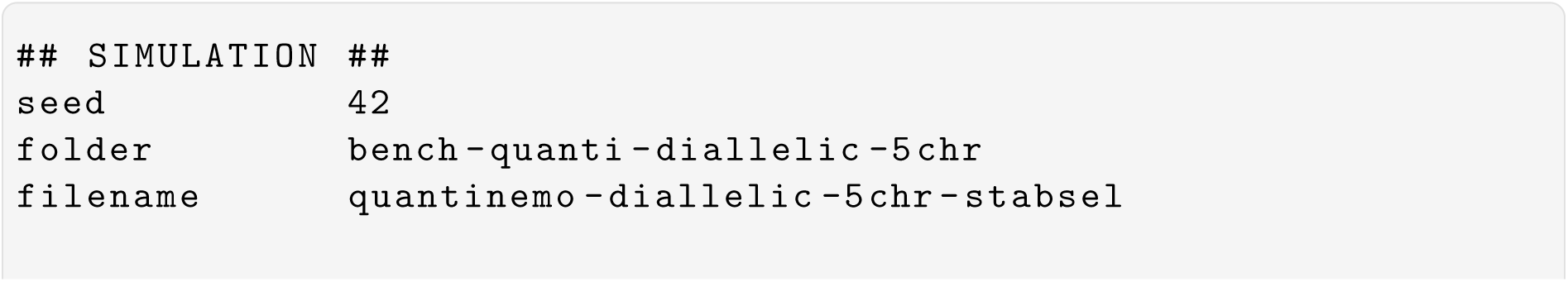
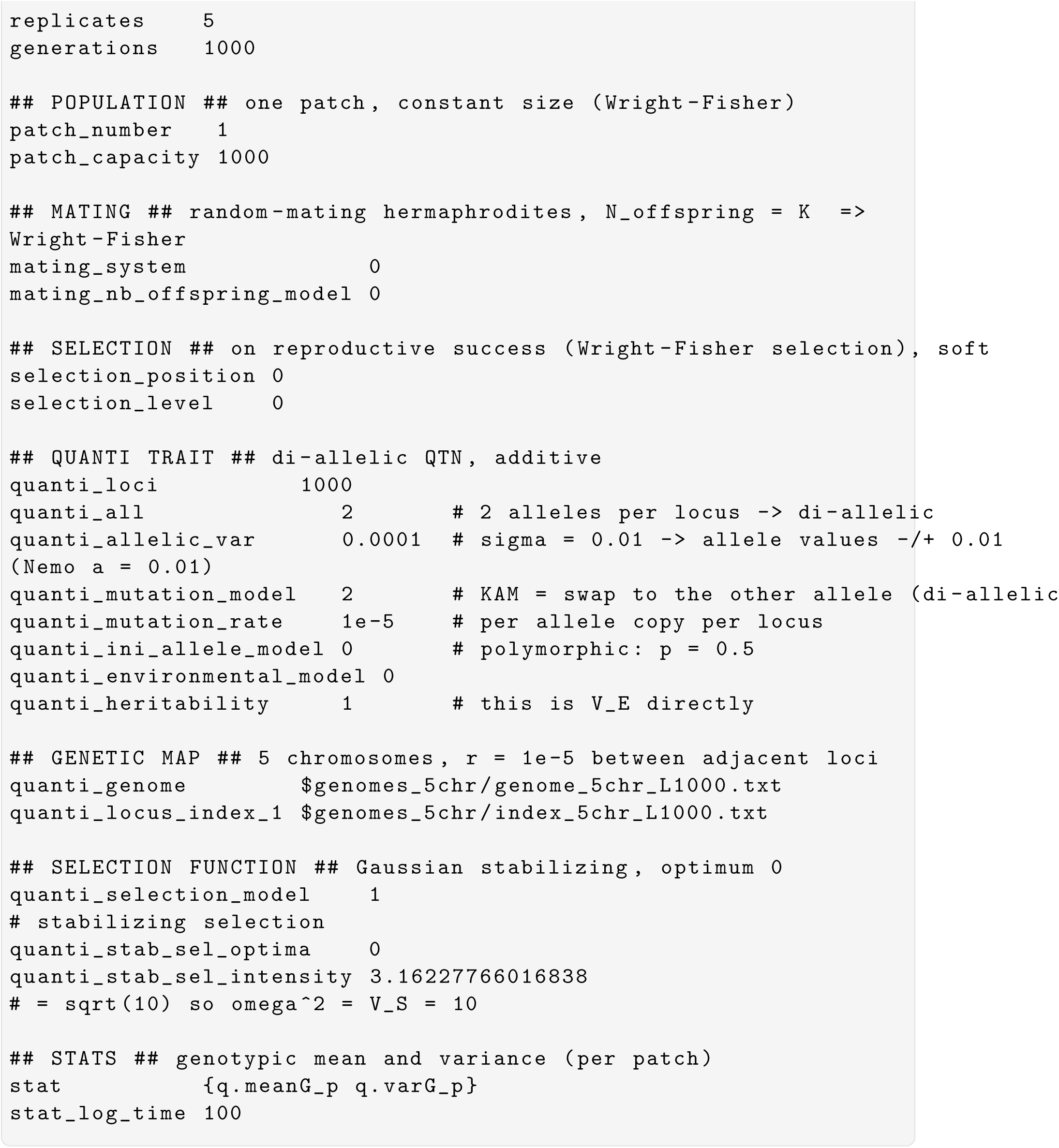

### 2.3 SLiM 5.2

SLiM parameters are specified in a .slim script. Parameters can be overridden on the command line with ‘-d param=…’, which allows for running simulations over a parameter grid within a bash script.

#### 2.3.1 QTL simulations

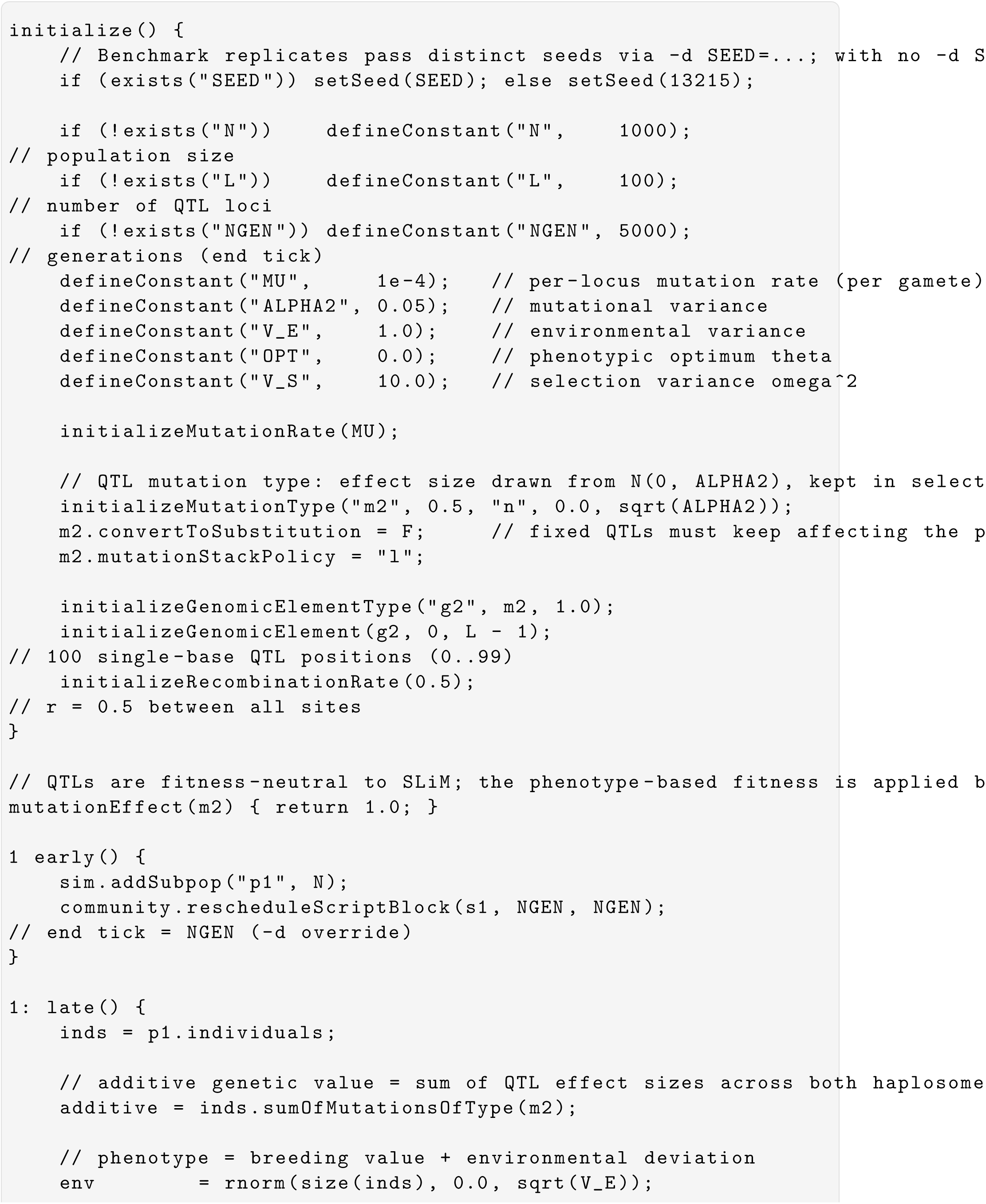
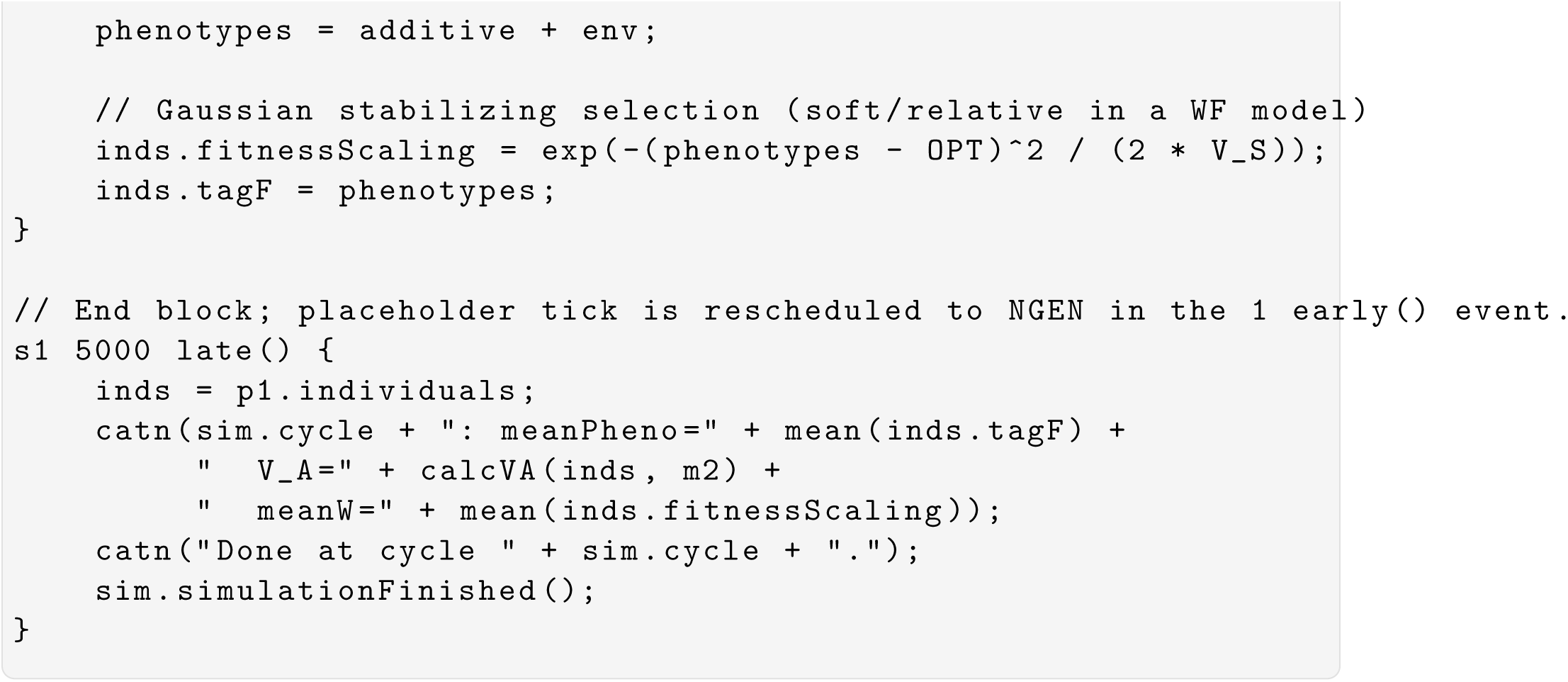

#### 2.3.2 QTN simulations

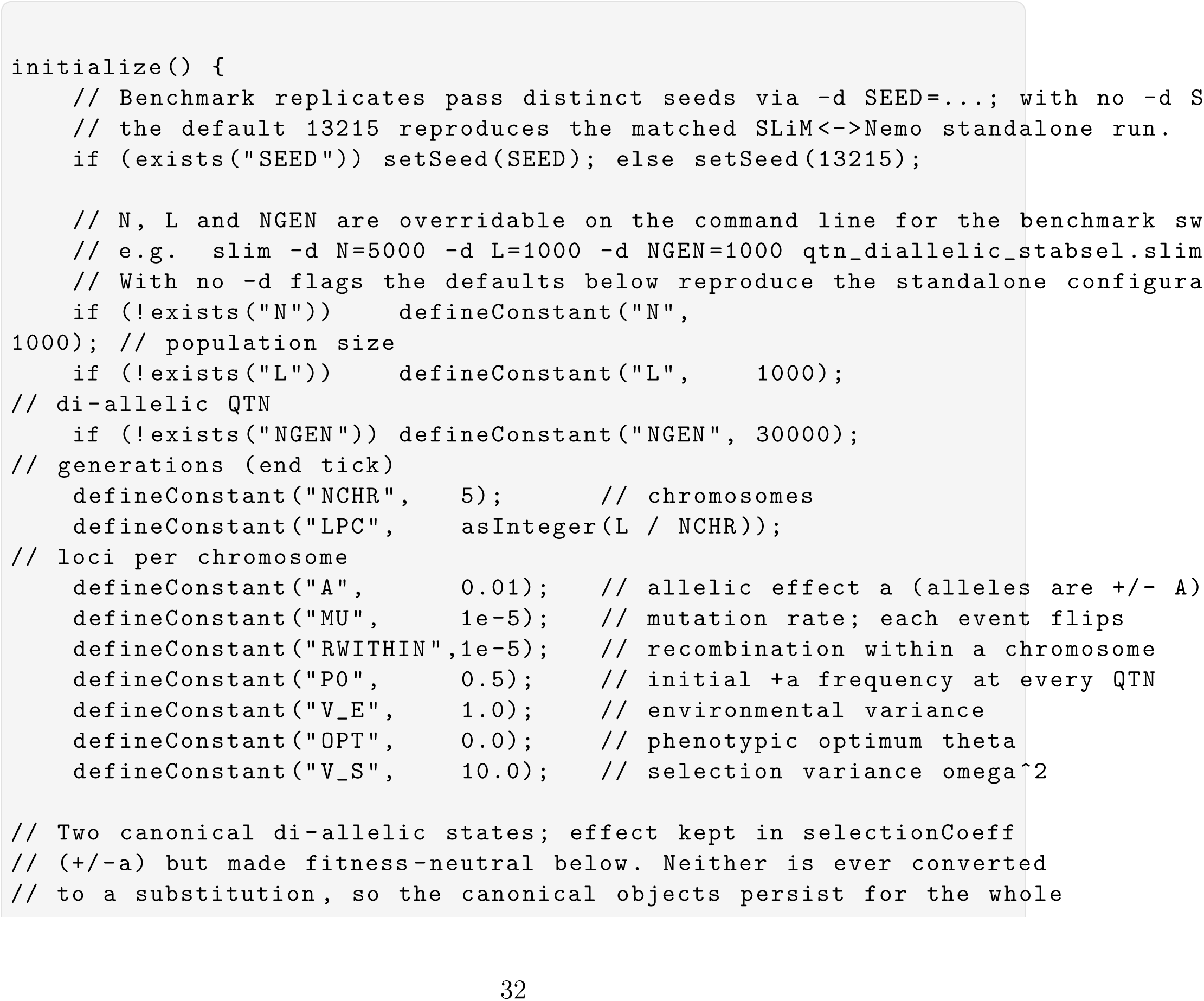
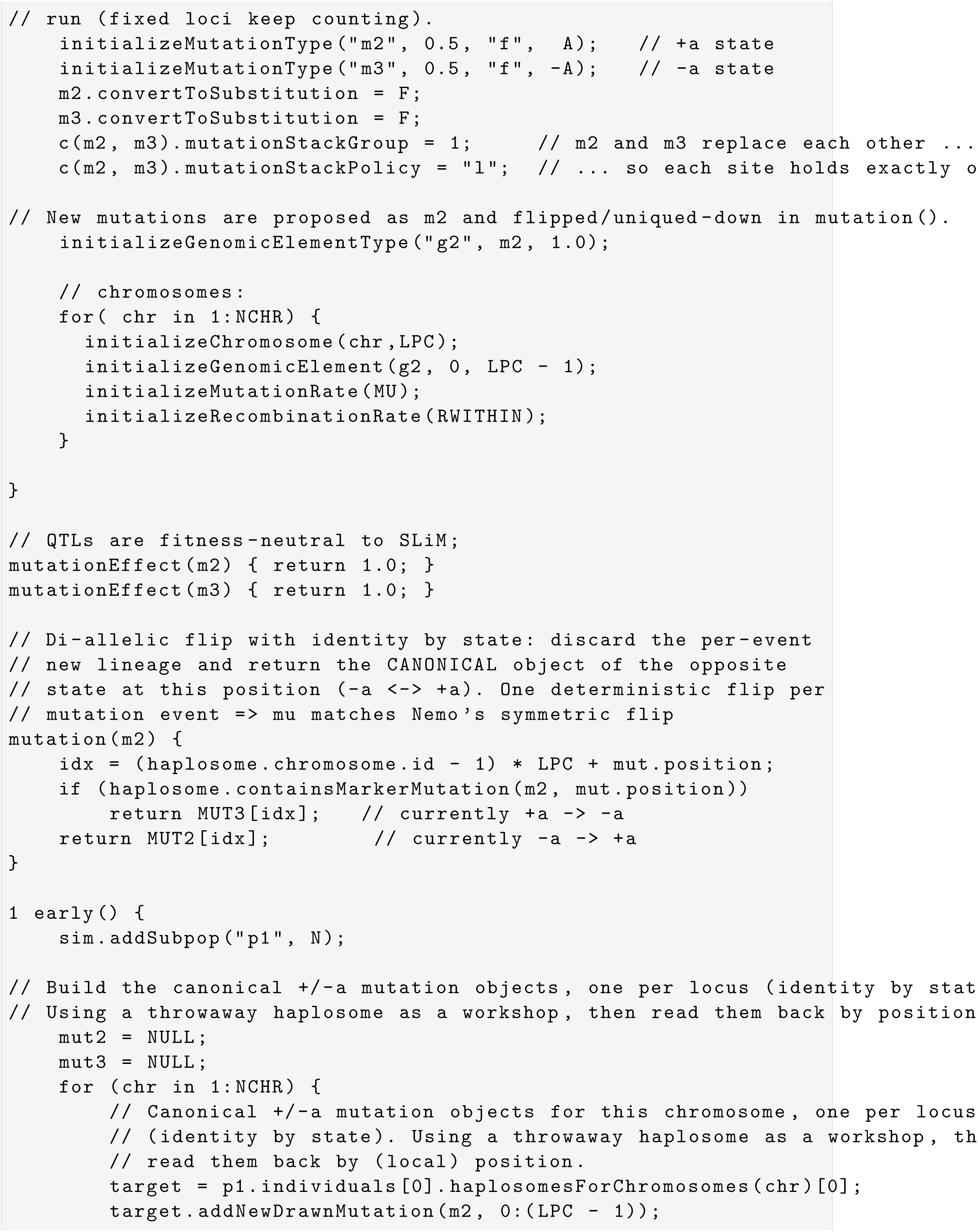
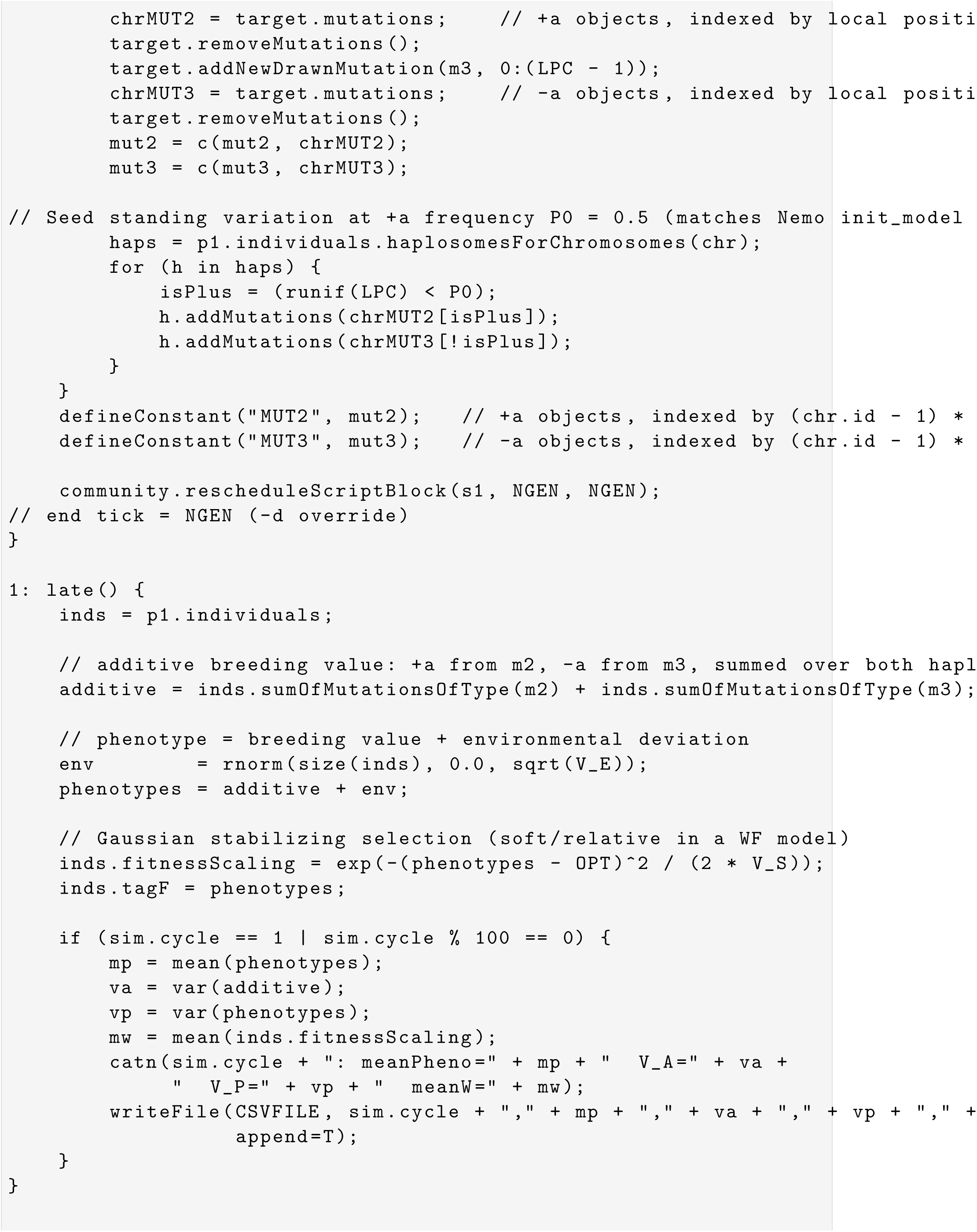
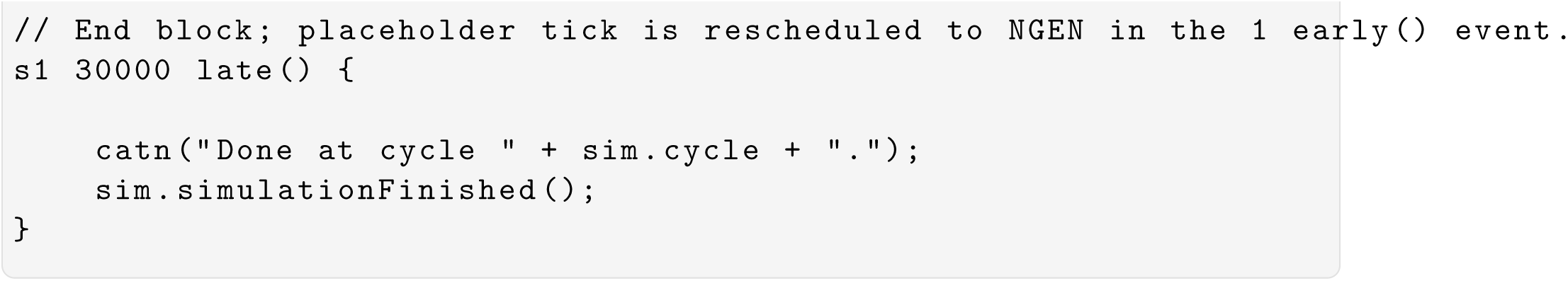

## References

Bürger, R. (2000). The mathematical theory of selection, recombination, and mutation. In Levin, S., editor, Wiley series in Mathematical and Computational Biology. Wiley & Sons, Ltd, UK.

Burger, R., Wagner, G. P., and Stettinger, F. (1989). How much heritable variation can be maintained in finite populations by mutation-selection balance? Evolution, 43(8):1748–1766.

Carter, A. J. R., Hermisson, J., and Hansen, T. F. (2005). The role of epistatic gene interactions in the response to selection and the evolution of evolvability. Theor. Popul. Biol., 68(3):179–196.

Chang, C. C., Chow, C. C., Tellier, L. C., Vattikuti, S., Purcell, S. M., and Lee, J. J. (2015). Second-generation PLINK: rising to the challenge of larger and richer datasets. GigaScience, 4(1):s13742–015–0047–8.

Chebib, J. and Guillaume, F. (2017). What affects the predictability of evolutionary constraints using a G-matrix? The relative effects of modular pleiotropy and mutational correlation. Evolution, 71(10):2298–2312.

Chebib, J. and Guillaume, F. (2021). Pleiotropy or linkage? Their relative contributions to the genetic correlation of quantitative traits and detection by multitrait GWA studies. Genetics, 219(4):iyab159.

Cotto, O., Schmid, M., and Guillaume, F. (2020). Nemo-age: spatially explicit simulations of eco-evolutionary dynamics in stage-structured populations under changing environments. Methods Ecol Evol, 11(10):1227–1236.

Cotto, O., Wessely, J., Georges, D., Klonner, G., Schmid, M., Dullinger, S., Thuiller, W., and Guillaume, F. (2017). A dynamic eco-evolutionary model predicts slow response of alpine plants to climate warming. Nat Commun, 8:15399.

Crow, J. F. and Kimura, M. (1964). The theory of genetic loads. In Proc. XI Int. Congr. Genetics, vol. 2, volume 2, pages 495–505. Oxford: Pergamon Press.

Falconer, D. S. (1965). The inheritance of liability to certain diseases, estimated from the incidence among relatives. Annals of Human Genetics, 29(1):51–76.

Falconer, D. S. and Mackay, T. F. (1996). Introduction to Quantitative Genetics. Long-man, London, fourth edition.

Fraimout, A., Guillaume, F., Li, Z., Sillanpaää, M. J., Rastas, P., and Merilä, J. (2024). Dissecting the genetic architecture of quantitative traits using genome-wide identity-by-descent sharing. Molecular Ecology, 33(6):e17299.

Gilbert, K. J., Sharp, N. P., Angert, A. L., Conte, G. L., Draghi, J. A., Guillaume, F., Hargreaves, A. L., Matthey-Doret, R., and Whitlock, M. C. (2017). Local adaptation interacts with expansion load during range expansion: maladaptation reduces expansion load. Am. Nat., 189:368–380.

Gilbert, K. J. and Whitlock, M. C. (2015). QST–FST comparisons with unbalanced half-sib designs. Mol Ecol Resour, 15(2):262–267.

Gower, G., Pope, N. S., Rodrigues, M. F., Tittes, S., Tran, L. N., Alam, O., Cavassim, M. I. A., Fields, P. D., Haller, B. C., Huang, X., Jeffrey, B., Korfmann, K., Kyriazis, C. C., Min, J., Rebollo, I., Rehmann, C. T., Small, S. T., Smith, C. C. R., Tsambos, G., Wong, Y., Zhang, Y., Huber, C. D., Gorjanc, G., Ragsdale, A. P., Gronau, I., Gutenkunst, R. N., Kelleher, J., Lohmueller, K. E., Schrider, D. R., Ralph, P. L., and Kern, A. D. (2025). Accessible, Realistic Genome Simulation with Selection Using stdpopsim. Molecular Biology and Evolution, 42(11):msaf236.

Grossen, C., Guillaume, F., Keller, L. F., and Croll, D. (2020). Purging of highly deleterious mutations through severe bottlenecks in Alpine ibex. Nat Commun, 11:1001.

Guillaume, F. (2011). Migration-induced phenotypic divergence: the migration-selection balance of correlated traits. Evolution, 65(6):1723–1738.

Guillaume, F. and Rougemont, J. (2006). Nemo: an evolutionary and population genetics programming framework. Bioinformatics, 22(20):2556–2557.

Haller, B. C., Ralph, P. L., and Messer, P. W. (2026). SLiM 5: Eco-evolutionary Simulations Across Multiple Chromosomes and Full Genomes. Molecular Biology and Evolution, 43(1):msaf313.

Hansen, T. F. and Pelabon, C. (2021). Evolvability: A quantitative-genetics perspective. Annu Rev Ecol Evol S, 52:153–175.

Hansen, T. F. and Wagner, G. P. (2001). Modeling genetic architecture: A multilinear theory of gene interaction. Theor. Popul. Biol., 59(1):61–86.

Hernandez, R. D. (2008). A flexible forward simulator for populations subject to selection and demography. Bioinformatics, 24(23):2786–2787.

Kemppainen, P. and Guillaume, F. (2026). Linkage disequilibrium scaling improves robustness and power to detect genomic regions under selection. BioRxiv.

Lande, R. (1975). The maintenance of genetic variability by mutation in a polygenic character with linked loci. Genet Res, 26(3):221–235. Publisher: Cambridge University Press.

Lande, R. (1980). The genetic covariance between characters maintained by pleiotropic mutations. Genetics, 94(1):203–215.

Mackay, T. F. C. and Anholt, R. R. H. (2024). Pleiotropy, epistasis and the genetic architecture of quantitative traits. Nature Reviews Genetics, 25(9):639–657. Publisher: Nature Publishing Group.

Mackay, T. F. C., Stone, E. A., and Ayroles, J. F. (2009). The genetics of quantitative traits: challenges and prospects. Nat. Rev. Genet., 10(8):565–577.

Matthey-Doret, R. (2021). SimBit: A high performance, flexible and easy-to-use population genetic simulator. Molecular Ecology Resources, 21(5):1745–1754. eprint: https://onlinelibrary.wiley.com/doi/pdf/10.1111/1755-0998.13372.

Messer, P. W. (2013). SLiM: Simulating evolution with selection and linkage. Genetics, 194(4):1037–1039.

Neuenschwander, S., Hospital, F., Guillaume, F., and Goudet, J. (2008). quantinemo: an individual-based program to simulate quantitative traits with explicit genetic architecture in a dynamic metapopulation. Bioinformatics, 24(13):1552–1553.

Neuenschwander, S., Michaud, F., and Goudet, J. (2019). QuantiNemo 2: a Swiss knife to simulate complex demographic and genetic scenarios, forward and backward in time. Bioinformatics, 35(5):886–888. Publisher: Oxford Academic.

Nietlisbach, P., Keller, L. F., Camenisch, G., Guillaume, F., Arcese, P., Reid, J. M., and Postma, E. (2017). Pedigree-based inbreeding coefficient explains more variation in fitness than heterozygosity at 160 microsatellites in a wild bird population. Proc. R. Soc. B, 284(1850):20162763.

Peng, B. and Kimmel, M. (2005). simuPOP: a forward-time population genetics simulation environment. Bioinformatics, 21(18):3686 –3687.

Pickrell, J. K., Berisa, T., Liu, J. Z., Ségurel, L., Tung, J. Y., and Hinds, D. A. (2016). Detection and interpretation of shared genetic influences on 42 human traits. Nat. Genet., 48(7):709–717.

Schmid, M., Dallo, R., and Guillaume, F. (2019). Species’ range dynamics affect the evolution of spatial variation in plasticity under environmental change. Am. Nat., 193(6):798–813.

Schmid, M. and Guillaume, F. (2017). The role of phenotypic plasticity on population differentiation. Heredity, 119:214–225.

Schmid, M., Paniw, M., Postuma, M., Ozgul, A., and Guillaume, F. (2022). A tradeoff between robustness to environmental fluctuations and speed of evolution. Am. Nat., 200(1):E16–E35.

Thornton, K. R. (2014). A c++ template library for efficient forward-time population genetic simulation of large populations. Genetics, 198(1):157–166.

Turelli, M. (1984). Heritable genetic variation via mutation-selection balance: Lerch’s zeta meets the abdominal bristle. Theor. Popul. Biol., 25(2):138–193.

Walsh, B. and Lynch, M. (2018). Evolution and selection of quantitative traits. Sinauer Associates, Oxford University Press, Clarendon st., London, UK.

Wright, S. (1934). An analysis of variability in number of digits in an inbred strain of guinea pigs. Genetics, 19(6):506–536.

Yeaman, S., Gerstein, A. C., Hodgins, K. A., and Whitlock, M. C. (2018). Quantifying how constraints limit the diversity of viable routes to adaptation. PLOS Genetics, 14(10):e1007717.

Yeaman, S. and Guillaume, F. (2009). Predicting adaptation under migration load: the role of genetic skew. Evolution, 63(11):2926–2938.

